# An Extensive Atlas of Proteome and Phosphoproteome Turnover Across Mouse Tissues and Brain Regions

**DOI:** 10.1101/2024.10.15.618303

**Authors:** Wenxue Li, Abhijit Dasgupta, Ka Yang, Shisheng Wang, Nisha Hemandhar-Kumar, Jay M. Yarbro, Zhenyi Hu, Barbora Salovska, Eugenio F. Fornasiero, Junmin Peng, Yansheng Liu

## Abstract

Understanding how proteins in different mammalian tissues are regulated is central to biology. Protein abundance, turnover, and post-translational modifications like phosphorylation, are key factors that determine tissue-specific proteome properties. However, these properties are challenging to study across tissues and remain poorly understood. Here, we present *Turnover-PPT*, a comprehensive resource mapping the abundance and lifetime of 11,000 proteins and 40,000 phosphosites across eight mouse tissues and various brain regions, using advanced proteomics and stable isotope labeling. We revealed tissue-specific short- and long-lived proteins, strong correlations between interacting protein lifetimes, and distinct impacts of phosphorylation on protein turnover. Notably, we discovered that peroxisomes are regulated by protein turnover across tissues, and that phosphorylation regulates the stability of neurodegeneration-related proteins, such as Tau and α-synuclein. Thus, *Turnover-PPT* provides new fundamental insights into protein stability, tissue dynamic proteotypes, and the role of protein phosphorylation, and is accessible via an interactive web-based portal at https://yslproteomics.shinyapps.io/tissuePPT.

## INTRODUCTION

Protein turnover is a fundamental process involving the continuous synthesis and degradation of proteins in all living organisms ^1^. Protein turnover is critical for maintaining proteostasis, replacing damaged proteins, ensuring the functional integrity of tissues, and enabling a dynamic response to environmental changes ^2 3 4 5^. Maintaining proper turnover in different tissues is particularly challenging. Some proteins must be expressed in different tissues, such as key housekeeping macromolecular complexes involved in basic cellular functions, but at the same time, different tissues have dissimilar needs and must respond in different ways to maintain homeostasis. Information on protein turnover rates or lifetimes within and between tissues can help understand the principles behind tissue regulation and allow the development of targeted strategies to interfere with specific proteins and processes, opening up novel therapeutic avenues. For example, a drug targeting a protein that is rapidly synthesized and degraded in one tissue may lead to more effective treatment by minimizing the impact on other tissues where the protein’s turnover is much slower. Furthermore, protein turnover is a critical molecular regulatory layer at the post-translational level buffering, tuning, or amplifying variability in proteomic abundance. High turnover rates for proteins with many copies become energetically costly for cells ^6^ and may signal a critical role for these proteins in tissue-specific functions. Thus, a comprehensive analysis of both protein abundance (<u>PA</u>) and protein lifetime (<u>PT</u>) in mammalian tissues can enhance our understanding of cell- and tissue-specific “economic” principles and provide significant biomedical insights ^7^.

Protein turnover rates in one or multiple mouse tissues were previously analyzed in a handful of studies, most of which utilized mass spectrometry (MS)-based shotgun proteomic approaches integrated with the experimental strategy of pulsed stable isotope labeling by amino acids in cells (pSILAC) applied in animals ^8 9 7 10 11 12 13 14^. The general underlying assumption is that, once animal growth and development have ceased and a “steady state” has been reached, the rate of protein synthesis equals the rate of protein removal^15^. Thus, the metabolic labeling of proteins with ^15^N or stable heavy isotopic amino acids through dietary intake can be monitored to represent the newly synthesized protein molecules, which are then quantified by MS over time, to derive protein-specific turnover rates. Notably, due to the significant reuse of amino acids in multicellular organisms, specific mathematical modeling frameworks had to be developed to robustly determine *in vivo* protein lifetimes ^9 11 16^. Together, these pioneering studies illustrated the turnover diversity among multiple tissues and protein families, but they also suffered from a number of limitations. First, they only achieved limited proteome coverage, compared to deep analysis of protein abundance ^17 18^, often due to the limited sensitivity and throughput of the MS methods available at the time. Moreover, they examined fewer than 4-5 tissue types ^9 11 13^, and did not assess the full diversity of protein turnover among *multiple regions* of a complex organ, such as the brain. Notably, most of the previous studies have focused solely on protein turnover rates without considering the absolute and relative mRNA and protein quantitative abundances among the tissues, prohibiting a systematic understanding of protein turnover regulation.

In addition to the whole protein level, post-translational modifications (PTM) such as phosphorylation often determine protein activity ^19 20^. The abundance of tens of thousands of phosphosites (P-sites) has been profiled across various mouse tissues ^21 17^, the *in vivo* turnover of phosphorylated proteins in animals has yet to be measured. In cultured cancer cells, we and others have examined the effects of phosphorylation on protein degradation and clearance using pSILAC^15^ combined with phosphoproteomic enrichment ^22 23 24 25^. Nevertheless, how site-specific phosphorylation regulates *in vivo* protein lifetime and stability across different tissues and tissue regions remains fully unexplored. Such knowledge is critical as it may reveal new P-site nodes that can be targeted in human diseases such as neurodegeneration.

Here, we harness advanced quantitative proteomic strategies, namely data-independent acquisition (DIA) ^26 27 28^ and tandem mass tagging (TMTpro) ^29 30^ to quantify proteome and phosphoproteome turnover across multiple samples and labeling points, overcoming the challenges of irreproducibility and inconsistency that have limited previous studies. We extensively map protein and P-site turnover behaviors across eight tissues and nine brain regions in mice. Our datasets feature high coverage and increase by *three-fold* the number of known protein lifetimes *in vivo*. The resulting atlas, called *Tissue-PPT*, is a comprehensive resource that provides in-depth information on both <u>PA</u> and <u>PT</u> in a mammalian tissue-specific context. A key strength of our study is the well-matched nature of the datasets, where protein abundance and turnover, unmodified proteins and phosphorylated proteins, and tissue-specific profiles are closely aligned across multi-omics layers. This precise matching enables more accurate comparisons and deeper insights into the regulation of proteostasis. Our findings reveal that proteostasis networks can be widely rewired by protein-protein interactions (PPI), organellar localizations, and site-specific phosphorylation such as critical P-sites on Tau and α-synuclein in the brain. Our *Resource* of <u>tissue</u> <u>p</u>roteome and <u>p</u>hosphoproteome turnover atlas, or *Tissue-PPT,* is easily accessible online via an interactive web portal (https://yslproteomics.shinyapps.io/tissuePPT).

## RESULTS

### In-depth quantitative protein turnover landscape of mouse tissues and brain regions

We profiled the proteome-wide protein turnover kinetics in terms of protein half-lives (i.e., T_50_; hereafter, protein lifetime or <u>PT</u> in short) of the heart, liver, spleen, lung, kidney, gut, plasma, and nine brain regions, including the cerebellum, frontal cortex, substantia nigra, thalamus, amygdala, entorhinal cortex, hippocampus, and olfactory bulb (**Figure 1**). The PTs were measured *in vivo* using pSILAC labeling of Lysine-6 containing food fed to mice ^31^ over periods of 8 and 32 days. This approach was firstly validated using five biological replicates of whole brain tissues and four labeling time points (**Figure S1A-B**) and then applied for multi-tissue proteomic measurements employing both DIA-MS ^26 27 28^ and TMTpro 16-plex labeling ^29 30^ (**Methods**). To enhance the precision of protein turnover quantification, we implemented the BoxCarmax-DIA multiplexing schema ^32^ for DIA-MS and performed extensive peptide-level fractionation (>80 fractions) for TMT ^33 34^ to effectively reduce the MS/MS data complexity for determining heavy-to-light (H/L) ratios (**Figure 1A**). An ordinary differential equation-based computational framework was subsequently employed to model the amino acid recycling and fit the Lysine-6 kinetics for DIA-MS and TMT datasets ^16^. DIA-MS and TMT generated highly consistent and reproducible lifetimes across proteins (Spearman *rho* =0.88, **Figure 1B, Figure S1C**) and across tissues (*rho*=0.96, **Figure 1C**).

**Figure 1.**
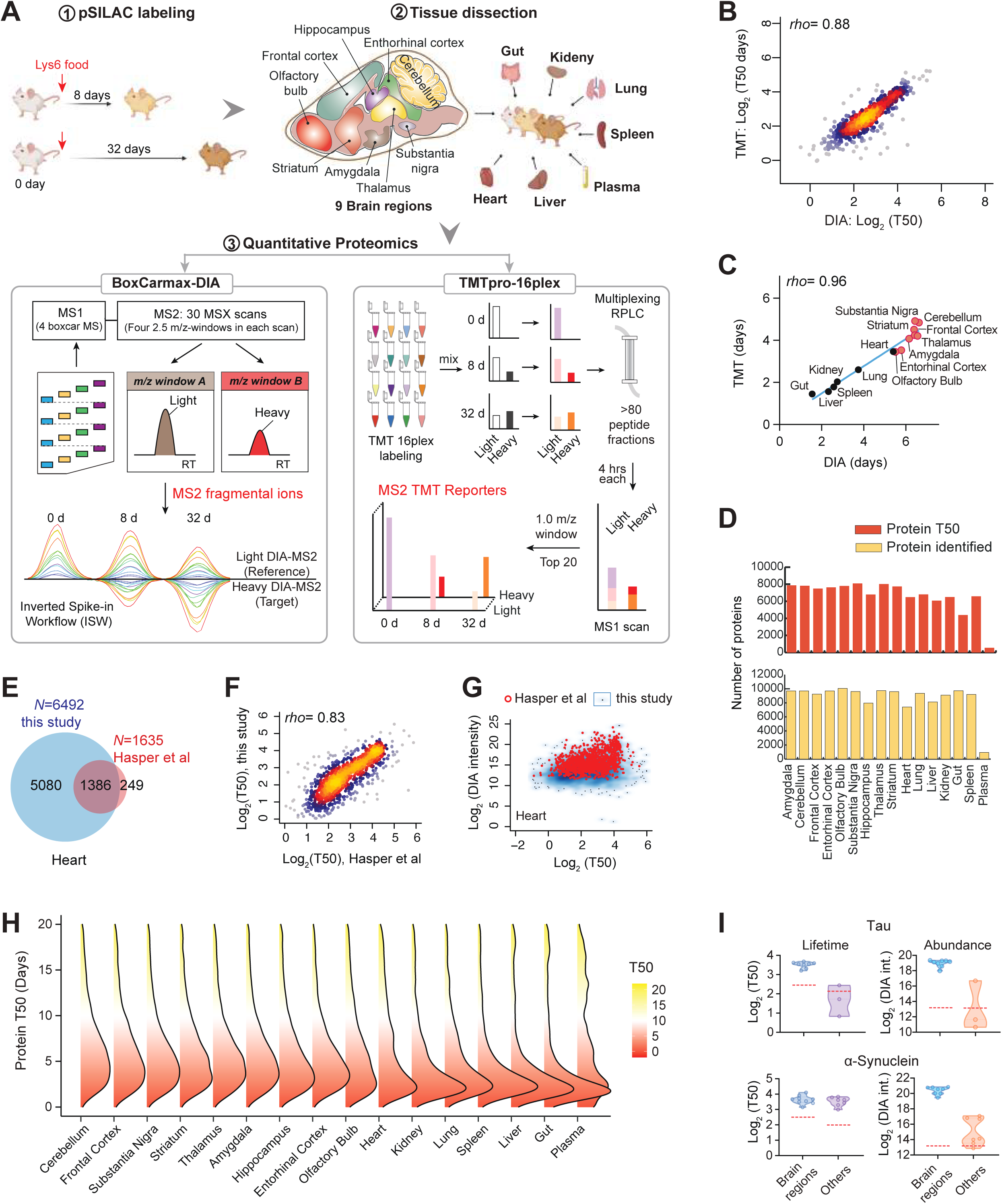
Generation of a high-quality protein turnover atlas across mouse tissues and brain regions. (A) pSILAC-MS workflow used for cross-tissue protein turnover analysis in mice. The BoxCarmax-DIA and TMTpro methods were employed to improve quantification accuracy. (B) Spearman correlation of protein lifetimes across the proteome, as quantified by DIA and TMT methods. (C) Spearman correlation of protein lifetimes within each tissue, as quantified by DIA and TMT methods. (D) Summary of proteome coverage for both protein identification and protein lifetime profiling. (E) Venn diagram comparing mouse heart proteome coverage between this study and Hasper et al. (F) Spearman correlation of PT results (i.e., T_50_ values) between the two studies for the heart proteome. (G) Scatterplot displaying the comparison of protein abundance and lifetime between this study and Hasper et al. (red dots). (H) Density plot of protein lifetimes across 16 mouse tissue samples. (I) Violin plots summarizing the protein abundances and lifetimes of Tau and alpha-synuclein. The red dashed lines indicate the proteome-wide averaged levels of abundances and lifetimes.

By integrating all measurements (**Methods**), we quantified lifetimes for 11,171 unique protein groups across various tissues and brain regions. On average, 9275 proteins were detected and quantified in non-plasma tissues, and turnover rates were measured for 7075 proteins per tissue from Lysine-containing peptides from the same datasets (**Figure 1D)**. We developed the *Tissue-PPT*, a Web-page App to support both individual protein- and protein list-level exploration of this extensive dataset. The analytical depth of *Tissue-PPT* is comparable to recent large-scale proteome mapping efforts of mouse tissues ^17 18^, but it effectively *triples* the number of protein lifetimes measured across multiple tissues ^11 13 14^. For instance, using Lysine-8 SILAC labeling, Rolfs et al.^11^ measured protein turnover in five mouse tissue types, including liver (n =2004 proteins) and plasma (n =155). In comparison, *Tissue-PPT* measured lifetimes for 6077 liver proteins and 516 plasma proteins. Similarly, Hasper et al. ^13^, utilizing ^15^N labeling, analyzed four tissues, including liver (n =2099) and heart (n =1635). In contrast, *Tissue-PPT* determined turnover rates for 6492 proteins in heart. A closer comparison shows that *Tissue-PPT* covers 84.6% of heart proteins measured by Hasper et al. and reports highly correlated protein lifetimes (*rho* =0.83, **Figure 1E-F**). Reassuringly, based on protein DIA-MS signals, *Tissue-PPT* achieved a much deeper heart proteome, because it effectively measured turnover rates for more low-abundance proteins (*P* = 7.3e-137, Wilcoxon test, **Figure 1G**). A similar observation was made when comparing the liver proteome results (P =1.01e-150, **Figure S1D-G**), confirming the substantial analytical depth of *Tissue-PPT*.

The overall proteome turnover patterns across tissues showed moderate similarity, with most of the proteome (66.7%-80.04%) having a PT of less than 10 days, indicative of the basic metabolism dynamics in mice (**Figure 1H**). The median PT ranged from 3.27 days in the gut to 6.45 days in the cerebellum. The nine brain regions overall displayed significantly higher median PT (5.89 ± 0.42 days) than other tissues (3.27 days for the gut, 3.28 days for the liver, 3.62 days for the spleen, 3.82 days for the kidney, and 4.21 days for the lung), except for the heart (5.61 days). The longer protein lifetime in brain regions is consistent with previous reports ^8 14^ and can be mainly attributed to the brain’s lower regenerative capacity. Similarly, the heart has a very limited regenerative capacity, and cardiomyocytes rarely divide after birth, contributing to its slow proteome turnover ^35^. Interestingly, the overall differences in protein turnover between tissues cannot be solely explained by cell proliferation. For example, despite the cell doubling time in the liver was determined to be 51 days ^13^, its overall proteome lifetime is short, indicating additional *in vivo* factors significantly influencing protein turnover. More importantly, protein-specific turnover demonstrates considerable diversity: while the first 1% percentile have a PT of less than 1 day, the 99% percentile and above have a PT of greater than 100 days on average in all tissues. The mostly short-lived and long-lived proteins are different between tissues, enriching a variety of functions (**Figure S2A**). Only 49 proteins are consistently the top 5% most long-lived in each of the nine brain regions (**Figure S2B**), comprising those proteins enriched in TCA cycle and respiratory electron transport (*P* =3.91E-12), myelin sheath (*P* =2.43E-11), chromatin assembly (*P* =8.59E-10), and collagen-containing extracellular matrix (*P* =4.96E-13). Among them, only six structural proteins are commonly top 5% long-lived across all the other tissues, including Plp1 (PT, 87.54 days on average), Cldn11 (153.42 days), Mog (107.53 days), Nfasc (49.16 days), Col5a2 (114.16 days), and Ccdc177 (50.31 days). Together, these results underscore the importance of measuring and understanding individual protein turnover rates in various tissues.

### Protein abundance and lifetime profiling jointly define tissue proteome function and stability

Protein turnover depicts a functional dimension that is largely independent of protein expression ^6 32^. In this regard, our *Tissue-PPT* integrates <u>matched PA and PT</u> of the same proteins, which may enhance our understanding of protein essentiality in tissues. For example, by examining proteins Tau (MAPT) and α-synuclein (SNCA), we confirmed that both proteins exhibit significantly higher abundance in brain regions compared to other tissues. Furthermore, while α-synuclein shows comparable lifetimes in the brain and other tissues, the lifetime of Tau is significantly longer in the brain (**Figure 1I**), where it would be prone to accumulate changes that can lead to pathologies such as tauopathies ^36 37^.

To explore biological insights from PA and PT quantification, we firstly examined the total variance of PA and PT datasets using principal component analysis (PCA). As expected, all brain regions formed a distinct cluster separated from other tissues in the PCA of both PA and PT (**Figure 2A**), indicating a smaller biological variability between brain regions. The PT for the gut appeared as an outlier among solid tissues, possibly owing to its rapid cell proliferation. Overall, the cross-tissue correlation of PT was lower than that of PA (**Figure 2B**), suggesting substantial protein turnover control in different tissue contexts. Following, we annotated the functions of the 5% shortest- and longest-lived proteins based on their averaged PTs in brain regions and non-brain tissues (**Figure S2C-E**). Intriguingly, distinct biological processes enriching either short- or long-lived proteins were identified (**Figure 2C and Figure S2F**). Just as examples, the “core matrisome” and “aerobic respiration” are associated with long-lived proteins, whereas “DNA damage response” and “protein polyubiquitination” are linked to rapid turnover, all consistent with previous reports ^38 39 40 41 42^. To further analyze how PT’s dependency on PA affects functional analysis, we profiled the correlation between PA and PT across all tissues for different protein functional classes (**Figure 2D**). This analysis reveals that certain protein groups, such as the chaperonin complex, respiratory chain complex I, and proteins involved in organ formation, exhibited a positive correlation between PA and PT, indicating a coordination between protein expression and lifetime. In contrast, proteins from the ribosome and the 48S preinitiation complex show no or even negative PA-PT correlations, suggesting that cells use protein turnover to regulate the protein translation machinery, thereby promoting tissue homeostasis.

**Figure 2.**
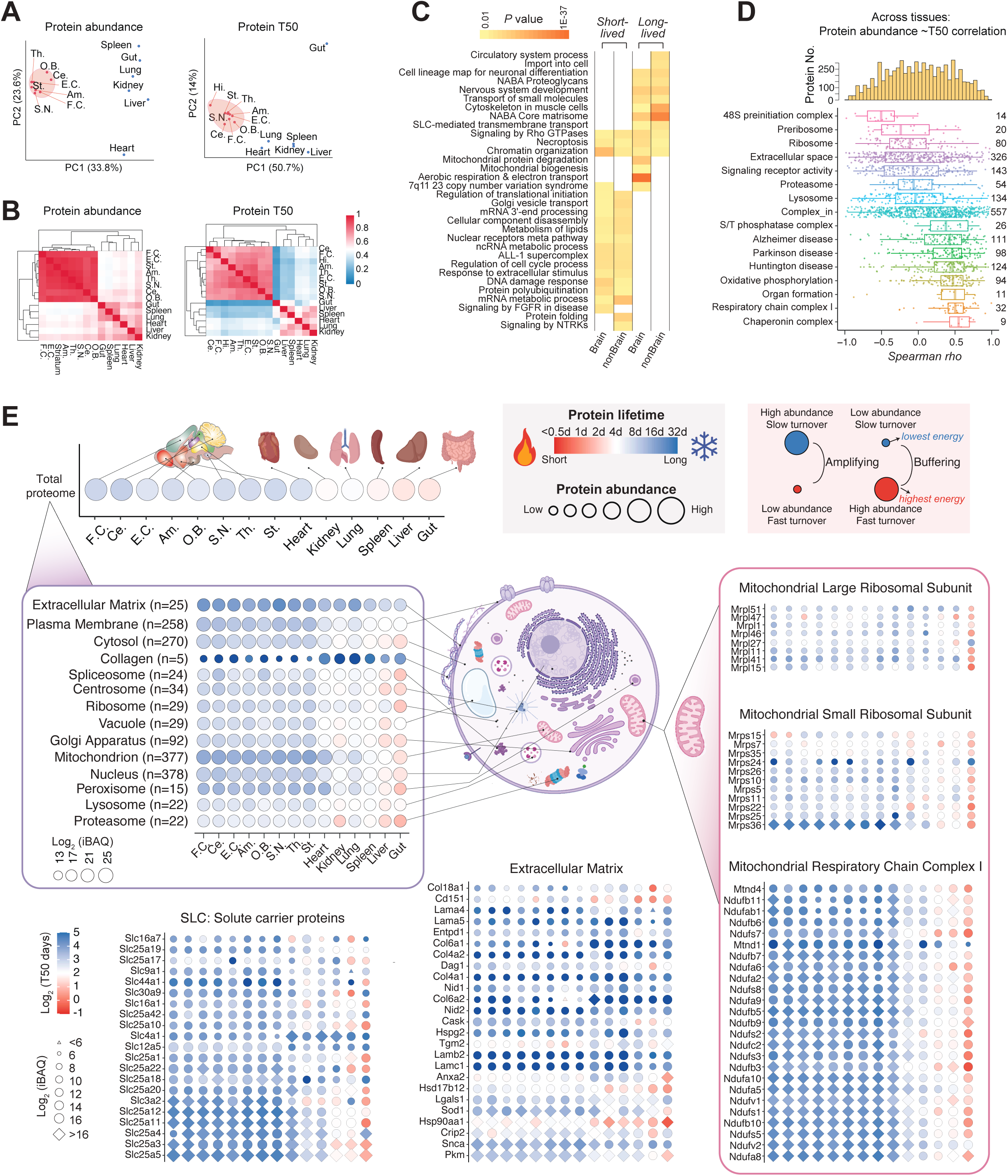
Concurrent protein abundance (PA) and lifetime (PT) profiling of the mouse tissue proteome. (A) Principal Component Analysis (PCA) of PA and PT in brain regions and solid tissues. (B) Hierarchical clustering heatmap showing Pearson correlation between tissues and brain regions based on PA and PT, respectively. (C) Selected biological processes enriched among the 5% shortest- and longest-lived proteins, based on their average PTs in brain regions and non-brain tissues. Enrichment p-values were reported by Metascape. (D) Distribution of cross-tissue Spearman correlation between PA and PT for all proteins across tissues and regions. Upper panel: Density histogram of Spearman rho values for all proteins. Lower panel: Boxplots of protein-specific Spearman rho values for selected GO terms. (E) Heat-circle (HC) plot visualization of PA and PT across tissues at different levels of cellular organization. Upper left panel: PT across all tissues, with proteome abundance normalized. Middle left panel: HC plot visualizing main cellular components across samples. Other panels: HC plot examples for individual proteins within selected protein groups. The blue-to-red color gradient denotes protein lifetime from long to short. The size of the HC plot circles is proportional to the Log_2_(iBAQ) value indicating PA. Triangle: iBAQ value is in the bottom 5% (i.e., Log_2_(iBAQ) < 6). Diamond: iBAQ value is in the top 5% (i.e., Log2(iBAQ) > 16).

Next, to illustrate the synergistic profiling of PA and PT, we developed a heat-circle (HC) plot. In this synchronized plot, protein iBAQ values (a proxy for protein copy number) ^43 44^ are derived from DIA-MS readouts (**Methods**), determining the relative size of the circles. The color gradient from red to blue indicates the lifetime of the proteins or protein sets, with red representing short-lived proteins and blue representing long-lived ones (**Figure 2E**). The HC plot essentially provides a comprehensive view of proteome stability and cellular energy expenditure across various *levels*, including e.g., tissues, functional protein groups, organelles, sub-organelles, and individual proteins. *At the organelle level*, the HC plot reveals that the extracellular matrix (EM) consistently enriches long-lived proteins across tissues, indicating that EM proteins do not undergo rapid turnover compared to other cellular components, consistently with previous reports ^9 38^. A similar pattern is observed for components of the plasma membrane, such as solute carrier (SLC) proteins, which are likely critical for maintaining tissue integrity. Collagen proteins, although few in number, are highly abundant in non-brain tissues and are extremely long-lived in all tissues, reflecting their critical roles in maintaining tissue structure ^45 46^. In contrast, mitochondrial and nuclear proteins exhibit higher cross-tissue variability, indicating tissue-specific dynamic regulation. *At the sub-organelle level*, the HC plot shows that respiratory chain complex I proteins in the brain have higher abundances and longer lifetimes compared to other tissues and other mitochondrial proteins, such as mitochondrial ribosomal subunits, emphasizing the importance of oxidative phosphorylation in the brain ^12^, consistent to **Figure 2C** results. *At the individual protein level*, HC plot indicates that SLCs exhibit a wide range of PAs but have similar PTs across tissues. Interestingly, certain proteins, such as Mrps24 in the mitochondrial ribosome, MTND1 in respiratory chain complex I, HSP90aa1 and Hsd17b12 in the EM, and Slc4a1 and SLC12a5 in the SLC, show exceptional lifetimes, potentially pointing to *moonlighting* protein functions independent of their complexes and functional classes. Together, the HC plot, which is fully supported in our *Tissue-PPT* App (**Figure S3A-B**), effectively visualizes both PA and PT, offering complementary insights into tissue functional diversity.

### Lifetime diversity of E3 ligases is critical for determining tissue-specific proteolysis

To understand the proteolysis landscape across different tissues, we examined PA and PT profiles for the major cellular protein degradation machineries ^47^: the ubiquitin (Ub)-proteasome system (UPS), lysosome, E3 Ubiquitin Ligases (E3), E3 accessory proteins, deubiquitinating enzymes (DUBs), and protein folding chaperones (**Figure 3**). Using HC plots, we found that both 19S and 20S proteasomes are tightly regulated by similar PA between subunits and correlated PT across tissues (**Figure 3A**). The kidney has an exceptionally fast turnover of the proteasome compared to the total proteome turnover (**Figure 2E**), which might be crucial for the kidney’s function in degrading and reabsorbing the high load of proteins and small peptides filtered by the glomerulus. Compared to other protein degradation components, lysosomal proteins display a much higher PA dynamic range and variability (**Figure 3B** and **3D**). Individual enzymatic proteins in the lysosome, such as Man2b1, Pla2g15, and Capn1, are particularly short-lived in brain regions. Conversely, lysosomal proteins are relatively long-lived in the spleen (**Figure 3C**), likely due to the spleen’s role in immune surveillance and phagocytosis ^48^. Notably, among all protein degradation machineries, DUBs maintain the most stable PA levels across tissues (**Figure 3D** and **Figure S4A-C**), suggesting the core activities mediated by DUBs are fundamental and universally required across all cell types. Despite general high PA levels, specific molecular chaperones, like heat shock proteins (HSPs), exhibit diverse PT profiles (**Figure S4A**). For example, Hspa12a is long-lived not only in the brain but also in other tissues, potentially due to its localization in extracellular exosomes. Again, Hsp90aa1, the stress-inducible isoform of the cytosolic chaperone protein HSP90 ^49^, has a short lifespan across tissues including brain, possibly reflecting its role in rapid proteostatic responses. Interestingly, E3 ligases and their accessory proteins, despite being the least abundant of the protein degradation machineries, are the most dynamic, exhibiting the highest PT variability across tissues (**Figure 3E**). Together, these findings underscore that the key steps of protein degradation can vary in different tissues and highlight the critical role of E3 ligase pathway turnover in maintaining tissue-specific proteostasis.

**Figure 3.**
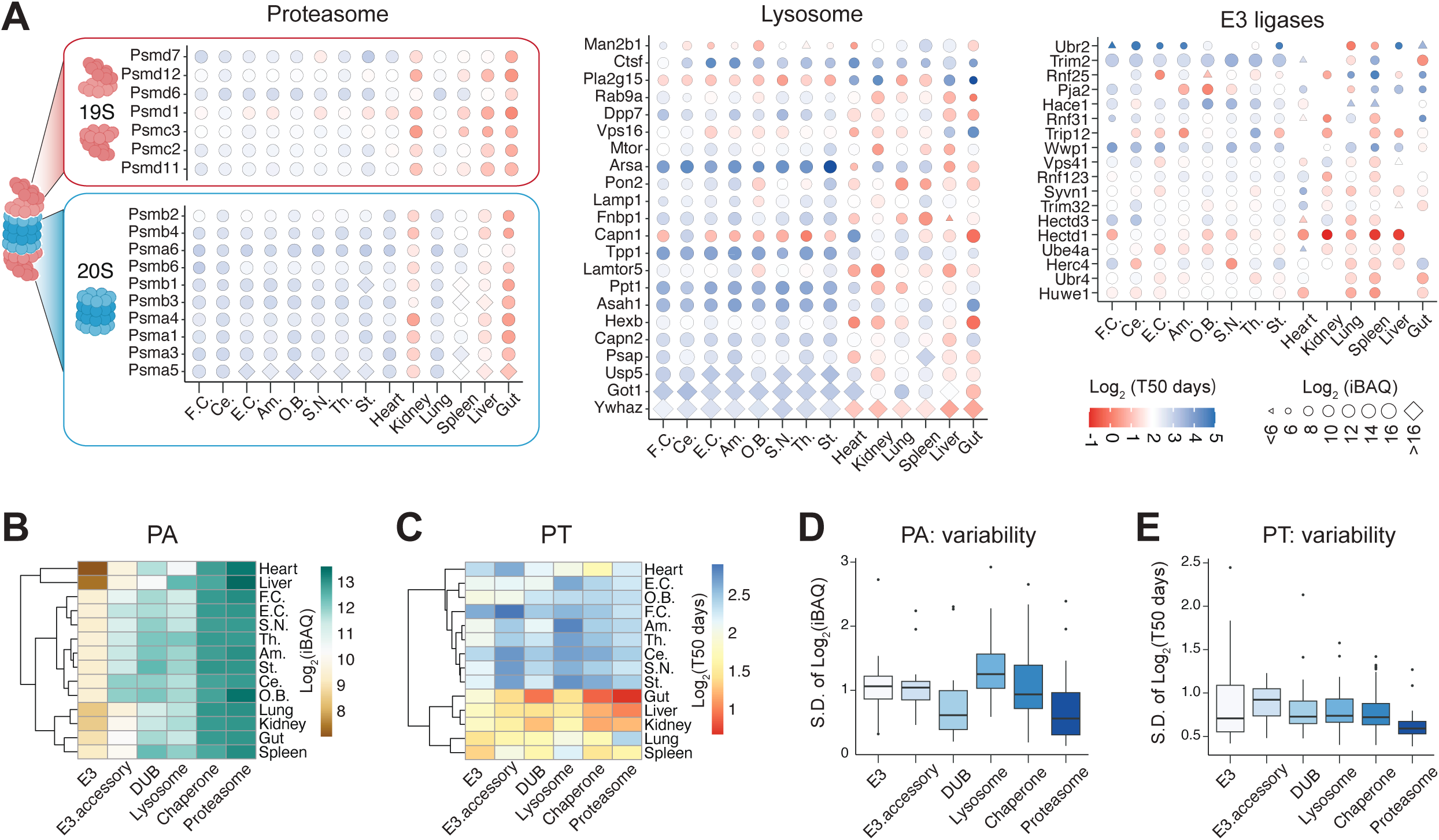
Characterization of protein removal processes across tissues. (A) Heat-circle (HC) plot of proteasome subunits (19S and 20S), lysosomal proteins across tissues, ad E3 ubiquitin ligases (E3s) across tissues. The color and size are defined as in Figure 2E. Those E3 proteins with PA quantified in less than 12 tissue samples were filtered. (B) Hierarchical clustering heatmap of PA profiles of the five proteins representing protein degradation machineries. The brown-to-green color bar indicates the increasing relative abundance in terms of Log_2_ (iBAQ values). (C) The same heatmap as (D) for PT profiles. The red-to-blue color bar indicates the increasing relative lifetime in terms of Log_2_ (T50 days). (D) The boxplot of standard deviation S.D. of [Log_2_ (PA of each protein)-Log_2_(PA of averaged level)] for each protein list indicating the PA variability across tissues. (E) The boxplot of standard deviation S.D. of [Log_2_ (T_50_ of each protein)-Log_2_(T_50_ of averaged level)] for each protein list indicating the PT variability across tissues.

Furthermore, we investigated the utilization of the ubiquitin resource across tissues. By re-searching our DIA-MS data for Ub modifications, we assessed signature peptides that distinguish Ub chains linked to different lysine (K) residues ^50 51 47^. including K48, K63, K11, K27, and K6, which were detected in most tissues ^52 53^ (**Figure S4D**). Label-free quantification (**Methods** ^47^) revealed that K48-linked Ub, which plays a classic role in the UPS, is the predominant Ub chain. In line with the slow turnover in brain regions, we observed the lowest levels of K48-linked Ub in brain regions. Beyond relative abundance, our analysis additionally assessed the diversity of recycling Ub resources. Strikingly, K48-linked Ub, once synthesized, appears to be retained significantly longer than K63-linked Ub and other proteins across most tissues (**Figure S4E**). This finding might indicate the complex coupling mechanism between substrate deubiquitination of K48-linked Ub and degradation ^54^ and differential recycling strategy of variable polyubiquitins ^55^. Additionally, the liver demonstrated an exceptionally high turnover of K63-linked Ub which has the diverse signaling roles such as endocytosis and autophagy ^56^. Collectively, our findings highlight tissue specific Ub distribution, architectures ^57^, and dynamics.

### Protein lifetime tightly correlates with protein-protein interaction in tissues

Proteins rarely act alone in a living cell. When two proteins are engaged in a physical protein-protein interaction (PPI), it is tempting to hypothesize that they are synthesized and degraded in a coordinated manner. Previous studies have demonstrated that PTs of the same organelle, family, or complex are often correlated ^58 59 9^. Recently, Skinnider et al. established a comprehensive PPI dataset across various mouse tissues, using protein correlation profiling mass spectrometry (PCP-MS) ^60^. By integrating that dataset with our *Tissue-PPT*, we were able to correlate PT and PPI across tissues in detail and obtain new insights. First, we confirmed that the *binary correlations* of PA between PPI partners based on CORUM ^61^, BioPlex ^62^, and Skinnider et al. ^60^ are all significantly higher than those between random protein pairs that do not interact (**Figure 4A**), consistent with the established proteome organization mechanisms through PPIs ^6 63^. Remarkably, PT profiles across tissues showed a similar correlating pattern (**Figure 4B**): e.g., Skinnider et al. identified 107,553 PPIs overlapping with our PT data, which are 2.69 and 14.76 times more than PPIs extracted from CORUM and BioPlex; and this PT data demonstrates the most dramatic aforementioned correlation difference (mean *Pearson* correlation, 0.53 for PPI partners vs. 0.07 for non-PPI pairs). The even more pronounced correlation of PTs between PPI partners, compared to PAs (*right* panels of **Figure 4A-B**), demonstrates the importance of resolving tissue-specific PPI networks ^60 64^, underscoring an previously underappreciated dependency of PT on PPI. We additionally mapped the *Pearson* correlations based on PA and PT across tissues to PPIs confidence levels in hu.MAP ^65^. Consistently, this analysis reveals that PTs provided significantly better discrimination than PAs for those most confident PPIs (Level 5, Extremely High confidence, **Figure 4C**).

**Figure 4.**
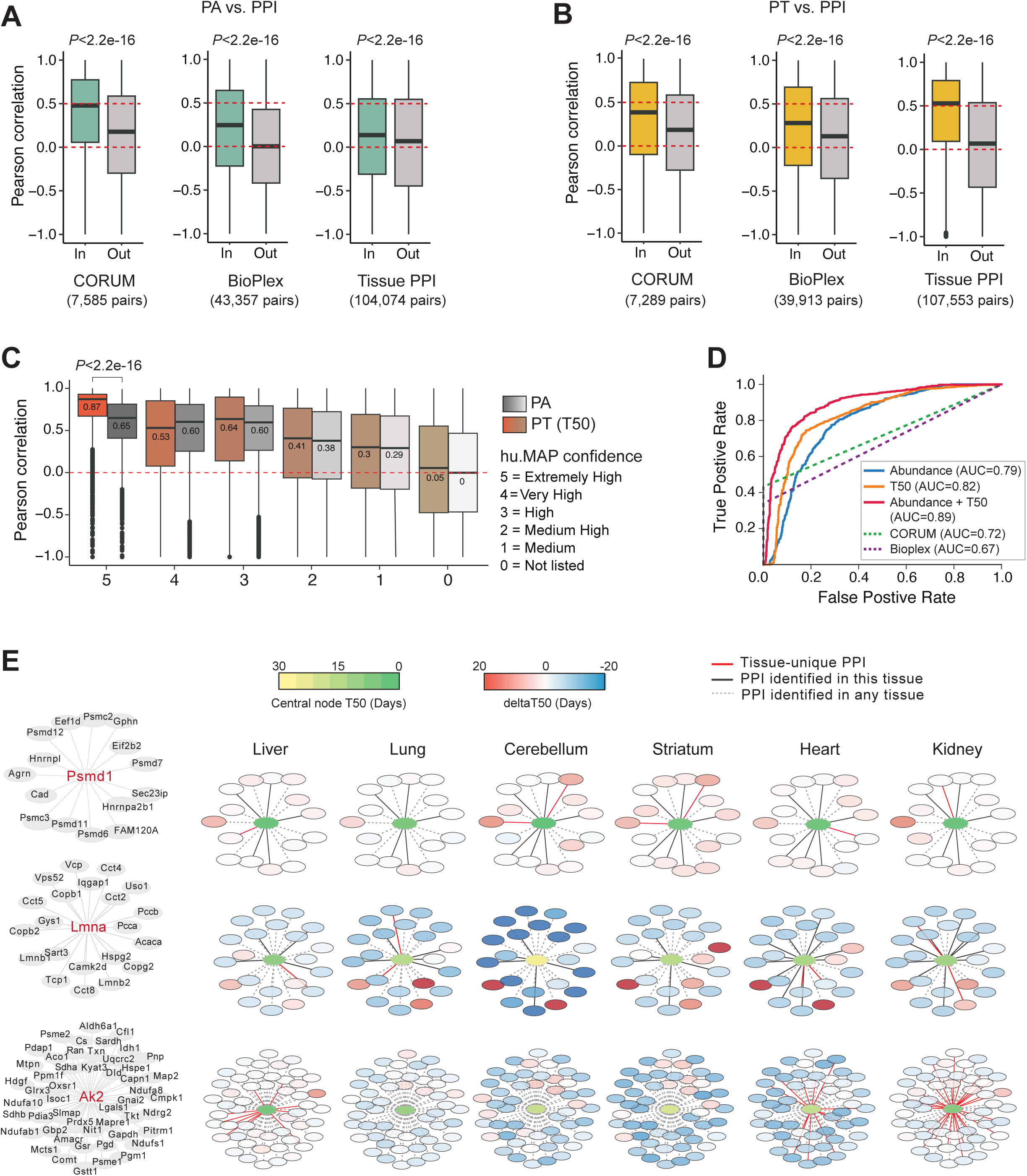
Strong association between protein lifetime and protein-protein interaction (PPI) across tissues. (A) Boxplots of correlation coefficients for PA between PPI partners, based on CORUM, Bioplex 3.0, and PCP-derived mouse tissue-specific PPI lists (Skinnider et al.). P-values were calculated using the Wilcoxon test. “In” and “Out” denote PPIs included or not described in these resources. (B) The same boxplot as in (A) for PT. (C) The same boxplot as in (A) and (B) for both PA and PT, based on PPI confidence levels retrieved from the hu.MAP database. Levels 1-5 indicate increasing PPI confidence in hu.MAP. (D) Receiver operating characteristic (ROC) curves indicating the predictive power of PA, PT, and their combined panel using logistic regression, alongside CORUM- and Bioplex-derived lists. The Extremely High and Very High confidence groups of PPIs from hu.MAP were used as true positives (TP). An equal number of randomly generated false pairs were used as false positives (FP) (Methods). AUC, Area Under the Curve. (E) Visualization of PT for PPI partners of PSMD1, LMNA, and AK2 proteins (the central nodes) in selected tissues. PT values of the central nodes are visualized using a yellow-to-green color gradient. The red-to-blue color bar denotes the relative PT difference between PPI partners and central nodes (i.e., T_50_ difference). The red, black, and dashed black edges represent PPIs unique to the specific tissue, all PPIs in the specific tissue (not necessarily unique), and PPIs in any of the mouse tissues (the whole dataset), respectively, according to Skinnider et al.

Conversely, we asked whether PT correlation between protein pairs would suffice to predict a PPI, by using the receiver operating characteristic (ROC) analysis, which essentially assesses the accuracy of model predictions. We herein firstly assigned Level 4 (High) or 5 (Extremely High) PPIs in hu.MAP as true positives. We found that PT alone could effectively predict PPIs (AUC =0.82, **Figure 4D**). Notably, combining PT and PA yielded an AUC of 0.89. And both PA and PT covariations outperformed CORUM and BioPlex in predicting hu.MAP PPIs, highlighting the potential of leveraging PT and PA across tissues to predict PPIs or refine PPI lists in animals. Intriguingly, examining PT differences between a given protein and its PPI partners in one specific tissue or in any of the measured tissues identifies PPI partners with constantly deviated turnover rates (**Figure 4E and Figure S3A**), such as Agrn for Psmd1, Lmnb2 for Lmna, and Map2 for Ak2 (in liver and kidney). These results might indicate the presence of additional partners and roles for these proteins which show very peculiar “PT-deviating” protein turnover profiles.

Taken together, our findings demonstrate that cross-tissue PT is tightly constrained by PPIs, offering new insights into turnover dynamics of individual proteins and protein networks.

### Cross-tissue multi-omic analysis reveals that peroxisome is particularly regulated through protein turnover

How does our proteome turnover data, representing the post-translational layer of regulation, contribute to understanding the regulatory principle at the basis of tissue diversity? To address this question, we integrated *Tissue-PPT* with a recent dataset ^66^ describing both the transcriptome and translatome – measured using RNA sequencing and ribosome profiling, respectively – from six mouse tissues, five of which overlap with the ones in *Tissue-PPT* (liver, heart, lung, kidney, and brain).

***First*,** we performed multi-layered absolute correlation analysis across all proteins per tissue. The highest correlation is observed between mRNA and the translatome (Spearman *rho*=0.76-0.83), followed by the correlation between the translatome and the proteome (*rho*=0.41-0.54) and the correlation between mRNA and the proteome (*rho* =0.42-0.55) **(Figure 5A**). Additionally, absolute PT exhibits weak or no correlation with absolute PA, but a slight yet notable negative correlation with levels of both mRNA and RNA being translated. This result aligns with the reported buffering role of protein turnover in globally modulating transcriptional ^67^ and translational regulations to fine-tune the functional proteome across all tissues ^68^. ***Second,*** to discern which layer drives the specific tissue proteotype ^69^, we contrasted multi-layered correlations across tissues, between proteins identified in all tissues (n =1919, **Figure 5B**) and proteins exhibiting >4-fold higher PA levels in a particular tissue (n =418). We found that tissue-enrich proteins ^70^ are predominantly driven by regulatory steps *prior to* protein synthesis rather than by protein turnover, because of the much stronger correlations between mRNA, translatome, and proteome than the correlation between mRNA/translatome and PT. ***Third,*** to determine whether certain proteins and functions are particularly regulated by protein turnover, we combined protein-specific correlations based on the protein’s GO cellular compartment (**Figure 5C**). This analysis revealed a fascinating anti-correlation between mRNA and PT, as well as between translatome and PT (*rho* = −0.42 and −0.41, respectively), for peroxisomal proteins, demonstrating that peroxisomes are exceptionally and primarily controlled by protein degradation, a novel insight not previously reported. We then compared the averaged levels of mRNA, mRNAs being translated, PA, and PT in each organelle (**Figure 5D**) and for individual peroxisome proteins among tissues (**Figure 5E-F and Figure S5A-B**), verifying this exceptional pattern of peroxisome proteome. We found that, in general, 60% of peroxisomal proteins, despite having low transcript and translation levels, are long-lived, whereas the remaining 40% exhibit the opposite characteristics (i.e., high levels of transcript/translation but short-lived).

**Figure 5.**
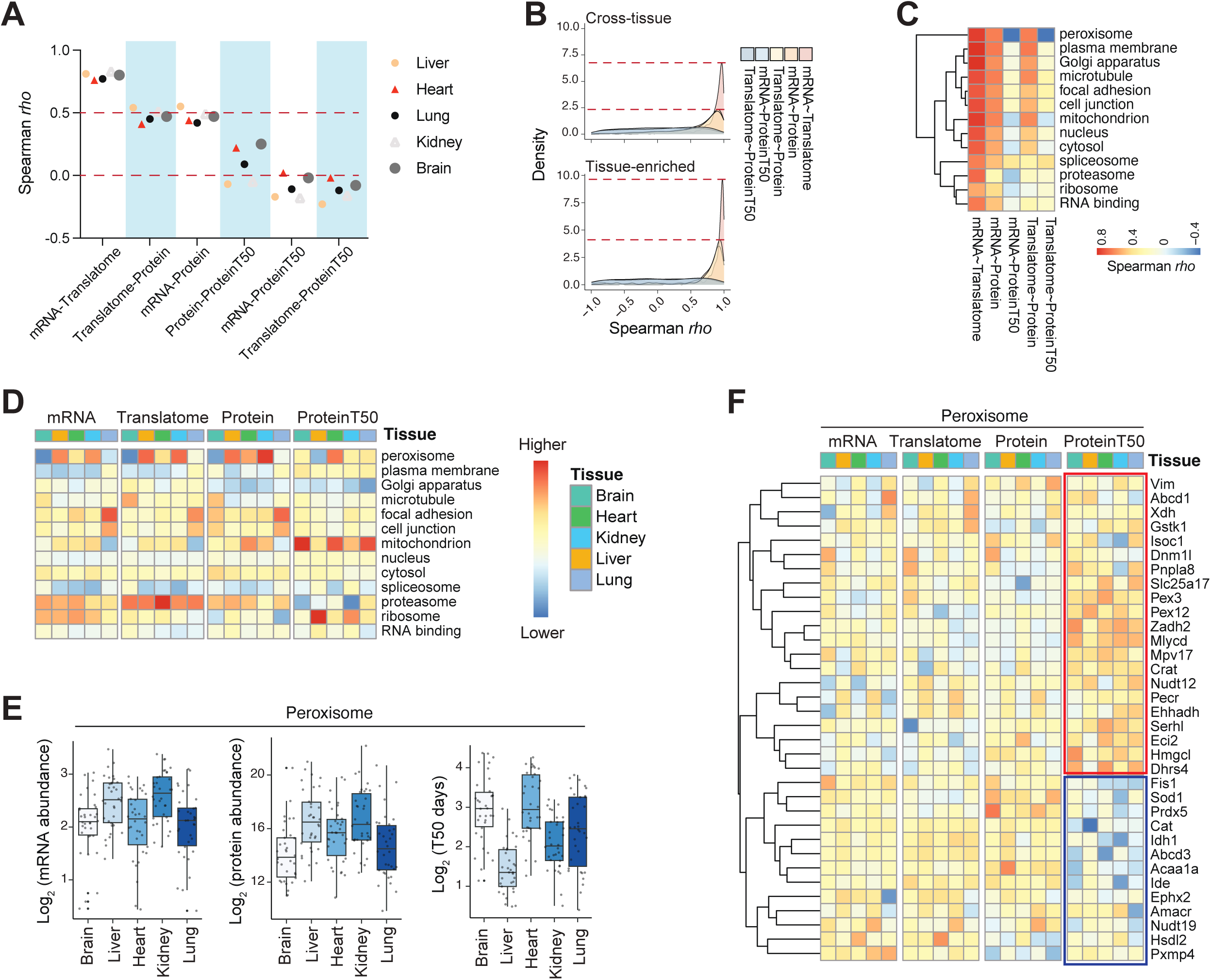
Cross-tissue multi-omic analysis and turnover control of peroxisome proteins. (A) Proteome-wide absolute Spearman correlation between measurements of mRNA, translatome, PA, and PT across five tissues. The brain results were determined by averaging all brain regions. (B) Density plots of protein-specific Spearman correlation rho values between multi-omic layers for all measured proteins (upper panel) and tissue-enriched proteins (lower panel). Tissue-enriched proteins are defined as those with protein abundance at least four times higher than the average of other tissues. (C) Heatmap visualizing the cross-tissue Spearman correlation between multi-omic layers. (D) Heatmap of quantitative results (column-scaled) for GO Cellular Components across multi-omic layers. The blue-to-red color bar represents the summed values of proteins associated with specific GO Cellular Components. (E) Boxplots of mRNA abundance, PA, and PT levels for peroxisome proteins. (F) Heatmap of quantitative results for individual peroxisome proteins measured across five tissues and multi-omic layers.

Previous studies reported that peroxisomes are regulated by a selective autophagic degradation process called pexophagy^71^ and that peroxisomal biochemical pathways are specialized in different organs ^72^. We thus checked PA and PT cross-tissue profiles for proteins involved in pexophagy in our data. Indeed, we observed a significant correlation between the abundance of pexophagy-associated proteins (n =27) and the overall lysosome levels (R =0.54, **Figure S5C**). Both HC plots and individual PA∼PT correlation plots, generated using *Tissue-PPT*, revealed negative PA vs. PT correlation for several pexophagy-related proteins (**Figure S5D-F**) including Pex3, an essential activator of pexophagy ^73^ and Atg12, a ubiquitin-like protein critical for the formation of autophagosome and autophagy ^74 75^. Together, the particularly enhanced control of peroxisomal proteins via turnover might play a crucial role in enabling cells to rapidly adapt to cellular stress and metabolic demands.

### A bimodal distribution of plasma protein abundances and lifetimes highlight the tissue origins

The plasma proteome contains proteins released from various tissues into the bloodstream. Our results uniquely allow us to address whether tissue proteins maintain their PTs in plasma. Previously, Niu et al. analyzed the levels of 420 proteins commonly detected in the liver and plasma in patients with alcohol-related liver disease (ALD) ^76^. They reported two groups of plasma proteins: a “diagonal cluster” showing largely correlated PAs between liver and plasma, and a “vertical cluster,” speculated to reflect tissue leakage, which exhibited no correlation ^76^. Strikingly, using our PA data, we confirmed both “diagonal” and “vertical” clusters, not only in the liver but also in most other tissues (**Figure S6A**), extending the previous observation. Proteins in the “diagonal cluster” were much fewer when compared to brain regions, likely due to the existence of blood-brain barrier (BBB). We further found that PTs in plasma are generally longer than in tissues and brain regions, possibly due to the lack of UPS and cell proliferation in blood plasma (**Figure S6B**). Despite this, our results show that proteins in the “diagonal cluster” tend to maintain their lifetimes in both tissue and plasma samples (**Figure S6C**). We speculate that this can be explained by the fact that these tissues are rich in blood capillaries. To summarize, the PT profiles of the plasma proteome support the existence of two *origination* groups of proteins in human plasma.

### Site-specific phosphorylation functionally shapes protein lifetime across tissues

In addition to the comprehensive analysis of whole protein turnover, *Tissue-PPT* presents the first tissue phosphoproteome turnover dataset. Overall, our DIA-MS and pSILAC-DIA measured the abundances of 67,169 P-sites and T_50_ of 40,573 P-sites, delineating the dynamics of *in vivo* phosphorylation on a large scale. In the brain regions and non-brain tissues, respectively, we quantified 34,157 ± 2207 and 22,821 ± 4650 P-site-carrying peptides in terms of abundance and 12,861 ± 1441 and 7575 ± 2681 of them in terms of T_50_ (**Figure 6A**, and **Figure S7A**). In contrast, only about 100 P-sites were quantified in the blood plasma. The PCA plots of phosphoproteomic abundance and turnover variances displayed the inter-tissue and inter-region patterns similar to total proteomic results (**Figure 6B**). P-site T_50_ correlations between brain regions (R =0.40-0.74) were markedly stronger than those between different tissues (R =0.13-0.53), although both were lower than the P-site abundance correlations, indicating the substantial diversity of P-site T_50_ (**Figure 6C** and **Figure S7**).

**Figure 6.**
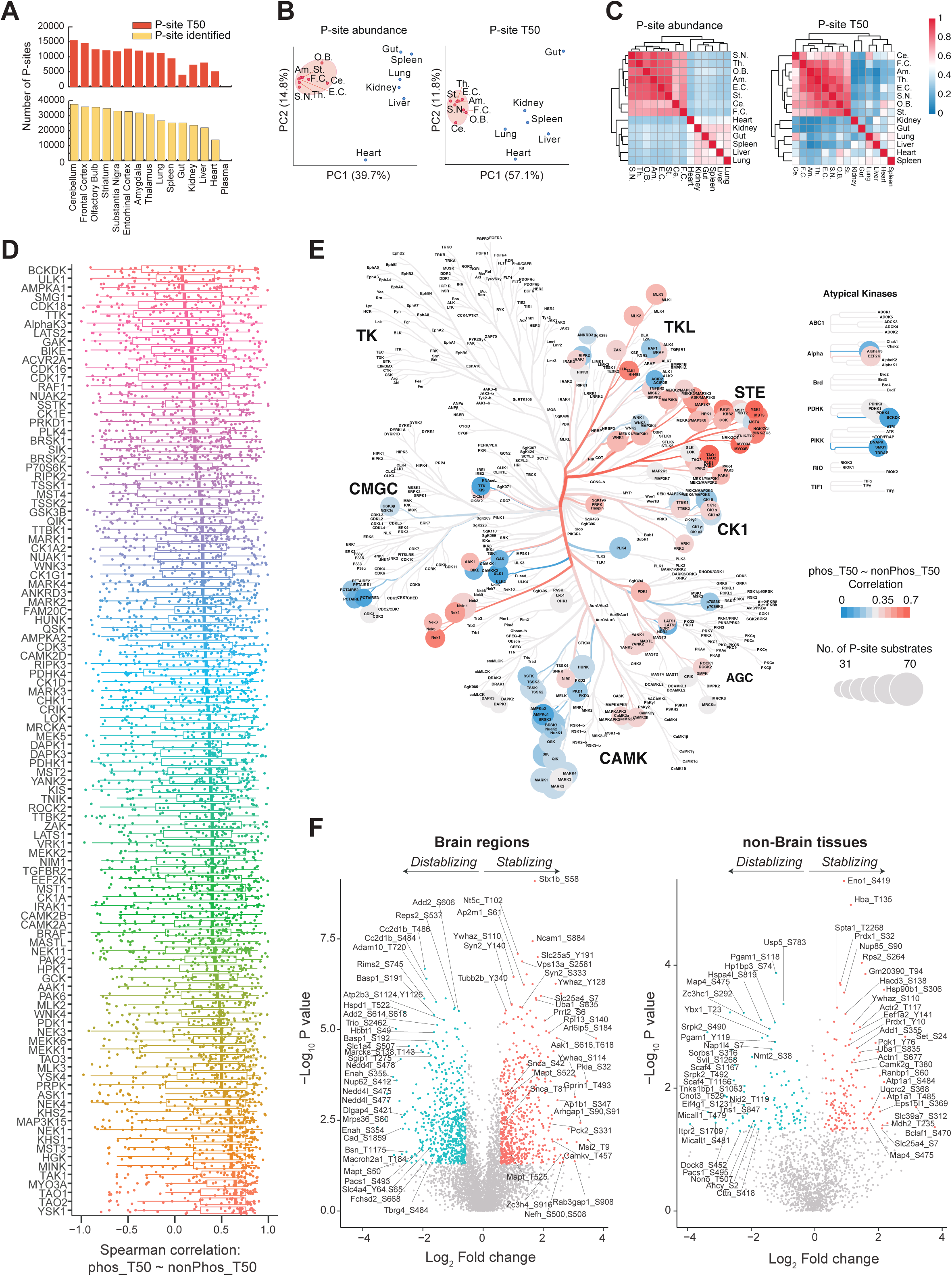
Profiling site-specific phosphorylation turnover and its impact across mouse tissues. (A) Number of quantified phosphorylation sites (P-sites) with abundance and lifetime values across tissues (Hippocampus not included for phosphoproteomics due to insufficient sample amount). (B) PCA plots of P-site abundance and lifetime across tissues. (C) Pearson correlation analysis of P-site abundance and lifetime between tissues, with the blue-to-red color bar indicating increasing Pearson correlation coefficients. (D) Distribution of Spearman correlation between T_50_ of the phosphorylated (phos_T50) and non-phosphorylated peptides (nonPhos_T50) for all specific P-sites across tissues, mapped to the kinase library based on kinase-substrate annotation. The 106 kinases with 30 or more putative P-site substrates (Percentile >0.99) quantified with respective T_50_ are shown. (E) Mapping of Spearman correlation between T_50_ of the phosphorylated (phos_T50) and non-phosphorylated peptides (nonPhos_T50) across tissues onto a kinase phylogenetic tree. The size of the kinase nodes represents the number of P-site substrates, and the blue-to-red color bar indicates increasing Spearman correlation coefficients (*rho*). (F) Volcano plots showing P-sites that increase or delay protein turnover (i.e., destabilizing or stabilizing the corresponding protein) across brain regions and non-brain tissues. The fold change in PT was determined by comparing phosphopeptides to non-phosphopeptides of the same peptide sequence. P-values were calculated using Student’s t-test. Blue and red dots denote the significant P-sites (P-value <0.05, Student’s t-test) showing the |fold change| >1.5 (in brain) and >1.2 (in non-brain tissues).

To explore how site-specific phosphorylation alters protein turnover across different tissues, we harnessed our previously developed DeltaSILAC method ^22^ by considering the non-phosphorylated protein forms ^20^. DeltaSILAC essentially integrates pSILAC, phosphoproteomics, and a peptide-level matching strategy ^22 20 25^ and was initially applied to growing HeLa cells ^22 23 24^. Herein, for a given P-site, we therefore compared the T_50_ of a phosphorylated (p) peptide to the T_50_ of its non-phosphorylated (np) counterpart within the same tissue’s whole proteomic results. We firstly distributed the correlation between T_50_ values of p and np peptides for all P-sites across tissues (**Figure S7B**) to the recently established kinase library by the Cantley group and others ^77,78^. A total of 106 kinases, each covering more than 30 P-site substrates quantified with respective T_50_ pairs (Percentile >0.99), were evaluated (**Figure 6D**). We found the phosphorylation-induced T_50_ alteration for the substrates of the same kinase can vary significantly. Mapping this result to kinase phylogenetic tree, we observed that substrates of Calcium/Calmodulin-dependent Protein Kinases (CAMK family) showed weak T_50_ cross-tissue consistency between p and np peptides overall (e.g., *rho* =0.297-0.340 for MARK kinases, **Figure 6E**). In contrast, the Serine/Threonine Kinases (STE family) present strongest corresponding T_50_ correlations, such as YSK1 (*rho* =0.700), TAO1 (*rho* =0.650), TAO2 (*rho* =0.650), and MYO3A (*rho* =0.646), much higher than CAMK’s results (*P* <2.2e-16). Thus, after being phosphorylated by a particular kinase, a P-site lifetime and the corresponding protein-level lifetime might be regulated by independent *in vivo* mechanisms. Furthermore, leveraging the high data completeness of T_50_ across brain regions, we also conducted a 2D functional enrichment analysis using protein-level functional annotation (**Figure S8**). We found that the abundance differences of P-sites across brain regions are particularly relevant to biological processes such as Electron transport chain (P =0.00995), Endosome (P =0.00601), Response to ER stress (P =0.0317). On the other hand, P-site lifetime variability in the brain tends to be associated with different processes, such as Actin nucleation (P =0.0319) and Positive regulation of protein localization in plasma membrane (P =0.0466). This suggests that the distribution of P-site abundance and the duration of P-site presence across brain regions are critical to the brain’s complex functions. The above findings together demonstrate that phosphorylation preferentially influences the stability of proteins according to their functional roles.

Next, we directly determined the real T_50_ difference for p and np pairs (i.e., ΔT_50_). In growing cells, more P-sites tended to associated with higher T_50_ ^22 23^, a trend that was found not as evident in tissue results (**Figure S7C**). Using volcano plots (**Figure 6F**), we identified 581 and 592 P-sites significantly extending or shortening T_50_ across brain regions (P <0.05, |FC| >1.5), and 146 and 105 P-sites doing the same across other tissues (P <0.05, |FC| >1.2). Based on the kinase library annotation ^77,78^, P-sites accelerating turnover in the brain were enriched as substrates for kinases TSSK2 (P =0.0160) and HUNK (P =0.0407), while those stabilizing proteins were enriched as substrates for SIK (P =0.0225), NEK11 (P =2.89E-4), and others (**Figure S7D**). Furthermore, for extremely *in vivo* long-lived proteins (ELLPs) such as nucleoporins, histone variants, and enzymes ^45 79^, our data here identified specific P-sites that further extend or fine-tune their PT (**Figure S7E**). Together, our results essentially profiled and prioritized P-sites based on their linkage to *in vivo* protein stability. And profiling Heavy/Light (H/L) ratios of p- and np-peptides for key P-sites and crucial proteins might provide new evidence on their functional roles and enable new opportunities to target them (see examples **Figure 7A**).

**Figure 7.**
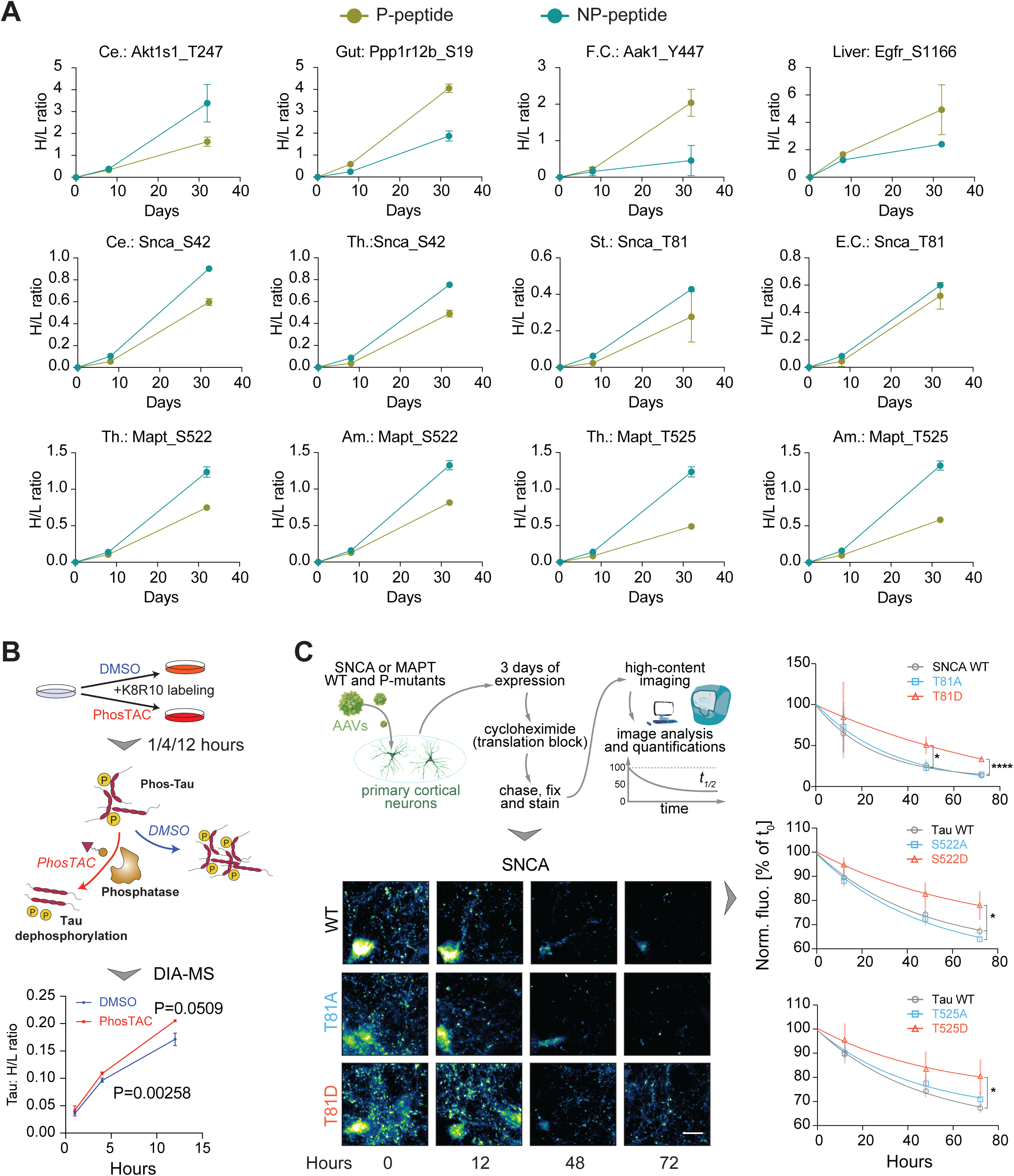
Demonstration and verification of phosphorylation sites (P-sites) linked to protein turnover in mouse tissues. (A) Heavy/Light (H/L) ratio curve examples during the labeling course for a phosphorylated (p) peptide and its non-phosphorylated (np) peptide counterpart of the same sequence and protein. (B) Validation of phosphorylation’s stabilizing effect on Tau protein using the PhosTAC approach. Upper panel: The pSILAC experiment comparing Tau protein turnover after PhosTAC or DMSO treatments. Lower panel: Heavy-to-light ratio curves during treatment and pSILAC labeling. P-values were calculated using Student’s t-test. (C) Validation of Tau and alpha-synuclein P-sites associated with protein degradation in primary hippocampal cortical neurons. Left panel: Neurons were infected with FLAG-tagged alpha-synuclein or Tau, either as wild type (WT) or mimicking mutants dephosphorylated (T/S to A) or phosphorylated (T/S to D). After three days of expression, neurons were treated with cycloheximide, chased for different times, stained, automatically imaged, and FLAG fluorescence intensity was measured. Right panel: Fluorescence imaging results from three independent experiments for alpha-synuclein (T81) and Tau (MAPT, S522, T525). Bars in graphs represent SEM. Statistical test: ANOVA. *p < 0.05; ****p < 0.0001. Scale bar: 5 µm.

To verify the impact of phosphorylation on the degradation of key proteins, we focused on microtubule-associated protein Tau and α-synuclein, both of which are widely recognized for their crucial roles in neurodegenerative diseases. Our DeltaSILAC analysis determined that phosphorylation at S522 and T525 of Tau significantly extended its PT by 7.24 and 9.20 days, respectively, across brain regions. Similarly, phosphorylation at T81 of α-synuclein markedly prolonged its PT by 13.20 days on average (**Figure 7A**). Consistently, hyperphosphorylated Tau was reported to promote aggregation and self-assembly into paired helical filaments tangles^80^, affecting protein degradation ^81^. Promoting the removal of Tau phosphorylation might offer therapeutic potential. Similarly, the phosphorylation of α-synuclein affects its aggregation and neurotoxicity ^82^. In the first validation approach, we employed a recently developed phosphorylation-targeting chimera (PhosTAC) technology to promote Tau dephosphorylation through induced proximity with the active PP2A holoenzyme ^83^. Unlike proteolysis-targeting chimeras (PROTACs) that induce selective intracellular proteolysis, PhosTACs induce rapid and sustained protein dephosphorylation ^83 84^. We firstly confirmed the downregulation of multiple Tau phosphorylation sites by PhosTAC treatment ^83^. Then we measured the H/L ratios of Tau in an *in vitro* system, where the cell culture medium was replaced with heavy SILAC during PhosTAC administration (**Figure 7B**). We observed significantly accelerated Tau protein degradation (i.e., shorter PT) following PhosTAC treatment, indicating the potential to regulate Tau protein clearance through its dephosphorylation. As another independent verification, we overexpressed the wild-type and phosphomimetic mutants of Tau and α-synuclein in primary rodent cortical neurons. We then measured their lifetimes by imaging after cycloheximide blockade, as previously described ^22^ (**Figure 7C**). Our results indicate that for α-synuclein, the phosphomimetic mutation T81D significantly increases the protein’s lifetime by 3.81-fold compared to the wild-type. The Tau phosphomimetic mutations T522D and T525D exhibited analogous effects. In summary, the phosphoproteome turnover dataset included in *Tissue-PPT* provides promising opportunities for exploring the dynamics and turnover of individual P-sites.

## DISCUSSION

Resolving the molecular specifics of proteome level variation across distinctive mammalian tissues and organs will significantly advance our comprehension of physiology and disease. Elucidating protein turnover in tissues, can uncover how different tissues and organs develop their distinct phenotypes, coordinate their respective functions, and respond to various stimuli. In this study, we establish *Tissue-PPT*, a monumental inventory of proteome and phosphoproteome turnover across multiple tissues and brain regions, including more than 256,000 PT measurements in total. Our whole protein turnover profiling has tripled the number of protein analytes on average, and our phosphoproteomic dataset is entirely novel. *Tissue-PPT* identifies short-lived and long-lived proteins and pathways within and among tissues, and additionally reveals how phosphorylation is associated with the *in vivo* protein stability, both in detail and in large-scale. The *Tissue-PPT* Web App supports convenient navigation and discovery of PA and PT profiles for individual proteins and protein sets. Collectively, our datasets and analyses represent a significant step towards the molecular understanding of tissue phenotypic and functional diversity, elucidating not only the composition of a proteome and the quantity of its constituents, but also the lifespan and activity of each individual proteins among tissues and brain regions.

The high-coverage, precise, and reproducible determination of protein turnover kinetics reported in this study stems from the advanced MS techniques we employed. Both DIA and TMTpro were devised to improve the quantitative accuracy via either the gas-phase or LC fractionations ^28 32 30^, significantly ameliorating the missing value issue ^85^ in traditional data-dependent acquisition (DDA) based shotgun proteomics for multiplexed analysis. The *in vivo* SILAC strategy based on Lysine-6 was used to avoid the complexities associated with ^15^N labeling which requires sophisticated process algorithm ^86^ and potentially leads to biased precision of measurement ^9^ and smaller proteome coverage ^10^.

The results included in *Tissue-PPT* underscore the critical importance of integrating *well-matched* datasets across diverse biological contexts.

***First***, matching PA with PT profiling enables the quantification of protein turnover’s role in determining biological diversity. This analysis, valuable for understanding cardiac remodeling^35^, has now been extended across multiple tissues. Conceivably, the observed positive PA-PT correlation indicates concerted post-transcriptional regulation. Proteins with the highest PAs and PTs can be considered housekeeping proteins essential for tissue function diversity. Conversely, maintaining shorter PTs for the most abundant proteins allows for rapid responses to environmental perturbations. Stabilizing low-abundance proteins, particularly those in protein complexes and rate-limiting enzymes, may promote proteome buffering. Our HT plots, using a blue-to-red color gradient to indicate turnover rates, succinctly summarize these reciprocal PA-PT relationships.

***Second***, matching phosphorylated and non-phosphorylated (p and np) peptides ensures precise estimation of turnover effects due to site-specific phosphorylation ^22^. This is essential, given that many proteoforms carrying other PTMs were not profiled ^20^. Using this approach, we discovered and confirmed that *in vivo* phosphorylation can alter the stability of key proteins in different tissues, such as Tau and α-synuclein. Our results therefore emphasize that phosphorylation is not only crucial in incurring rapid cell signaling response, but also in regulating protein stability in steady-state tissues, possibly through kinase selectivity and variable mechanisms such as PPI ^25 24^. Concurrent turnover analysis of phosphorylation sites encoded by the same gene will significantly enhance our current understanding of phosphoproteomics ^87^ and guide future biochemical and functional analyses of specific P-sites.

***Third***, matching tissue-specific PTs with PPIs revealed a previously underestimated correlation. Prior inter-partner correlation was achieved from tissue-resolved PPIs rather than Bioplex and CORUM annotations. Accordingly, our findings indicate that most PPI partners have similar PTs, and deviations from this trend suggest additional moonlighting functions. These results may help inferring PPI networks and understanding proteome organization.

***Fourth***, matching multi-layered omic profiles across multiple tissues provides powerful insights into how cells orchestrate organellar pathways to maintain tissue diversity. Our analysis presents the first comprehensive characterization of multiple tissues incorporating transcriptome, translatome, proteome, proteome turnover, and phosphoproteome quantifications. For instance, while peroxisome proteins were previously reported to have variable PTs ^9^, our multi-omic analysis uncovers that they also exhibit significant variability at the mRNA and PA levels, and that strikingly, peroxisomes are regulated primarily by PT to counteract PA variability. A similar but less-pronounced trend was observed for mitochondrial proteome. As the reference for peroxisome, our results underscored the preeminence of transcriptional and translational processes in shaping the global tissue-specific proteomes, in which turnover playing a lesser role in general. Discovering these fundamental regulatory mechanisms will be difficult without matched multi-omics analysis ^88^. We observed the significant PT regulation for pexophagy-associated proteins involved in autophagosome biogenesis and peroxisome designation ^71^. The molecular mechanism associating peroxisome turnover with tissue phenotype, however, remains to be established. By employing multiple *matching* strategies during data generation and interpretation, our study significantly extends the current understanding of protein post-translational regulation and turnover control in mammals.

Other concrete findings include an unexpected turnover trend that K48-linked ubiquitin is replaced slower across tissues compared to K63-linked ubiquitin, which might suggest that K48-linked Ub is recycled by proteasome-associated DUBs during degradation which are expressed stable across tissues ^50^, whereas K63-linked Ub is degraded primarily by the lysosome ^52^. Furthermore, the variable PT of E3-associated proteins also points to the potential for developing tissue-specific protein degraders as therapeutic modules targeting underexplored E3 ligases ^89^.

### Limitations of the study

It should be emphasized that the PT values in our study represent the *de facto* protein turnover in tissues within a live organism. While these values accurately describe protein dynamics and are relevant for *in vivo* experiments such as drug discovery where compounds are administered to whole-body animals, they differ from the *cellular protein turnover rates*. Indeed, the protein clearance on a per-cell basis is influenced by the combined effects of degradation kinetics and cellular dilution due to cell division. For instance, the global PT is shortest in the gut, likely due to the rapid cell turnover in gut. While ^15^N labeling yielded technical challenges ^10 90^, it was recently used to track both protein and DNA labeling in mouse tissues via a technique termed TRAIL^13^ to account for tissue proliferation differences when comparing protein turnover. This means that our *absolute* PT data should be applied cautiously in experiments comparing proliferative tissues versus post-mitotic tissue types. We however note that neither our approach nor TRAIL can resolve protein turnover for different cell types within tissues, which may significantly impact proteome readouts ^91–93^. In this regard, a single-cell proteomic turnover analysis ^94^ for all different cells within and across tissues might be needed. Furthermore, we discovered intriguing patterns in PA and PT suggesting bimodal biological origins of plasma proteins. However, the limited coverage of the plasma proteome and phosphoproteome by mass spectrometry precluded a deeper investigation in the present study. Lastly, while we verified several phosphorylation events linked to altered protein stability in brain cells, mechanistically defining the relationship between specific phosphorylation sites and protein stability is beyond the scope of the present study.

In conclusion, we established a high-quality comprehensive resource portraying the protein turnover dynamics in mammalian tissues, providing deep and novel insights into the proteostasis regulation underlying tissue phenotypic and functional diversity.

## ACKNOWLEDGMENTS

Y.L. thanks the support from the National Institute of General Medical Sciences (NIGMS), National Institutes of Health (NIH) through Grant R01GM137031 and RM1GM149406. J.P. is supported by RF1AG064909 and RF1AG068581 from NIH. EFF is supported by a CZI Collaborative Pairs Pilot Project Awards (Cycle 2; Phase 1). EFF also acknowledges the support of the SFB1286, Göttingen, Germany. We would like thank Drs. Mandar Muzumdar and Ines Chen for their critical comments on the manuscript.

## AUTHOR CONTRIBUTION

W.L. performed DIA-MS measurements, conducted phosphoproteomic experiments, prepared figures and tables, and led the data analysis under the supervision of Y.L. A.D. processed the protein and phosphoproteomic lifetime data and conducted bioinformatic analyses. K.Y. prepared the proteomic samples and carried out TMT experiments with assistance from J.M.Y. S.W. developed the Tissue-PPT web portal. N.H.K. conducted verification experiments for phosphorylation-associated turnover under the supervision of E.F.F. Z.H. performed the PhosTAC experiments. B.S. contributed to data analysis. J.P., and Y.L. conceptualized the study. E.F.F., J.P., and Y.L. secured funding and co-supervised the entire project. Y.L. wrote the initial manuscript with input from all authors. All authors contributed to the writing of the final manuscript.

## STAR METHODS

### KEY RESOURCES TABLE

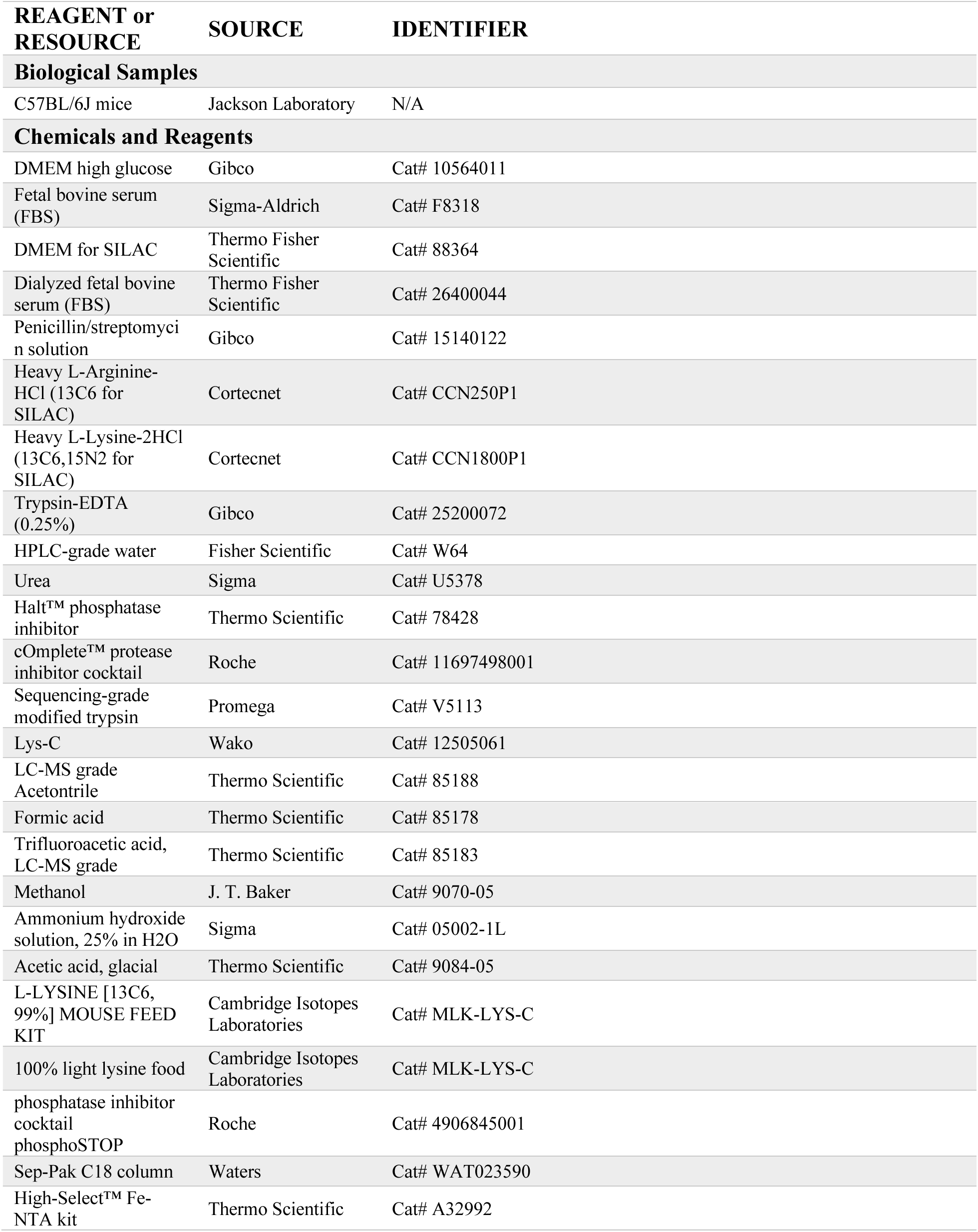
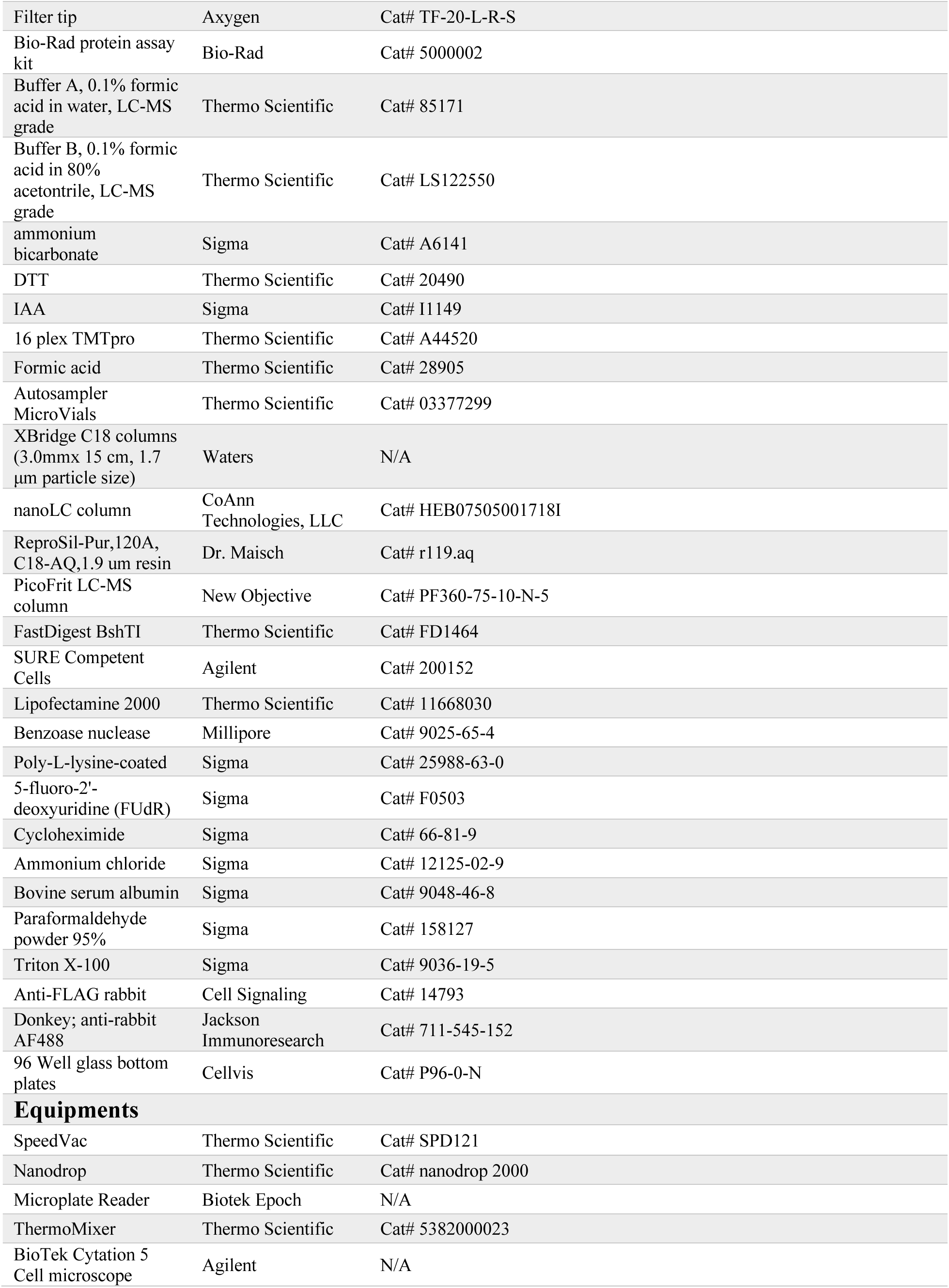
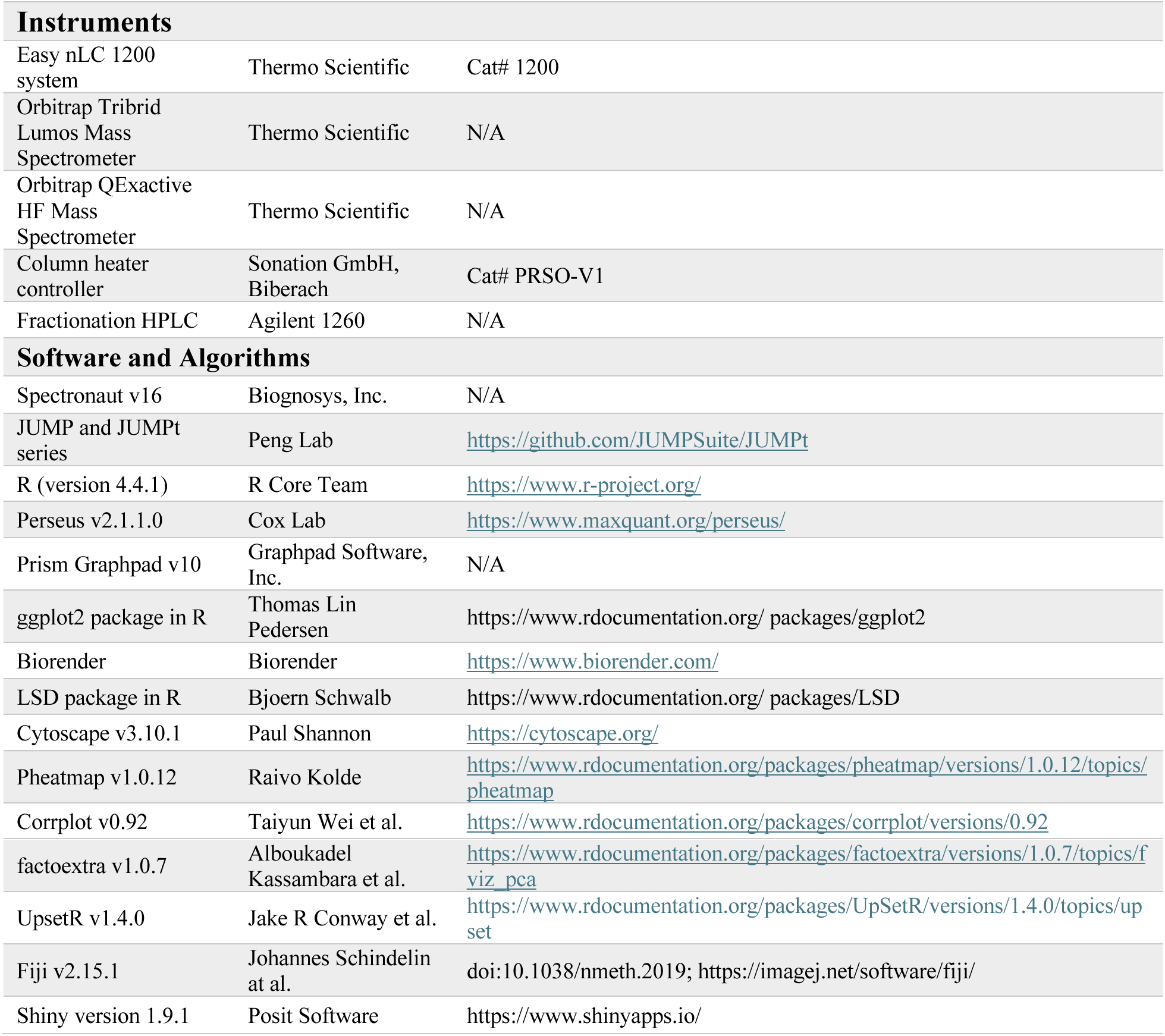

| REAGENT or RESOURCE | SOURCE | IDENTIFIER |
| --- | --- | --- |
| **Biological Samples** |  |  |
| C57BL/6J mice | Jackson Laboratory | N/A |
| **Chemicals and Reagents** |  |  |
| DMEM high glucose | Gibco | Cat# 10564011 |
| Fetal bovine serum (FBS) | Sigma-Aldrich | Cat# F8318 |
| DMEM for SILAC | Thermo Fisher Scientific | Cat# 88364 |
| Dialyzed fetal bovine serum (FBS) | Thermo Fisher Scientific | Cat# 26400044 |
| Penicillin/streptomycin solution | Gibco | Cat# 15140122 |
| Heavy L-Arginine-HCl (13C6 for SILAC) | Cortecnet | Cat# CCN250P1 |
| Heavy L-Lysine-2HCl (13C6,15N2 for SILAC) | Cortecnet | Cat# CCN1800P1 |
| Trypsin-EDTA (0.25%) | Gibco | Cat# 25200072 |
| HPLC-grade water | Fisher Scientific | Cat# W64 |
| Urea | Sigma | Cat# U5378 |
| Halt™ phosphatase inhibitor | Thermo Scientific | Cat# 78428 |
| cOmplete™ protease inhibitor cocktail | Roche | Cat# 11697498001 |
| Sequencing-grade modified trypsin | Promega | Cat# V5113 |
| Lys-C | Wako | Cat# 12505061 |
| LC-MS grade Acetonitrile | Thermo Scientific | Cat# 85188 |
| Formic acid | Thermo Scientific | Cat# 85178 |
| Trifluoroacetic acid, LC-MS grade | Thermo Scientific | Cat# 85183 |
| Methanol | J. T. Baker | Cat# 9070-05 |
| Ammonium hydroxide solution, 25% in H<sub>2</sub>O | Sigma | Cat# 05002-1L |
| Acetic acid, glacial | Thermo Scientific | Cat# 9084-05 |
| L-LYSINE [13C6, 99%] MOUSE FEED KIT | Cambridge Isotopes Laboratories | Cat# MLK-LYS-C |
| 100% light lysine food | Cambridge Isotopes Laboratories | Cat# MLK-LYS-C |
| phosphatase inhibitor cocktail<br>phosphoSTOP | Roche | Cat# 4906845001 |
| Sep-Pak C18 column | Waters | Cat# WAT023590 |
| High-Select™ Fe-NTA kit | Thermo Scientific | Cat# A32992 |
| Filter tip | Axygen | Cat# TF-20-L-R-S |
| Bio-Rad protein assay kit | Bio-Rad | Cat# 5000002 |
| Buffer A, 0.1% formic acid in water, LC-MS grade | Thermo Scientific | Cat# 85171 |
| Buffer B, 0.1% formic acid in 80% acetonitrile, LC-MS grade | Thermo Scientific | Cat# LS122550 |
| ammonium bicarbonate | Sigma | Cat# A6141 |
| DTT | Thermo Scientific | Cat# 20490 |
| IAA | Sigma | Cat# I1149 |
| 16 plex TMTpro | Thermo Scientific | Cat# A44520 |
| Formic acid | Thermo Scientific | Cat# 28905 |
| Autosampler MicroVials | Thermo Scientific | Cat# 03377299 |
| XBridge C18 columns (3.0mmx 15 cm, 1.7 µm particle size) | Waters | N/A |
| nanoLC column | CoAnn Technologies, LLC | Cat# HEB07505001718I |
| ReproSil-Pur,120A, C18-AQ,1.9 µm resin | Dr. Maisch | Cat# r119.aq |
| PicoFrit LC-MS column | New Objective | Cat# PF360-75-10-N-5 |
| FastDigest BshTI | Thermo Scientific | Cat# FD1464 |
| SURE Competent Cells | Agilent | Cat# 200152 |
| Lipofectamine 2000 | Thermo Scientific | Cat# 11668030 |
| Benzoase nuclease | Millipore | Cat# 9025-65-4 |
| Poly-L-lysine-coated | Sigma | Cat# 25988-63-0 |
| 5-fluoro-2'-deoxyuridine (FUdR) | Sigma | Cat# F0503 |
| Cycloheximide | Sigma | Cat# 66-81-9 |
| Ammonium chloride | Sigma | Cat# 12125-02-9 |
| Bovine serum albumin | Sigma | Cat# 9048-46-8 |
| Paraformaldehyde powder 95% | Sigma | Cat# 158127 |
| Triton X-100 | Sigma | Cat# 9036-19-5 |
| Anti-FLAG rabbit | Cell Signaling | Cat# 14793 |
| Donkey; anti-rabbit AF488 | Jackson ImmunoResearch | Cat# 711-545-152 |
| 96 Well glass bottom plates | Cellvis | Cat# P96-0-N |
| **Equipments** |  |  |
| SpeedVac | Thermo Scientific | Cat# SPD121 |
| Nanodrop | Thermo Scientific | Cat# nanodrop 2000 |
| Microplate Reader | Biotek Epoch | N/A |
| ThermoMixer | Thermo Scientific | Cat# 5382000023 |
| BioTek Cytation 5 Cell microscope | Agilent | N/A |

| **Instruments** |  |  |
| --- | --- | --- |
| Easy nLC 1200 system | Thermo Scientific | Cat# 1200 |
| Orbitrap Tribrid Lumos Mass Spectrometer | Thermo Scientific | N/A |
| Orbitrap QExactive HF Mass Spectrometer | Thermo Scientific | N/A |
| Column heater controller | Sonation GmbH, Biberach | Cat# PRSO-V1 |
| Fractionation HPLC | Agilent 1260 | N/A |
| **Software and Algorithms** |  |  |
| Spectronaut v16 | Biognosys, Inc. | N/A |
| JUMP and JUMPt series | Peng Lab | https://github.com/JUMPSuite/JUMPt |
| R (version 4.4.1) | R Core Team | https://www.r-project.org/ |
| Perseus v2.1.1.0 | Cox Lab | https://www.maxquant.org/perseus/ |
| Prism Graphpad v10 | Graphpad Software, Inc. | N/A |
| ggplot2 package in R | Thomas Lin Pedersen | https://www.rdocumentation.org/packages/ggplot2 |
| Biorender | Biorender | https://www.biorender.com/ |
| LSD package in R | Bjoern Schwalb | https://www.rdocumentation.org/packages/LSD |
| Cytoscape v3.10.1 | Paul Shannon | https://cytoscape.org/ |
| Pheatmap v1.0.12 | Raivo Kolde | https://www.rdocumentation.org/packages/pheatmap/versions/1.0.12/topics/pheatmap |
| Corrplot v0.92 | Taiyun Wei et al. | https://www.rdocumentation.org/packages/corrplot/versions/0.92 |
| factoextra v1.0.7 | Alboukadel Kassambara et al. | https://www.rdocumentation.org/packages/factoextra/versions/1.0.7/topics/fviz_pca |
| UpsetR v1.4.0 | Jake R Conway et al. | https://www.rdocumentation.org/packages/UpSetR/versions/1.4.0/topics/upset |
| Fiji v2.15.1 | Johannes Schindelin at al. | doi:10.1038/nmeth.2019; https://imagej.net/software/fiji/ |
| Shiny version 1.9.1 | Posit Software | https://www.shinyapps.io/ |

### RESOURCE AVAILABLITY

#### Lead Contact

Further information and requests should be directed to and will be fulfilled by the lead author.

#### Materials Availability

This study did not generate new unique reagents.

### EXPERIMENTAL MODEL AND SUBJECT DETAILS

The B6SJL (C57BL/6 x SJL) mice were purchased from the Jackson Laboratory. Mice were maintained in the Animal Resources Center at St. Jude Children’s Research Hospital according to the Guidelines for the Care and Use of Laboratory Animals. All animal procedures were approved by Institutional Animal Care and Use Committee (IACUC) at St. Jude Children’s Research Hospital. Male mice of approximately 9 months were used for global protein turnover profiling. Mice were maintained on a 12:12 h light/dark cycle in a temperature and humidity-controlled room with food and water *ad libitum*.

### METHOD DETAILS

#### *In vivo* Pulsed SILAC Labeling and Tissue Dissection

Each mouse was provided with 5 g SILAC food (Mouse Express L-LYSINE [13C6, 99%] MOUSE FEED kit, Cambridge Isotopes Laboratories) per day for metabolic labeling *in vivo*. Three days prior to metabolic labeling, mice were fed with the SILAC food composed of 100% light lysine to minimize the perturbation of protein homeostasis due to the switching from regular food to SILAC food. The mice were then fed with SILAC food for specified periods before sacrificed by cervical dislocation. All the anatomical samples of body tissues (heart, liver, spleen, lung, kidney, gut, plasma) and brain regions (cerebellum, frontal cortex, substantia nigra, thalamus, amygdala, entorhinal cortex, hippocampus, and olfactory bulb) were dissected rapidly, frozen in liquid nitrogen, and stored at −80 °C.

#### Tissue Protein Extraction and Digestion

Protein extraction and MS analysis were performed based on an optimized protocol^95^. About 20 mg of the mouse tissue was weighed and lysed in ∼200 μL lysis buffer (8 M urea, 50 mM HEPES, pH 8.5, 0.5% sodium deoxycholate, phosphatase inhibitor cocktail (phosphoSTOP, Roche)) at 4°C in a bullet blender. Protein concentration was measured by the BCA assay. Lysate containing ∼1 mg proteins were digested with Lys-C (Wako, 1:100 w/w) in lysis buffer at 21 °C for 3 h. The digested mixture was diluted 4 times with 50 mM HEPES (pH 8.5) to reduce urea to 2 M, and digested with trypsin (Promega, 1:50 w/w) overnight at 21 °C. The digestion condition was selected to ensure sufficient protein digestion while minimizing the potential for urea-derived protein carbamylation. The digested peptides were reduced by freshly prepared dithiothreitol (DTT, 1 mM) for 2 h, followed by alkylating with 10 mM iodoacetamide (IAA) in the dark for 30 min. The unreacted IAA was quenched by adding DTT to 30 mM and incubate for 30 min at RT. The samples were then acidified by addition of 1% trifluoroacetic acid (TFA). Acidification of peptides by trifluoroacetic acid was followed with desalting on Sep-Pak C18 column (Waters). Samples were split into two parts for DIA and TMT-DDA analyses and dried by SpeedVac.

#### TMTpro Labeling and basic pH HPLC Fractionation

About 100 µg of dried peptides were resuspended in 50 mM HEPES (pH 8.5) and labeled by 16-plex TMTpro reagent (Thermo Fisher Scientific, ∼1:2 w/w) ^96^. Peptides labeled with each channel of TMT was mixed equally and desalted using Sep-Pak C18 column (Waters). The TMT-labeled peptides were fractionated by offline basic pH reverse phase LC (RPLC). Injected peptides were separated using two tandem XBridge C18 columns (3.0 mm x 15 cm, 1.7 μm particle size, Waters) in a 3-h 10-45% gradient (buffer A: 10 mM ammonium formate, pH 8.0; buffer B: 95% acetonitrile, 10 mM ammonium formate, pH 8.0) to yield total of 80 or 96 concatenated fractions.

#### Phosphopeptide Enrichment

About 500 μg peptide per each sample was used for phosphoproteomic sample preparation^28^. The phosphopeptide enrichment was performed using High-Select™ Fe-NTA kit (Thermo Scientific, #A32992) according to the kit instructions, as described previously^97^. Briefly, the resins of one spin column in the kit were divided into 5 equal aliquots, each used for one sample. The peptide-resin mixture was mixed and incubated for 30 min at 21 °C and gently shake per 10 min, and then transferred into the filter tip (TF-20-L-R-S, Axygen) to remove the supernatant by centrifugation. Then the resins adsorbed with phosphopeptides were washed sequentially with 200 µL × 3 washing buffer (80% ACN, 0.1% TFA) and 200 µL × 3 H_2_O to remove nonspecifically adsorbed peptides. The phosphopeptides were eluted off the resins by 100 µL × 2 elution buffer (50% ACN, 5% NH_3_•H_2_O) and dried with SpeedVac (Thermo Scientific). All centrifugation steps above were conducted at 500 g × 30 s. The eluates were dried immediately and resuspended with buffer A for mass spectrometry analysis. ∼1.5 μg phosphopetide was injected into mass spectrometry for phosphoproteomic analysis.

#### BoxCarmax-DIA and DIA Mass Spectrometry

The total pulsed SILAC proteome samples (digested peptides) were measured by the BoxCarmax-DIA method optimized for protein turnover analysis ^32^. The Orbitrap Fusion Lumos Tribrid mass spectrometer (Thermo Scientific) was coupled with a NanoFlex ion source was used for the data acquisition. The spray voltage at 2000 V and heating capillary at 275 °C. Briefly, one BoxCarmax consist of four MS runs (1^st^, 2^nd^, 3^rd^ and 4^th^ injection) to reconstruct a full MS1 scan ^32^. Each run took 60 min. The MS1 AGC was set to be 2 × 10^6^ and the maximum injection time was set at 256 ms. The MS1 resolution was 120,000 at m/z 200 and the normalized HCD collision energy was 28%. The MS2 AGC was set to be 1.5 × 10^6^ and the maximum injection time was 50 ms. The MS2 resolution was set to be 30 000 and the MS2 scan range was 200−1800 m/z. Both MS1 and MS2 spectra were recorded in profile mode. All the phosphorylation samples were measured by a traditional DIA method including a 150-min gradient^98,99^ to ensure the correct detection and analysis on the same LC-MS of the phosphopeptide samples (that usually have low amounts). The DIA-MS consisted of one MS1 scan and 33 MS2 scans with variable windows ^32 100^, Except for the MS1 scan range set to cover 350 – 1650 m/z, other MS1 and MS2 settings remain identical to those in BoxCarmax. To strictly match the phosphorylation data in DeltaSILAC analysis^22^, the total pSILAC proteomic samples were measured repeatedly by the same DIA method using a 240-min LC gradient. LC separation was performed on EASY-nLC 1200 systems (Thermo Scientific, San Jose, CA) using a 75 µm × 50 cm length column (CoAnn Technology). To elute peptides, Buffer B (80% acetonitrile containing 0.1% formic acid) from 5% to 37% and the corresponding buffer A (0.1% formic acid in H_2_O) were used in all the gradients. The flow rate was kept at 300 nL/min with the temperature controlled at 60 °C using a column oven (PRSO-V1, Sonation GmbH, Biberach, Germany).

#### TMTpro Mass Spectrometry

Around 200 ng of peptides from every basic pH HPLC fraction were loaded on a reverse phase column (75 µm × 25 cm, 1.7 µm C18 resin, CoAnn Technology) interfaced with a Q Exactive HF mass spectrometer (Thermo Fisher Scientific) ^101^. Peptides were eluted in a 90 min 10-35% gradient of buffer B (buffer A: 0.2% formic acid, 3% DMSO; buffer B: 67% acetonitrile, 0.2% formic acid, 3% DMSO). The mass spectrometer was operated in a data-dependent mode with MS1 set with 60,000 resolution, 1 × 10^6^ AGC target and 50 ms maximal ion time. The MS1 was followed by top 20 MS2 high resolution scans that were set as follows: 1.0 m/z isolation window, 0.2 m/z offset, 60,000 resolution, 110 ms maximal ion time, 1 × 10^5^ AGC target, HCD, 32% normalized collision energy, and 15 s dynamic exclusion.

#### Molecular Biology and Adeno-associated Virus (AAV) Production

SNCA and MAPT sequences were retrieved from Ensembl (https://www.ensembl.org/) and ordered in puc57-KANA from GenScript, flanked by restriction sites (AgeI/BshTI and SdaI/SbfI) for one-step insertion into AAV constructs. Appropriate mutations were synthesized. For detection and direct comparison in imaging experiments, a C-terminal 3X-FLAG sequence was included in the synthesized sequences. After insertion into the AAV backbone, a single cassette was generated under the control of a human synapsin 1 gene promoter ^102^. After ligation, AAVs plasmids were amplified in SURE competent cells (Agilent) to avoid ITR loss. Final constructs were verified by sequencing and the absence of ITRs was confirmed by DNA restriction analysis. Sequences from gene synthesis are provided as supplementary files, and all plasmids are available from the authors upon reasonable request. Viruses were prepared as previously described^102^ by cotransfection of helper plasmids with the target plasmid using Lipofectamine 2000 (Thermo Fisher). At 72-hour post-transfection, cells were harvested and lysed by 4 cycles of thawing and freezing followed by treatment with Benzoase nuclease (Millipore) and incubated at 37°C for ∼30 min. After pelleting cell debris (14000 rpm, 30 min at 4°C), supernatants were filtered, aliquoted and snap frozen in liquid nitrogen. Viruses were stored at −80°C until use. Viruses were titrated by imaging to achieve comparable expression levels upon FLAG-staining (see below).

#### Primary Cortical Neuron Preparation and Infection

Primary cortical cultures were prepared from P2 neonatal rats (*Rattus norvegicus*, Wistar) with minor adaptations to those previously described ^103,104^. Cortical neurons were plated on 1 mg/ml poly-L-lysine-coated (Sigma) 96-well glass bottom plates optimized for imaging (Cellvis) at a concentration of 30,000 cells per well and maintained at 37°C in 5% CO_2_. On the second day in vitro (DIV), 5-fluoro-2’-deoxyuridine (FUdR; Sigma) was added to the culture at a final concentration of 5 µM to prevent glial proliferation. At 5 DIV, neurons were infected with AAVs containing the sequence for the gene of interest. At 8 DIV, cells were treated with cycloheximide (0.5 µg/ml final, Sigma) and followed for different times to determine the dynamics of protein turnover.

#### Immunofluorescence, Imaging, and Analysis

At the end of the experiments, neurons were fixed for 30 minutes in 4% paraformaldehyde (PFA) in phosphate-buffered saline (PBS) at room temperature (RT). After rinsing with PBS, the cells were quenched with 10 mM ammonium chloride in PBS at RT for 15 minutes. The cells were then washed three times for 5 minutes, blocked and permeabilized in permeabilization buffer (PB) containing 4% bovine serum albumin and 0.1% (v/v) Triton X-100 for 30 minutes at room temperature (RT). Primary antibody (anti-FLAG rabbit; Cell Signaling cat. 14793) was applied at a final concentration of 1:1000 in PB buffer for 1.5 hours at RT with gentle shaking. After four 15-minute washes with PBS, the secondary antibody (donkey; anti-rabbit AF488, Jackson Immunoresearch cat. 711-545-152) was applied in PB buffer for 1 hour at RT with gentle shaking. After four 15-minute washes with PBS, cells were imaged using a BioTek Cytation 5 Cell microscope with constant illumination and exposure. Images were analyzed as described previously^105^. Experiments were repeated on 3 independent cultures. In each experiment, at least 3 wells for each time point were analyzed amounting overall to ∼3000 neurons per time point.

#### Tau-PhosTAC Experiment and Measurement

A Tau expressing cell line (HeLa tau/PP2A) was generated as previously described ^83^. The HeLa tau/PP2A expressing cells were treated with doxycycline (2 μg/mL) for 24 h to induce tau expression. The media was then replaced with fresh SILAC media supplemented with DMSO, PhosTAC (1 μM) for 1h, 4h or 12h before harvest. The cells were washed with precooled PBS twice and snap-frozen when still in the dish with liquid nitrogen. Subsequently, a lysis buffer containing 10 M urea and the cOmplete™ protease inhibitor cocktail (Roche, #11697498001) was added. All the content of the plate was transferred into 2 ml tube upon scraping and stored at - 80 °C until sample preparation. Cells in this lysis buffer were thawed and a VialTweeter device (Hielscher-Ultrasound Technology) was used to sonicate the samples (4 °C; 1 min; two cycles). Uoon sonication, the samples were centrifuged at 20,000 g for 1 hour to remove all the insoluble material. Protein concentration was measured using the Bio-Rad protein assay dye (Bio-Rad, cat. no. 5000006). Reduction and alkylation were carried out using 10 mM Dithiothreitol (DTT) for 1 hour at 56°C, followed by 20 mM iodoacetamide (IAA) in darkness for 45 minutes at room temperature. A precipitation-based digestion method was used here ^28 106^. Briefly, five volumes of precooled precipitation solution (50% acetone, 50% ethanol, and 0.1% acetic acid) were added to the sample vortex 30 s. After overnight incubation at −20 °C, the samples were centrifuged (20,000 x g; 4 °C; 40 min). The precipitate was washed with precooled 100% acetone, centrifuged (20,000 × g; 4 °C; 40 min), and the remaining acetone was evaporated in a SpeedVac. For protein digestion, 300 µL of 100 mM NH_4_HCO_3_ with sequencing grade porcine trypsin (Promega) at a trypsin-to-protein ratio of 1: 20 were added and incubated overnight at 37 °C. The resulting peptide samples were acidified with formic acid and desalted using a C18 column (MarocoSpin Columns, NEST Group INC.) according to the manufacturer’s instructions. The peptide concentration was assayed by nanodrop, 1 μg peptide was inject into mass spectrometry. The same DIA method as descripted above was used for the data acquisition. The Spectronaut software was used for data analysis (see below).

### QUANTIFICATION AND STASTISTICAL ANALYSIS

#### DIA Data Procession and Analysis

The DIA-MS data analyses were performed using Spectronaut version 16 ^44,107^. All the raw datasets were firstly used for the library generation by the Pulsar search of Spectronaut. For the pulsed SILAC DIA library generation, the labels were specified in the “Labeling” setting, the “Labeling Applied” option was enabled, the “Lys6” were specified as “SILAC labeling” in the second channel, and the “In-Silico Generate Missing Channels” and “Label” in the Workflow setting were selected. Methionine oxidation was set as variable modification and cysteine carbamidomethylation was selected as fixed modification. For the phosphoproteomic data, the phosphorylation at S/T/Y was enabled as variable modification.

For the targeted data extraction of the pulsed SILAC datasets and for the subsequent identification and quantification, the Inverted Spike-In workflow (ISW) was used as described previously ^67^. The “Qvalue” was selected for all data filtering. Both peptide and protein FDR cutoffs were controlled at 1%. For the phosphoproteomic data, the probability of PTM cutoff was strictly kept at >0.75 to ensure the phosphosites were localized ^108^, similar to Class I confidence ^109,110^. The PTM score >0.01 table was also exported from the Spectronaut and then filtered by the PTM score >0.75 result for the following data analysis ^22,111^. The number and turnover of phosphosites were summarized based on the unique phosphopeptidoform level (i.e., the phosphopeptides with multiple modifications were regarded as different P-sites), as previously described ^22,111^. The pulsed SILAC plasma dataset was analyzed separately due to the low number of plasma protein identities which may potentially impact protein FDR control. The unlabeled tissues data (day 0) were analyzed by directDIA algorithm on Spectronaut 16. All the other Spectronaut settings were kept as Default. The quantitative results for heavy and light peptide precursors were exported by Spectronaut for following up turnover calculations. For the protein lifetime calculation in JUMPt ^16^, the peptide heavy-to-light ratio was initially filtered based on a time-dependent increase (e.g., 32d > 8d), then summarized to the protein level. The protein iBAQ value ^43^ was also directly exported from Spectronaut. For the phosphopeptide turnover calculation in JUMPt, both phosphopeptides and their corresponding non-phosphopeptides were analyzed using the same free lysine turnover curve, which was independently measured by DIA-MS from the same set of samples.

#### TMTpro Data Procession and Analysis

The JUMP search engine transformed peptide and protein identification by combining pattern-based scoring and de novo tag scoring, significantly improving the accuracy of peptide-spectrum matches (PSMs), as previously demonstrated ^112^. This innovation utilized MS/MS raw data and a composite target/decoy database; a concept introduced to estimate false discovery rates (FDR) ^113^. Generating a decoy database involved reversing target protein sequences and merging them with the accurate target database. FDR calculations were based on the (nd/nt) formula, assuming uniform mismatch distribution. The UniProt Mouse database (59,423 entries) was used to create the SILAC-TMT mouse protein database. Mass tolerances for precursor and fragment ions were set to 15 ppm and 20 ppm, respectively. Up to two missed cleavage sites were permitted per peptide. TMTpro labeling at Lys or the N-terminus, along with Cys carbamidomethylation, were defined as static modifications, while Met oxidation was treated as a dynamic modification. Protein FDR was maintained below 1% by applying filters based on mass accuracy and JUMP-based matching scores (Jscore and ΔJn). Following the rule of parsimony, peptides shared by multiple proteins were generally assigned to the canonical protein form. In cases where no canonical form was defined in the database, the peptide was assigned to the protein with the highest peptide-spectrum match (PSM) count. The identified PSMs, peptides, and proteins were quantified using TMT reporter ions in the MS2 scans ^34^.

To correct for ratio compression in reporter-based quantification in the MS2 scans, we used fully SILAC-labeled mouse tissues, generated over two generations ^53^, as negative controls to detect noise signals. For light Lys peptides, the noise detection process involved the following steps: (i) experimental TMT ions were extracted for each PSM; (ii) the abundance in the channel of the fully labeled SILAC tissues was considered noise; (iii) this noise was subtracted from all TMT channels; and (iv) PSM data were summarized into peptide and protein data. For heavy Lys peptides, we applied a similar approach, using the channel of unlabeled mouse tissues as a negative control to detect and remove noise.

#### Computational procedure to obtain free lysine decay curve by double-K-peptides

During the Lys-based pSILAC analysis of animals, the labeling process is significantly affected by free Lys recycled from protein degradation. Therefore, it is critical to obtain the free Lys decay curve during the labeling process, which is used as input in the JUMPt program. In BoxCarmax-DIA, DIA-MS, and TMTpro results, a small portion of the peptides contained two Lys residues (i.e., double-K-peptides). These double-K-peptides were used to derive the Lys decay curve.

Briefly, the double-K-peptides should exhibit three peaks: light, mixed, and heavy. If each protein (P) has a synthesis rate (S) following zero-order kinetics, and H_A_% represents the average percentage of heavy amino acid (e.g. K) over time (t), we can derive the following:

***P_M_* (mixed) = *S*** × ***t*** × ***H_A_%*** × ***(1-H_A_%)*** × ***2*** (as two Lys have an equal probability of being heavy)

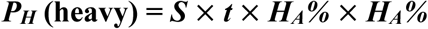

The ratio of the mixed peptide to the heavy peptide 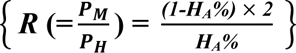 was independent of synthesis rates. Thus,

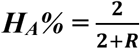

***L_A_%*** (the average percentage of light amino acid) ***= 1 - H_A_%***

Using these equations, we can derive the average LA% during the pulse (e.g., eight days) from double-K-peptides.

#### Mouse Tissue Protein and Phosphopeptide Half-life Analysis

For both BoxCarmax-DIA and TMTpro results, we used the JUMPt pipeline ^16^ to calculate protein and phosphopeptide half-lives, utilizing setting 2, which incorporates the free Lys decay curve and protein turnover data to fit an ordinary differential equation-based model to determine protein degradation rates.

For the final data analysis, which is included in the Tissue-PPT Web, both BoxCarmax-DIA and TMT data were further filtered and combined based on the following rules: (i) for proteins with a half-life of at least 0.5 days, the multiplex-DIA BoxCarmax results were used as the primary data, supplemented by TMT method results; (ii) for proteins with a half-life of less than 0.5 days (0.1-0.5% of the total results), DIA data with a half-life CV of less than 0.3 and TMT results were averaged to generate the results.

#### Kinase Substrate Mapping and Annotation

For kinase-substrate mapping, the mouse phosphopeptide sequence was analyzed using the PTMoreR (https://github.com/wangshisheng/PTMoreR)^114^ and MotifeR^115^ (https://github.com/wangshisheng/motifeR) software, which aligned the mouse phosphopeptide with human sequences with a 14-amino-acid window (Window similarity score > 14, full match). Additonally, the resulting human sequences was used to retrieve kinase substrates from the kinase library ^77,78^ (https://kinase-library.mit.edu/sites) with the percentile threshold 99%.

#### Ubiquitin Linkage Identification and Turnover Quantification

To search and determine the different type of ubiquitin chains via its lysine residue (i.e., the “ubiquitin code’), a “Gly-Gly” or diGly modification was set up as a variable modification in a separated search and summarized, as previously described ^47^. The DIA raw data was directly searched by setting K-GlyGly as variable modification in Spectronaut with the PTM score > 0.75. The quantitative data from Spectronaut was reported as described above for the phosphoproteomics analysis and ISW workflow was also used ^67^. The search results were further manually inspected. The quantities of K6-, K11-, K27-, K48-, and K63-were inferred based on most abundant peptide precursors for MQIFVK_GG_TLTGK (K6), TLTGK_GG_TITLEVEPSDTIENVK (K11), TITLEVEPSDTIENVK_GG_AK (K27), LIFAGK_GG_QLEDGR (K48), and TLSDYNIQK_GG_ESTLHLVLR (K63), respectively. To accurately determine the relative quantitative variability between K6-, K11-, K27-, K48-, and K63-linked chains, the above peptides carrying diGly were compared to the adjacent unmodified counterpart peptide with no miss-cleavage (TLSDYNIQK was used for K48 and K63, and TITLEVEPSDTIENVK was used for K11, K27 and K6). The heavy-to-light ratio of ubiquitin diGly peptide LIFAGK_GG_QLEDGR (K48), and TLSDYNIQK_GG_ESTLHLVLR (K63) and unmodified ubiquitin peptide TLSDYNIQK were also exported from Spectronaut to determine the respective turnover kinetics.

#### Protein-protein Interaction Mapping and Comparison

Databases of hu.MAP ^65^ (http://humap2.proteincomplexes.org/), CORUM ^61^ (https://mips.helmholtz-muenchen.de/corum/), Bioplex 3.0 ^62^ (https://bioplex.hms.harvard.edu/interactions.php), and mouse protein-protein interaction discovery (by Skinnider et al. ^60^) were downloaded and compiled separately for PPI mapping based on gene symbols. Next, respective Pearson correlations between the proteins participating the database-matched PPI pairs were calculated based on PA and PT levels measured across mouse tissues in our results. The hu.MAP confidence levels (Level 5-1 indicating Extremely High, Very High, High, Medium High, and Medium) were additionally used to group the PPI pairs for comparing the Pearson correlations of PA and PT across tissues between PPIs.

To evaluate the predictive power of PPI based on PA and PT’s cross-tissue correlations and existence of PPI in CORUM and Bioplex, Receiver Operating Characteristic (ROC) curves were generated with the corresponding Area Under the Curve (AUC) computed ^116^. To enable a fair comparison, the “positive” PPI pairs were retrieved from the Extremely High and High Levels in hu.MAP database. The same number of “false” PPIs was randomly generated from any protein pairs excluding those pairs listed in any of hu.MAP, CORUM, Bioplex 3.0 and Skinnider et al. The logistic regression model was employed to evaluate the combined predictive power of PA and PT using Scikit-Learn (Python)^117^, employing default parameters unless otherwise specified.

#### mRNA-seq and Ribo-seq Data in Multiple Tissues

The mRNA-seq and Ribo-seq data (i.e., the translatome) of adult wild-type C57BL/6 mouse tissues were downloaded from a published paper via NCBI Gene Expression Omnibus under accession number GSE94982 (https://www.ncbi.nlm.nih.gov/geo/query/acc.cgi?acc=GSE94982: GSE94982_P42_RNA-seq_exon_level_tpm.txt.gz and GSE94982_P42_Ribo-seq_exon_level_tpm.txt.gz). All Ribo-seq and RNA-seq samples were combined and the transcripts per kilobase million (TPM) values >1 was used for mapping to the proteomic data.

#### Other Bioinformatics Analysis

Most data visualization was performed in R and GraphPad Prism version 10 (GraphPad Software, San Diego, California USA). The following R packages were used to visualize the data: ‘ggplot2’ (boxplots, density plots, volcano plots and histograms), ‘factoextra’ (principal component analysis [PCA]), ‘pheatmap’ (heatmaps), ‘corrplot’ (correlation plots) and ‘UpsetR’ (UpSet plots). The Cytoscape v3.10.1 ^118^ was used for the PPI plots. The Perseus software^119^ was used for the protein Gene Oncology Cellular Component (GO CC) and Gene Oncology Biological Process (GO BP) annotation with the mouse species. The 2D enrichment function^120^ was used to generate the data for the bubble plots. The protein-level functional annotation was also performed using Metascape (https://metascape.org/) ^121^. The lists of E3 ubiquitin ligases, E3 ubiquitin ligase accessory proteins, and De-ubiquitination enzymes (DUBs) were downloaded from NIH (https://esbl.nhlbi.nih.gov/Databases/KSBP2/Targets/Lists). The list of molecular chaperons was downloaded from literature ^122^. BioMart (https://useast.ensembl.org/info/data/biomart/index.html) was used to map the gene symbols between human and mouse species. Figure 1A and 2E were generated with the assistance of BioRender.

#### Tissue-PPT Website Inventory

The website of Tissue-PPT (https://yslproteomics.shinyapps.io/tissuePPT/) was created by Shiny framework (version 1.9.1) in R environment (version 4.4.1) and deployed on the shinyapps.io platform (https://www.shinyapps.io/) to facilitate navigation of the database (**Figure S3**). This website interactively provides queries about protein/phosphosite abundance and lifetime in various tissues. It offers four major functions (a) including Heat-circle (HC) plot across tissues, (b) Heatmap analysis for protein sets, (c) Protein-specific barplots, and (d) Correlation analysis between molecular layers for individual proteins or protein sets of interest, as well as convenient options to download all the resultant figures and tables.

#### Data Availability

The mass spectrometry raw data and searched results have been all deposited to the ProteomeXchange Consortium via the PRIDE ^123^.

## REFERENCE

1. Glass, R.D., and Doyle, D. (1972). On the measurement of protein turnover in animal cells. J Biol Chem 247, 5234–5242.

2. Pratt, J.M., Petty, J., Riba-Garcia, I., Robertson, D.H., Gaskell, S.J., Oliver, S.G., and Beynon, R.J. (2002). Dynamics of protein turnover, a missing dimension in proteomics. Molecular & cellular proteomics : MCP 1, 579–591.

3. Claydon, A.J., and Beynon, R. (2012). Proteome dynamics: revisiting turnover with a global perspective. Molecular & cellular proteomics : MCP 11, 1551–1565. 10.1074/mcp.O112.022186.

4. Ross, A.B., Langer, J.D., and Jovanovic, M. (2021). Proteome Turnover in the Spotlight: Approaches, Applications, and Perspectives. Molecular & cellular proteomics : MCP 20, 100016. 10.1074/mcp.R120.002190.

5. Liu, Y., Borel, C., Li, L., Muller, T., Williams, E.G., Germain, P.L., Buljan, M., Sajic, T., Boersema, P.J., Shao, W., et al. (2017). Systematic proteome and proteostasis profiling in human Trisomy 21 fibroblast cells. Nature communications 8, 1212. 10.1038/s41467-017-01422-6.

6. Liu, Y., Beyer, A., and Aebersold, R. (2016). On the Dependency of Cellular Protein Levels on mRNA Abundance. Cell 165, 535–550. 10.1016/j.cell.2016.03.014.

7. Fornasiero, E.F., and Savas, J.N. (2023). Determining and interpreting protein lifetimes in mammalian tissues. Trends Biochem Sci 48, 106–118. 10.1016/j.tibs.2022.08.011.

8. McClatchy, D.B., Dong, M.Q., Wu, C.C., Venable, J.D., and Yates, J.R., 3rd (2007). 15N metabolic labeling of mammalian tissue with slow protein turnover. Journal of proteome research 6, 2005–2010. 10.1021/pr060599n.

9. Fornasiero, E.F., Mandad, S., Wildhagen, H., Alevra, M., Rammner, B., Keihani, S., Opazo, F., Urban, I., Ischebeck, T., Sakib, M.S., et al. (2018). Precisely measured protein lifetimes in the mouse brain reveal differences across tissues and subcellular fractions. Nature communications 9, 4230. 10.1038/s41467-018-06519-0.

10. Alevra, M., Mandad, S., Ischebeck, T., Urlaub, H., Rizzoli, S.O., and Fornasiero, E.F. (2019). A mass spectrometry workflow for measuring protein turnover rates in vivo. Nature protocols 14, 3333–3365. 10.1038/s41596-019-0222-y.

11. Rolfs, Z., Frey, B.L., Shi, X., Kawai, Y., Smith, L.M., and Welham, N.V. (2021). An atlas of protein turnover rates in mouse tissues. Nature communications 12, 6778. 10.1038/s41467-021-26842-3.

12. Kluever, V., Russo, B., Mandad, S., Kumar, N.H., Alevra, M., Ori, A., Rizzoli, S.O., Urlaub, H., Schneider, A., and Fornasiero, E.F. (2022). Protein lifetimes in aged brains reveal a proteostatic adaptation linking physiological aging to neurodegeneration. Sci Adv 8, eabn4437. 10.1126/sciadv.abn4437.

13. Hasper, J., Welle, K., Hryhorenko, J., Ghaemmaghami, S., and Buchwalter, A. (2023). Turnover and replication analysis by isotope labeling (TRAIL) reveals the influence of tissue context on protein and organelle lifetimes. Molecular systems biology 19, e11393. 10.15252/msb.202211393.

14. Rao, N.R., Upadhyay, A., and Savas, J.N. (2024). Derailed protein turnover in the aging mammalian brain. Molecular systems biology 20, 120–139. 10.1038/s44320-023-00009-2.

15. Schwanhausser, B., Busse, D., Li, N., Dittmar, G., Schuchhardt, J., Wolf, J., Chen, W., and Selbach, M. (2011). Global quantification of mammalian gene expression control. Nature 473, 337–342. 10.1038/nature10098.

16. Chepyala, S.R., Liu, X., Yang, K., Wu, Z., Breuer, A.M., Cho, J.H., Li, Y., Mancieri, A., Jiao, Y., Zhang, H., and Peng, J. (2021). JUMPt: Comprehensive Protein Turnover Modeling of In Vivo Pulse SILAC Data by Ordinary Differential Equations. Analytical chemistry 93, 13495–13504. 10.1021/acs.analchem.1c02309.

17. Giansanti, P., Samaras, P., Bian, Y., Meng, C., Coluccio, A., Frejno, M., Jakubowsky, H., Dobiasch, S., Hazarika, R.R., Rechenberger, J., et al. (2022). Mass spectrometry-based draft of the mouse proteome. Nature methods 19, 803–811. 10.1038/s41592-022-01526-y.

18. Lu, T., Qian, L., Xie, Y., Zhang, Q., Liu, W., Ge, W., Zhu, Y., Ma, L., Zhang, C., and Guo, T. (2022). Tissue-Characteristic Expression of Mouse Proteome. Molecular & cellular proteomics : MCP 21, 100408. 10.1016/j.mcpro.2022.100408.

19. Aebersold, R., and Mann, M. (2016). Mass-spectrometric exploration of proteome structure and function. Nature 537, 347–355. 10.1038/nature19949.

20. Liu, Y. (2022). A peptidoform based proteomic strategy for studying functions of post-translational modifications. Proteomics 22, e2100316. 10.1002/pmic.202100316.

21. Huttlin, E.L., Jedrychowski, M.P., Elias, J.E., Goswami, T., Rad, R., Beausoleil, S.A., Villen, J., Haas, W., Sowa, M.E., and Gygi, S.P. (2010). A tissue-specific atlas of mouse protein phosphorylation and expression. Cell 143, 1174–1189. 10.1016/j.cell.2010.12.001.

22. Wu, C., Ba, Q., Lu, D., Li, W., Salovska, B., Hou, P., Mueller, T., Rosenberger, G., Gao, E., Di, Y., et al. (2021). Global and Site-Specific Effect of Phosphorylation on Protein Turnover. Developmental cell 56, 111–124 e116. 10.1016/j.devcel.2020.10.025.

23. Zecha, J., Gabriel, W., Spallek, R., Chang, Y.C., Mergner, J., Wilhelm, M., Bassermann, F., and Kuster, B. (2022). Linking post-translational modifications and protein turnover by site-resolved protein turnover profiling. Nature communications 13, 165. 10.1038/s41467-021-27639-0.

24. Hammaren, H.M., Geissen, E.M., Potel, C.M., Beck, M., and Savitski, M.M. (2022). Protein-Peptide Turnover Profiling reveals the order of PTM addition and removal during protein maturation. Nature communications 13, 7431. 10.1038/s41467-022-35054-2.

25. Li, W., Salovska, B., Fornasiero, E.F., and Liu, Y. (2022). Toward a hypothesis-free understanding of how phosphorylation dynamically impacts protein turnover. Proteomics, e2100387. 10.1002/pmic.202100387.

26. Venable, J.D., Dong, M.Q., Wohlschlegel, J., Dillin, A., and Yates, J.R. (2004). Automated approach for quantitative analysis of complex peptide mixtures from tandem mass spectra. Nature methods 1, 39–45. 10.1038/nmeth705.

27. Gillet, L.C., Navarro, P., Tate, S., Rost, H., Selevsek, N., Reiter, L., Bonner, R., and Aebersold, R. (2012). Targeted data extraction of the MS/MS spectra generated by data-independent acquisition: a new concept for consistent and accurate proteome analysis. Molecular & cellular proteomics : MCP 11, O111 016717. 10.1074/mcp.O111.016717.

28. Gao, E., Li, W., Wu, C., Shao, W., Di, Y., and Liu, Y. (2021). Data-independent acquisition-based proteome and phosphoproteome profiling across six melanoma cell lines reveals determinants of proteotypes. Mol Omics 17, 413–425. 10.1039/d0mo00188k.

29. Li, J., Van Vranken, J.G., Pontano Vaites, L., Schweppe, D.K., Huttlin, E.L., Etienne, C., Nandhikonda, P., Viner, R., Robitaille, A.M., Thompson, A.H., et al. (2020). TMTpro reagents: a set of isobaric labeling mass tags enables simultaneous proteome-wide measurements across 16 samples. Nature methods. 10.1038/s41592-020-0781-4.

30. Liu, D., Yang, S., Kavdia, K., Sifford, J.M., Wu, Z., Xie, B., Wang, Z., Pagala, V.R., Wang, H., Yu, K., et al. (2021). Deep Profiling of Microgram-Scale Proteome by Tandem Mass Tag Mass Spectrometry. Journal of proteome research 20, 337–345. 10.1021/acs.jproteome.0c00426.

31. Kruger, M., Moser, M., Ussar, S., Thievessen, I., Luber, C.A., Forner, F., Schmidt, S., Zanivan, S., Fassler, R., and Mann, M. (2008). SILAC mouse for quantitative proteomics uncovers kindlin-3 as an essential factor for red blood cell function. Cell 134, 353–364. 10.1016/j.cell.2008.05.033.

32. Salovska, B., Li, W., Di, Y., and Liu, Y. (2021). BoxCarmax: A High-Selectivity Data-Independent Acquisition Mass Spectrometry Method for the Analysis of Protein Turnover and Complex Samples. Analytical chemistry 93, 3103–3111. 10.1021/acs.analchem.0c04293.

33. Yu, K., Wang, Z., Wu, Z., Tan, H., Mishra, A., and Peng, J. (2021). High-Throughput Profiling of Proteome and Posttranslational Modifications by 16-Plex TMT Labeling and Mass Spectrometry. Methods Mol Biol 2228, 205–224. 10.1007/978-1-0716-1024-4_15.

34. Niu, M., Cho, J.H., Kodali, K., Pagala, V., High, A.A., Wang, H., Wu, Z., Li, Y., Bi, W., Zhang, H., et al. (2017). Extensive Peptide Fractionation and y(1) Ion-Based Interference Detection Method for Enabling Accurate Quantification by Isobaric Labeling and Mass Spectrometry. Analytical chemistry 89, 2956–2963. 10.1021/acs.analchem.6b04415.

35. Lau, E., Cao, Q., Lam, M.P.Y., Wang, J., Ng, D.C.M., Bleakley, B.J., Lee, J.M., Liem, D.A., Wang, D., Hermjakob, H., and Ping, P. (2018). Integrated omics dissection of proteome dynamics during cardiac remodeling. Nature communications 9, 120. 10.1038/s41467-017-02467-3.

36. Sato, C., Barthelemy, N.R., Mawuenyega, K.G., Patterson, B.W., Gordon, B.A., Jockel-Balsarotti, J., Sullivan, M., Crisp, M.J., Kasten, T., Kirmess, K.M., et al. (2018). Tau Kinetics in Neurons and the Human Central Nervous System. Neuron 97, 1284–1298 e1287. 10.1016/j.neuron.2018.02.015.

37. Zhang, Y., Wu, K.M., Yang, L., Dong, Q., and Yu, J.T. (2022). Tauopathies: new perspectives and challenges. Mol Neurodegener 17, 28. 10.1186/s13024-022-00533-z.

38. Li, Q., Chang, Z., Oliveira, G., Xiong, M., Smith, L.M., Frey, B.L., and Welham, N.V. (2016). Protein turnover during in vitro tissue engineering. Biomaterials 81, 104–113. 10.1016/j.biomaterials.2015.12.004.

39. Di Camillo, B., Puricelli, L., Iori, E., Toffolo, G.M., Tessari, P., and Arrigoni, G. (2023). Modeling SILAC Data to Assess Protein Turnover in a Cellular Model of Diabetic Nephropathy. Int J Mol Sci 24. 10.3390/ijms24032811.

40. Polo, S.E., and Jackson, S.P. (2011). Dynamics of DNA damage response proteins at DNA breaks: a focus on protein modifications. Genes Dev 25, 409–433. 10.1101/gad.2021311.

41. Trulsson, F., Akimov, V., Robu, M., van Overbeek, N., Berrocal, D.A.P., Shah, R.G., Cox, J., Shah, G.M., Blagoev, B., and Vertegaal, A.C.O. (2022). Deubiquitinating enzymes and the proteasome regulate preferential sets of ubiquitin substrates. Nature communications 13, 2736. 10.1038/s41467-022-30376-7.

42. Prus, G., Satpathy, S., Weinert, B.T., Narita, T., and Choudhary, C. (2024). Global, site-resolved analysis of ubiquitylation occupancy and turnover rate reveals systems properties. Cell 187, 2875–2892 e2821. 10.1016/j.cell.2024.03.024.

43. Wilhelm, M., Schlegl, J., Hahne, H., Gholami, A.M., Lieberenz, M., Savitski, M.M., Ziegler, E., Butzmann, L., Gessulat, S., Marx, H., et al. (2014). Mass-spectrometry-based draft of the human proteome. Nature 509, 582–587. 10.1038/nature13319.

44. Bruderer, R., Bernhardt, O.M., Gandhi, T., Xuan, Y., Sondermann, J., Schmidt, M., Gomez-Varela, D., and Reiter, L. (2017). Optimization of Experimental Parameters in Data-Independent Mass Spectrometry Significantly Increases Depth and Reproducibility of Results. Molecular & cellular proteomics : MCP 16, 2296–2309. 10.1074/mcp.RA117.000314.

45. Toyama, B.H., Savas, J.N., Park, S.K., Harris, M.S., Ingolia, N.T., Yates, J.R., 3rd, and Hetzer, M.W. (2013). Identification of long-lived proteins reveals exceptional stability of essential cellular structures. Cell 154, 971–982. 10.1016/j.cell.2013.07.037.

46. Verzijl, N., DeGroot, J., Thorpe, S.R., Bank, R.A., Shaw, J.N., Lyons, T.J., Bijlsma, J.W., Lafeber, F.P., Baynes, J.W., and TeKoppele, J.M. (2000). Effect of collagen turnover on the accumulation of advanced glycation end products. J Biol Chem 275, 39027–39031. 10.1074/jbc.M006700200.

47. Ba, Q., Hei, Y., Dighe, A., Li, W., Maziarz, J., Pak, I., Wang, S., Wagner, G.P., and Liu, Y. (2022). Proteotype coevolution and quantitative diversity across 11 mammalian species. Sci Adv 8, eabn0756. 10.1126/sciadv.abn0756.

48. Nguyen, J.A., and Yates, R.M. (2021). Better Together: Current Insights Into Phagosome-Lysosome Fusion. Front Immunol 12, 636078. 10.3389/fimmu.2021.636078.

49. Zuehlke, A.D., Beebe, K., Neckers, L., and Prince, T. (2015). Regulation and function of the human HSP90AA1 gene. Gene 570, 8–16. 10.1016/j.gene.2015.06.018.

50. Xu, P., Duong, D.M., Seyfried, N.T., Cheng, D., Xie, Y., Robert, J., Rush, J., Hochstrasser, M., Finley, D., and Peng, J. (2009). Quantitative proteomics reveals the function of unconventional ubiquitin chains in proteasomal degradation. Cell 137, 133–145. 10.1016/j.cell.2009.01.041.

51. Kim, W., Bennett, E.J., Huttlin, E.L., Guo, A., Li, J., Possemato, A., Sowa, M.E., Rad, R., Rush, J., Comb, M.J., et al. (2011). Systematic and quantitative assessment of the ubiquitin-modified proteome. Molecular cell 44, 325–340. 10.1016/j.molcel.2011.08.025.

52. Tracz, M., and Bialek, W. (2021). Beyond K48 and K63: non-canonical protein ubiquitination. Cell Mol Biol Lett 26, 1. 10.1186/s11658-020-00245-6.

53. Dammer, E.B., Na, C.H., Xu, P., Seyfried, N.T., Duong, D.M., Cheng, D., Gearing, M., Rees, H., Lah, J.J., Levey, A.I., et al. (2011). Polyubiquitin linkage profiles in three models of proteolytic stress suggest the etiology of Alzheimer disease. J Biol Chem 286, 10457–10465. 10.1074/jbc.M110.149633.

54. Verma, R., Aravind, L., Oania, R., McDonald, W.H., Yates, J.R., 3rd, Koonin, E.V., and Deshaies, R.J. (2002). Role of Rpn11 metalloprotease in deubiquitination and degradation by the 26S proteasome. Science 298, 611–615. 10.1126/science.1075898.

55. Kudriaeva, A.A., Livneh, I., Baranov, M.S., Ziganshin, R.H., Tupikin, A.E., Zaitseva, S.O., Kabilov, M.R., Ciechanover, A., and Belogurov, A.A., Jr. (2021). In-depth characterization of ubiquitin turnover in mammalian cells by fluorescence tracking. Cell Chem Biol 28, 1192–1205 e1199. 10.1016/j.chembiol.2021.02.009.

56. Erpapazoglou, Z., Walker, O., and Haguenauer-Tsapis, R. (2014). Versatile roles of k63-linked ubiquitin chains in trafficking. Cells 3, 1027–1088. 10.3390/cells3041027.

57. Waltho, A., Popp, O., Lenz, C., Pluska, L., Lambert, M., Dotsch, V., Mertins, P., and Sommer, T. (2024). K48- and K63-linked ubiquitin chain interactome reveals branch- and length-specific ubiquitin interactors. Life Sci Alliance 7. 10.26508/lsa.202402740.

58. Currie, J., Manda, V., Robinson, S.K., Lai, C., Agnihotri, V., Hidalgo, V., Ludwig, R.W., Zhang, K., Pavelka, J., Wang, Z.V., et al. (2024). Simultaneous proteome localization and turnover analysis reveals spatiotemporal features of protein homeostasis disruptions. bioRxiv. 10.1101/2023.01.04.521821.

59. Mathieson, T., Franken, H., Kosinski, J., Kurzawa, N., Zinn, N., Sweetman, G., Poeckel, D., Ratnu, V.S., Schramm, M., Becher, I., et al. (2018). Systematic analysis of protein turnover in primary cells. Nature communications 9, 689. 10.1038/s41467-018-03106-1.

60. Skinnider, M.A., Scott, N.E., Prudova, A., Kerr, C.H., Stoynov, N., Stacey, R.G., Chan, Q.W.T., Rattray, D., Gsponer, J., and Foster, L.J. (2021). An atlas of protein-protein interactions across mouse tissues. Cell 184, 4073–4089 e4017. 10.1016/j.cell.2021.06.003.

61. Giurgiu, M., Reinhard, J., Brauner, B., Dunger-Kaltenbach, I., Fobo, G., Frishman, G., Montrone, C., and Ruepp, A. (2019). CORUM: the comprehensive resource of mammalian protein complexes-2019. Nucleic Acids Res 47, D559–D563. 10.1093/nar/gky973.

62. Huttlin, E.L., Bruckner, R.J., Navarrete-Perea, J., Cannon, J.R., Baltier, K., Gebreab, F., Gygi, M.P., Thornock, A., Zarraga, G., Tam, S., et al. (2021). Dual proteome-scale networks reveal cell-specific remodeling of the human interactome. Cell 184, 3022–3040 e3028. 10.1016/j.cell.2021.04.011.

63. Srivastava, H., Lippincott, M.J., Currie, J., Canfield, R., Lam, M.P.Y., and Lau, E. (2022). Protein prediction models support widespread post-transcriptional regulation of protein abundance by interacting partners. PLoS computational biology 18, e1010702. 10.1371/journal.pcbi.1010702.

64. Sonawane, A.R., Platig, J., Fagny, M., Chen, C.Y., Paulson, J.N., Lopes-Ramos, C.M., DeMeo, D.L., Quackenbush, J., Glass, K., and Kuijjer, M.L. (2017). Understanding Tissue-Specific Gene Regulation. Cell reports 21, 1077–1088. 10.1016/j.celrep.2017.10.001.

65. Drew, K., Wallingford, J.B., and Marcotte, E.M. (2021). hu.MAP 2.0: integration of over 15,000 proteomic experiments builds a global compendium of human multiprotein assemblies. Molecular systems biology 17, e10016. 10.15252/msb.202010016.

66. Wang, H., Wang, Y., Yang, J., Zhao, Q., Tang, N., Chen, C., Li, H., Cheng, C., Xie, M., Yang, Y., and Xie, Z. (2021). Tissue- and stage-specific landscape of the mouse translatome. Nucleic Acids Res 49, 6165–6180. 10.1093/nar/gkab482.

67. Salovska, B., Zhu, H., Gandhi, T., Frank, M., Li, W., Rosenberger, G., Wu, C., Germain, P.L., Zhou, H., Hodny, Z., et al. (2020). Isoform-resolved correlation analysis between mRNA abundance regulation and protein level degradation. Molecular systems biology 16, e9170. 10.15252/msb.20199170.

68. Wang, D., Eraslan, B., Wieland, T., Hallstrom, B., Hopf, T., Zolg, D.P., Zecha, J., Asplund, A., Li, L.H., Meng, C., et al. (2019). A deep proteome and transcriptome abundance atlas of 29 healthy human tissues. Molecular systems biology 15, e8503. 10.15252/msb.20188503.

69. Romanov, N., Kuhn, M., Aebersold, R., Ori, A., Beck, M., and Bork, P. (2019). Disentangling Genetic and Environmental Effects on the Proteotypes of Individuals. Cell 177, 1308–1318 e1310. 10.1016/j.cell.2019.03.015.

70. Uhlen, M., Fagerberg, L., Hallstrom, B.M., Lindskog, C., Oksvold, P., Mardinoglu, A., Sivertsson, A., Kampf, C., Sjostedt, E., Asplund, A., et al. (2015). Proteomics. Tissue-based map of the human proteome. Science 347, 1260419. 10.1126/science.1260419.

71. Germain, K., and Kim, P.K. (2020). Pexophagy: A Model for Selective Autophagy. Int J Mol Sci 21. 10.3390/ijms21020578.

72. Islinger, M., Cardoso, M.J., and Schrader, M. (2010). Be different--the diversity of peroxisomes in the animal kingdom. Biochim Biophys Acta 1803, 881–897. 10.1016/j.bbamcr.2010.03.013.

73. Burnett, S.F., Farre, J.C., Nazarko, T.Y., and Subramani, S. (2015). Peroxisomal Pex3 activates selective autophagy of peroxisomes via interaction with the pexophagy receptor Atg30. J Biol Chem 290, 8623–8631. 10.1074/jbc.M114.619338.

74. Sakai, Y., Oku, M., van der Klei, I.J., and Kiel, J.A. (2006). Pexophagy: autophagic degradation of peroxisomes. Biochim Biophys Acta 1763, 1767–1775. 10.1016/j.bbamcr.2006.08.023.

75. Mizushima, N., Noda, T., Yoshimori, T., Tanaka, Y., Ishii, T., George, M.D., Klionsky, D.J., Ohsumi, M., and Ohsumi, Y. (1998). A protein conjugation system essential for autophagy. Nature 395, 395–398. 10.1038/26506.

76. Niu, L., Thiele, M., Geyer, P.E., Rasmussen, D.N., Webel, H.E., Santos, A., Gupta, R., Meier, F., Strauss, M., Kjaergaard, M., et al. (2022). Noninvasive proteomic biomarkers for alcohol-related liver disease. Nat Med 28, 1277–1287. 10.1038/s41591-022-01850-y.

77. Yaron-Barir, T.M., Joughin, B.A., Huntsman, E.M., Kerelsky, A., Cizin, D.M., Cohen, B.M., Regev, A., Song, J., Vasan, N., Lin, T.Y., et al. (2024). The intrinsic substrate specificity of the human tyrosine kinome. Nature 629, 1174–1181. 10.1038/s41586-024-07407-y.

78. Johnson, J.L., Yaron, T.M., Huntsman, E.M., Kerelsky, A., Song, J., Regev, A., Lin, T.Y., Liberatore, K., Cizin, D.M., Cohen, B.M., et al. (2023). An atlas of substrate specificities for the human serine/threonine kinome. Nature 613, 759–766. 10.1038/s41586-022-05575-3.

79. Savas, J.N., Toyama, B.H., Xu, T., Yates, J.R., 3rd, and Hetzer, M.W. (2012). Extremely long-lived nuclear pore proteins in the rat brain. Science 335, 942. 10.1126/science.1217421.

80. Alonso, A., Zaidi, T., Novak, M., Grundke-Iqbal, I., and Iqbal, K. (2001). Hyperphosphorylation induces self-assembly of tau into tangles of paired helical filaments/straight filaments. Proceedings of the National Academy of Sciences of the United States of America 98, 6923–6928. 10.1073/pnas.121119298.

81. Dickey, C.A., Kamal, A., Lundgren, K., Klosak, N., Bailey, R.M., Dunmore, J., Ash, P., Shoraka, S., Zlatkovic, J., Eckman, C.B., et al. (2007). The high-affinity HSP90-CHIP complex recognizes and selectively degrades phosphorylated tau client proteins. J Clin Invest 117, 648–658. 10.1172/JCI29715.

82. Kawahata, I., Finkelstein, D.I., and Fukunaga, K. (2022). Pathogenic Impact of alpha-Synuclein Phosphorylation and Its Kinases in alpha-Synucleinopathies. Int J Mol Sci 23. 10.3390/ijms23116216.

83. Hu, Z., Chen, P.H., Li, W., Douglas, T., Hines, J., Liu, Y., and Crews, C.M. (2023). Targeted Dephosphorylation of Tau by Phosphorylation Targeting Chimeras (PhosTACs) as a Therapeutic Modality. J Am Chem Soc. 10.1021/jacs.2c11706.

84. Hu, Z., Chen, P.H., Li, W., Krone, M., Zheng, S., Saarbach, J., Velasco, I.U., Hines, J., Liu, Y., and Crews, C.M. (2024). EGFR targeting PhosTACs as a dual inhibitory approach reveals differential downstream signaling. Sci Adv 10, eadj7251. 10.1126/sciadv.adj7251.

85. Wang, S., Li, W., Hu, L., Cheng, J., Yang, H., and Liu, Y. (2020). NAguideR: performing and prioritizing missing value imputations for consistent bottom-up proteomic analyses. Nucleic Acids Res. 10.1093/nar/gkaa498.

86. Guan, S., Price, J.C., Ghaemmaghami, S., Prusiner, S.B., and Burlingame, A.L. (2012). Compartment modeling for mammalian protein turnover studies by stable isotope metabolic labeling. Analytical chemistry 84, 4014–4021. 10.1021/ac203330z.

87. Ochoa, D., Jarnuczak, A.F., Vieitez, C., Gehre, M., Soucheray, M., Mateus, A., Kleefeldt, A.A., Hill, A., Garcia-Alonso, L., Stein, F., et al. (2020). The functional landscape of the human phosphoproteome. Nature biotechnology 38, 365–373. 10.1038/s41587-019-0344-3.

88. Mihailovich, M., Germain, P.L., Shyti, R., Pozzi, D., Noberini, R., Liu, Y., Aprile, D., Tenderini, E., Troglio, F., Trattaro, S., et al. (2024). Multiscale modeling uncovers 7q11.23 copy number variation-dependent changes in ribosomal biogenesis and neuronal maturation and excitability. J Clin Invest 134. 10.1172/JCI168982.

89. Liu, Y., Yang, J., Wang, T., Luo, M., Chen, Y., Chen, C., Ronai, Z., Zhou, Y., Ruppin, E., and Han, L. (2023). Expanding PROTACtable genome universe of E3 ligases. Nature communications 14, 6509. 10.1038/s41467-023-42233-2.

90. Rahman, M., Previs, S.F., Kasumov, T., and Sadygov, R.G. (2016). Gaussian Process Modeling of Protein Turnover. Journal of proteome research 15, 2115–2122. 10.1021/acs.jproteome.5b00990.

91. Ding, C., Li, Y., Guo, F., Jiang, Y., Ying, W., Li, D., Yang, D., Xia, X., Liu, W., Zhao, Y., et al. (2016). A Cell-type-resolved Liver Proteome. Molecular & cellular proteomics : MCP 15, 3190–3202. 10.1074/mcp.M116.060145.

92. Nusinow, D.P., Szpyt, J., Ghandi, M., Rose, C.M., McDonald, E.R., 3rd, Kalocsay, M., Jane-Valbuena, J., Gelfand, E., Schweppe, D.K., Jedrychowski, M., et al. (2020). Quantitative Proteomics of the Cancer Cell Line Encyclopedia. Cell 180, 387–402 e316. 10.1016/j.cell.2019.12.023.

93. Kim-Hellmuth, S., Aguet, F., Oliva, M., Munoz-Aguirre, M., Kasela, S., Wucher, V., Castel, S.E., Hamel, A.R., Vinuela, A., Roberts, A.L., et al. (2020). Cell type-specific genetic regulation of gene expression across human tissues. Science 369. 10.1126/science.aaz8528.

94. Sabatier, P., Ye, Z., Lechner, M., Guzmán, U.H., Beusch, C.M., Izaguirre, F., Seth, A., Gritsenko, O., Rodin, S., Grinnemo, K.-H., and Olsen, J.V. (2024). Global analysis of protein turnover dynamics in single cells. bioRxiv, 2024.2005.2030.596745. 10.1101/2024.05.30.596745.

95. Bai, B., Tan, H., Pagala, V.R., High, A.A., Ichhaporia, V.P., Hendershot, L., and Peng, J. (2017). Deep Profiling of Proteome and Phosphoproteome by Isobaric Labeling, Extensive Liquid Chromatography, and Mass Spectrometry. Methods Enzymol 585, 377–395. 10.1016/bs.mie.2016.10.007.

96. Wang, Z., Yu, K., Tan, H., Wu, Z., Cho, J.H., Han, X., Sun, H., Beach, T.G., and Peng, J. (2020). 27-Plex Tandem Mass Tag Mass Spectrometry for Profiling Brain Proteome in Alzheimer’s Disease. Anal Chem 92, 7162–7170. 10.1021/acs.analchem.0c00655.

97. Gao, Q., Zhu, H., Dong, L., Shi, W., Chen, R., Song, Z., Huang, C., Li, J., Dong, X., Zhou, Y., et al. (2019). Integrated Proteogenomic Characterization of HBV-Related Hepatocellular Carcinoma. Cell 179, 561–577 e522. 10.1016/j.cell.2019.08.052.

98. Mehnert, M., Li, W., Wu, C., Salovska, B., and Liu, Y. (2019). Combining Rapid Data Independent Acquisition and CRISPR Gene Deletion for Studying Potential Protein Functions: A Case of HMGN1. Proteomics, e1800438. 10.1002/pmic.201800438.

99. Li, W., Chi, H., Salovska, B., Wu, C., Sun, L., Rosenberger, G., and Liu, Y. (2019). Assessing the Relationship Between Mass Window Width and Retention Time Scheduling on Protein Coverage for Data-Independent Acquisition. J Am Soc Mass Spectrom. 10.1007/s13361-019-02243-1.

100. Salovska, B., Gao, E., Muller-Dott, S., Li, W., Cordon, C.C., Wang, S., Dugourd, A., Rosenberger, G., Saez-Rodriguez, J., and Liu, Y. (2023). Phosphoproteomic analysis of metformin signaling in colorectal cancer cells elucidates mechanism of action and potential therapeutic opportunities. Clin Transl Med 13, e1179. 10.1002/ctm2.1179.

101. Wang, H., Yang, Y., Li, Y., Bai, B., Wang, X., Tan, H., Liu, T., Beach, T.G., Peng, J., and Wu, Z. (2015). Systematic optimization of long gradient chromatography mass spectrometry for deep analysis of brain proteome. Journal of proteome research 14, 829–838. 10.1021/pr500882h.

102. Keihani, S., Kluever, V., Mandad, S., Bansal, V., Rahman, R., Fritsch, E., Gomes, L.C., Gärtner, A., Kügler, S., Urlaub, H., et al. (2019). The long noncoding RNA neuroLNC regulates presynaptic activity by interacting with the neurodegeneration-associated protein TDP-43. Sci Adv 5, eaay2670. 10.1126/sciadv.aay2670.

103. Kaech, S., and Banker, G. (2006). Culturing hippocampal neurons. Nat Protoc 1, 2406–2415. 10.1038/nprot.2006.356.

104. Truckenbrodt, S., Viplav, A., Jähne, S., Vogts, A., Denker, A., Wildhagen, H., Fornasiero, E.F., and Rizzoli, S.O. (2018). Newly produced synaptic vesicle proteins are preferentially used in synaptic transmission. Embo j 37. 10.15252/embj.201798044.

105. Yousefi, R., Jevdokimenko, K., Kluever, V., Pacheu-Grau, D., and Fornasiero, E.F. (2021). Influence of Subcellular Localization and Functional State on Protein Turnover. Cells 10. 10.3390/cells10071747.

106. Collins, B.C., Hunter, C.L., Liu, Y., Schilling, B., Rosenberger, G., Bader, S.L., Chan, D.W., Gibson, B.W., Gingras, A.C., Held, J.M., et al. (2017). Multi-laboratory assessment of reproducibility, qualitative and quantitative performance of SWATH-mass spectrometry. Nature communications 8, 291. 10.1038/s41467-017-00249-5.

107. Bruderer, R., Bernhardt, O.M., Gandhi, T., Miladinovic, S.M., Cheng, L.Y., Messner, S., Ehrenberger, T., Zanotelli, V., Butscheid, Y., Escher, C., et al. (2015). Extending the limits of quantitative proteome profiling with data-independent acquisition and application to acetaminophen-treated three-dimensional liver microtissues. Molecular & cellular proteomics : MCP 14, 1400–1410. 10.1074/mcp.M114.044305.

108. Bekker-Jensen, D.B., Bernhardt, O.M., Hogrebe, A., Martinez-Val, A., Verbeke, L., Gandhi, T., Kelstrup, C.D., Reiter, L., and Olsen, J.V. (2020). Rapid and site-specific deep phosphoproteome profiling by data-independent acquisition without the need for spectral libraries. Nature communications 11, 787. 10.1038/s41467-020-14609-1.

109. Cox, J., and Mann, M. (2008). MaxQuant enables high peptide identification rates, individualized p.p.b.-range mass accuracies and proteome-wide protein quantification. Nat Biotechnol 26, 1367–1372. 10.1038/nbt.1511.

110. Olsen, J.V., Blagoev, B., Gnad, F., Macek, B., Kumar, C., Mortensen, P., and Mann, M. (2006). Global, in vivo, and site-specific phosphorylation dynamics in signaling networks. Cell 127, 635–648. 10.1016/j.cell.2006.09.026.

111. Zhou, W., Li, W., Wang, S., Salovska, B., Hu, Z., Tao, B., Di, Y., Punyamurtula, U., Turk, B.E., Sessa, W.C., and Liu, Y. (2023). An optogenetic-phosphoproteomic study reveals dynamic Akt1 signaling profiles in endothelial cells. Nature communications 14, 3803. 10.1038/s41467-023-39514-1.

112. Wang, X., Li, Y., Wu, Z., Wang, H., Tan, H., and Peng, J. (2014). JUMP: a tag-based database search tool for peptide identification with high sensitivity and accuracy. Mol Cell Proteomics 13, 3663–3673. 10.1074/mcp.O114.039586.

113. Peng, J., Elias, J.E., Thoreen, C.C., Licklider, L.J., and Gygi, S.P. (2003). Evaluation of multidimensional chromatography coupled with tandem mass spectrometry (LC/LC-MS/MS) for large-scale protein analysis: the yeast proteome. Journal of proteome research 2, 43–50. 10.1021/pr025556v.

114. Wang, S., Di, Y., Yang, Y., Salovska, B., Li, W., Hu, L., Yin, J., Shao, W., Zhou, D., Cheng, J., et al. (2024). PTMoreR-enabled cross-species PTM mapping and comparative phosphoproteomics across mammals. Cell Rep Methods 4, 100859. 10.1016/j.crmeth.2024.100859.

115. Wang, S., Cai, Y., Cheng, J., Li, W., Liu, Y., and Yang, H. (2019). motifeR: an integrated web software for identification and visualization of protein posttranslational modification motifs. Proteomics 19, 1900245.

116. Hajian-Tilaki, K. (2013). Receiver Operating Characteristic (ROC) Curve Analysis for Medical Diagnostic Test Evaluation. Caspian J Intern Med 4, 627–635.

117. Pedregosa, F., Varoquaux, G., Gramfort, A., Michel, V., Thirion, B., Grisel, O., Blondel, M., Prettenhofer, P., Weiss, R., Dubourg, V., et al. (2011). Scikit-learn: Machine Learning in Python. J. Mach. Learn. Res. 12, 2825–2830.

118. Shannon, P., Markiel, A., Ozier, O., Baliga, N.S., Wang, J.T., Ramage, D., Amin, N., Schwikowski, B., and Ideker, T. (2003). Cytoscape: a software environment for integrated models of biomolecular interaction networks. Genome research 13, 2498–2504. 10.1101/gr.1239303.

119. Tyanova, S., Temu, T., Sinitcyn, P., Carlson, A., Hein, M.Y., Geiger, T., Mann, M., and Cox, J. (2016). The Perseus computational platform for comprehensive analysis of (prote)omics data. Nature Methods 13, 731–740. 10.1038/nmeth.3901.

120. Cox, J., and Mann, M. (2012). 1D and 2D annotation enrichment: a statistical method integrating quantitative proteomics with complementary high-throughput data. BMC bioinformatics 13, S12. 10.1186/1471-2105-13-S16-S12.

121. Zhou, Y., Zhou, B., Pache, L., Chang, M., Khodabakhshi, A.H., Tanaseichuk, O., Benner, C., and Chanda, S.K. (2019). Metascape provides a biologist-oriented resource for the analysis of systems-level datasets. Nature communications 10, 1523. 10.1038/s41467-019-09234-6.

122. Shemesh, N., Jubran, J., Dror, S., Simonovsky, E., Basha, O., Argov, C., Hekselman, I., Abu-Qarn, M., Vinogradov, E., Mauer, O., et al. (2021). The landscape of molecular chaperones across human tissues reveals a layered architecture of core and variable chaperones. Nature communications 12, 2180. 10.1038/s41467-021-22369-9.

123. Perez-Riverol, Y., Bai, J., Bandla, C., Garcia-Seisdedos, D., Hewapathirana, S., Kamatchinathan, S., Kundu, D.J., Prakash, A., Frericks-Zipper, A., Eisenacher, M., et al. (2022). The PRIDE database resources in 2022: a hub for mass spectrometry-based proteomics evidences. Nucleic Acids Res 50, D543–D552. 10.1093/nar/gkab1038.

