## Supplemental Figures for "An Extensive Atlas of Proteome and Phosphoproteome Turnover Across Mouse Tissues and Brain Regions"

**Figure S1**

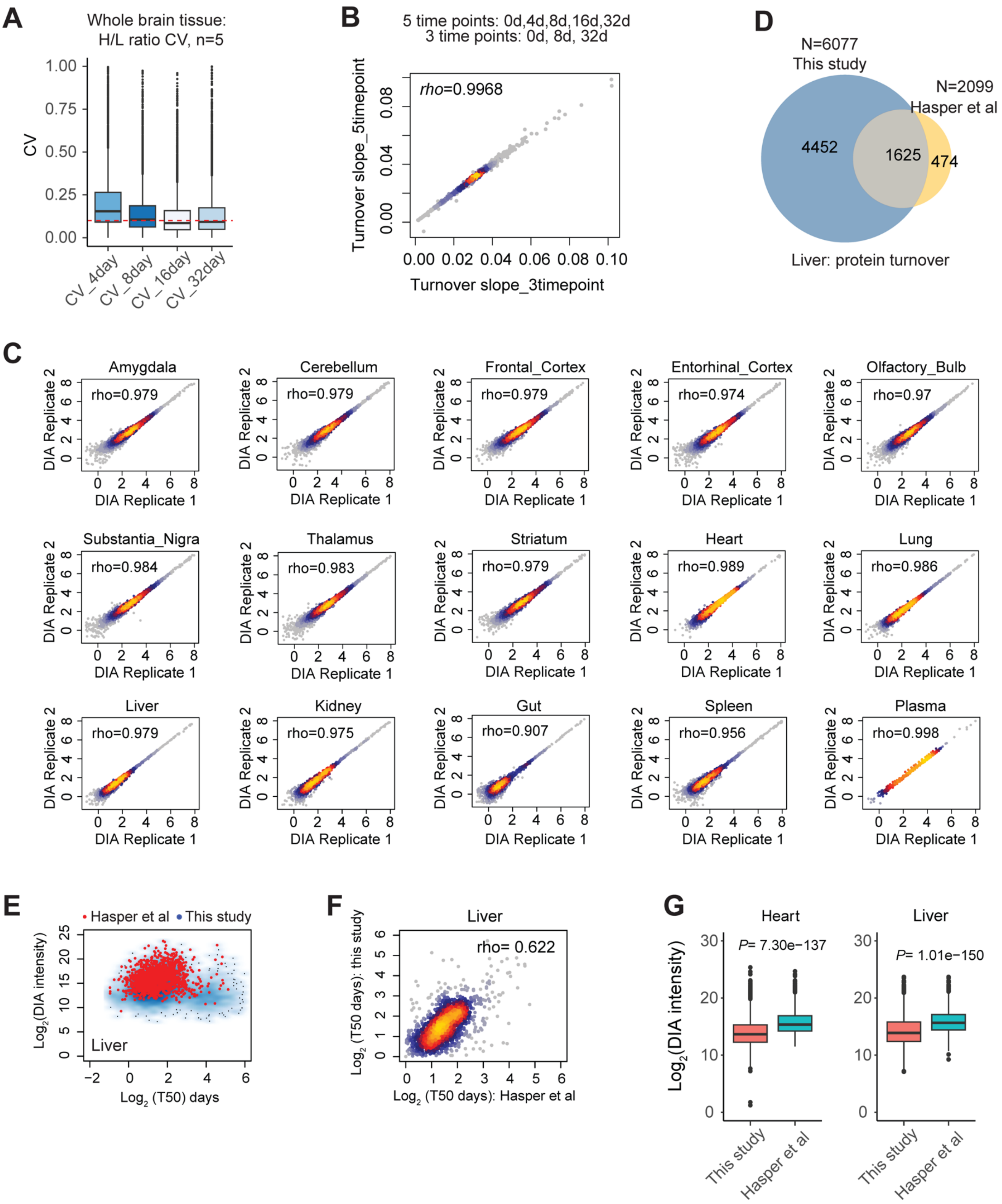
A
Whole brain tissue:
H/L ratio CV, n=5
CV
CV_4day
CV_8day
CV_16day
CV_32day
B
5 time points: 0d,4d,8d,16d,32d
3 time points: 0d, 8d, 32d
rho=0.9968
Turnover slope_5timepoint
Turnover slope_3timepoint
D
N=6077
This study
N=2099
Hasper et al
4452
1625
474
Liver: protein turnover
C
Amygdala
rho=0.979
Cerebellum
rho=0.979
Frontal_Cortex
rho=0.979
Entorhinal_Cortex
rho=0.974
Olfactory_Bulb
rho=0.97
Substantia_Nigra
rho=0.984
Thalamus
rho=0.983
Striatum
rho=0.979
Heart
rho=0.989
Lung
rho=0.986
Liver
rho=0.979
Kidney
rho=0.975
Gut
rho=0.907
Spleen
rho=0.956
Plasma
rho=0.998
DIA Replicate 1
DIA Replicate 2
E
Hasper et al
This study
Log2(DIA intensity)
Liver
Log2 (T50) days
F
Liver
rho= 0.622
Log2 (T50 days): this study
Log2 (T50 days): Hasper et al
G
Heart
P= 7.30e−137
Liver
P= 1.01e−150
Log2(DIA intensity)
This study
Hasper et al

**Figure S1. Establishing a reproducible and in-depth proteome turnover atlas across mouse tissues and brain regions. (Related to Figure 1).**

(A) Coefficient of variation (CV) distribution of proteome-wide Heavy/Light (H/L) ratios in analyzing the mouse whole brain tissue with four SILAC labeling time points of 4, 8, 16, and 32 days (n=5 biological replicates, TMTpro-based quantification).

(B) The quantitative slopes of the log-transformed H/L ratios [i.e., *ln*(H+L)/L] growing over time are highly correlated between the experiments of 5 time points (0d, 4d, 8d, 16d, 32d) and 3 time points (0d, 8d, 32d). The slope essentially determines the protein turnover rates. *Rho*, Spearman correlation coefficient.

(C) Spearman correlation of protein lifetimes between two biological replicates measured by BoxCarmax-DIA.

(D) Venn diagram comparing the turnover coverage of mouse liver proteome between this study and Hasper et al.

(E) Scatterplot displaying the comparison of protein abundance (indicated by DIA-MS intensity in this study) and lifetime for the liver proteome between this study and Hasper et al. (red dots).

(F) Spearman correlation of T50 results between the two studies for the mouse liver proteome.

(G) The boxplot displaying the proteome turnover coverage for the heart and liver proteomes measured by this study and Hasper et al.

**Figure S2**

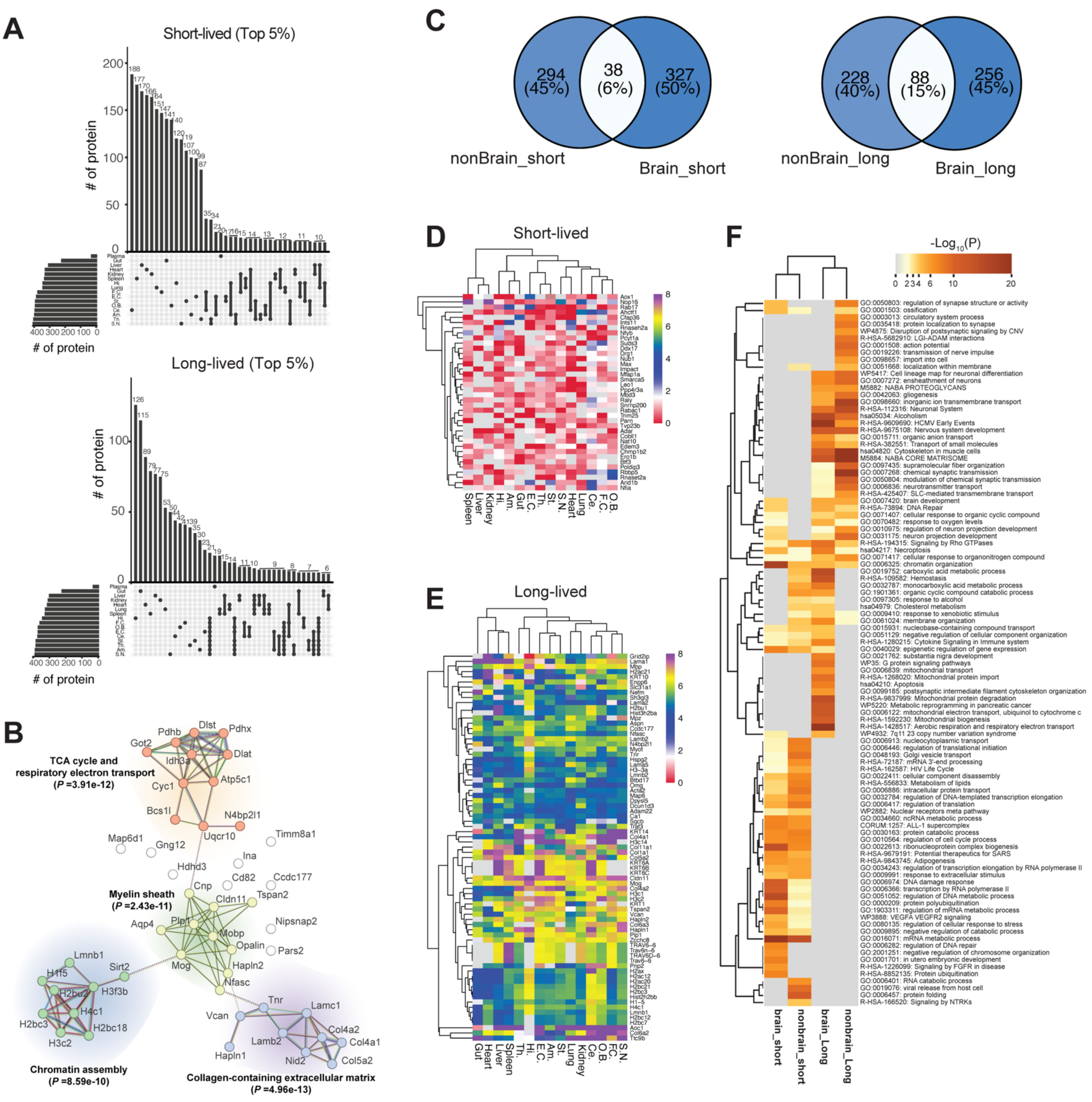

A
Short-lived (Top 5%)
### of protein
### of protein
Long-lived (Top 5%)
### of protein
### of protein
B
TCA cycle and respiratory electron transport (P =3.91e-12)
Myelin sheath (P =2.43e-11)
Chromatin assembly (P =8.59e-10)
Collagen-containing extracellular matrix (P =4.96e-13)
C
294 (45%)
38 (6%)
327 (50%)
nonBrain_short
Brain_short
228 (40%)
88 (15%)
256 (45%)
nonBrain_long
Brain_long
D
Short-lived
E
Long-lived
F
-Log10(P)
brain_short
nonbrain_short
brain_Long
nonbrain_Long

**Figure S2. Short-lived and long-lived proteins and their associated functional processes across tissues. (Related to Figure 2).**

(A) Upset plots of short-lived and long-lived proteins in each tissue. The top 5% shortest- and longest-lived proteins in each tissue were compared between tissues.

(B) STRING Protein-protein interaction network (medium confidence) of 49 proteins that are the top 5% most long-lived in each of the nine brain regions. The Enrichment P values were reported by STRING analysis on the protein clusters.

(C) Venn diagrams comparing short-lived and long-lived protein identities in brain regions and non-brain tissues.

(D) Heatmap of top 5% shortest-lived proteins based on their T50 in brain regions and non-brain tissues. The color bar denotes the $Log_2$ (T50) for each protein.

(E) The same heatmap as (D) for the top 5% longest-lived proteins.

(F) Metascape (https://metascape.org)-derived GO enrichment analysis from short-lived and long-lived proteins in brain regions and non-brain tissues. The Enrichment P values were reported by Metascape analysis.

### Figure S3

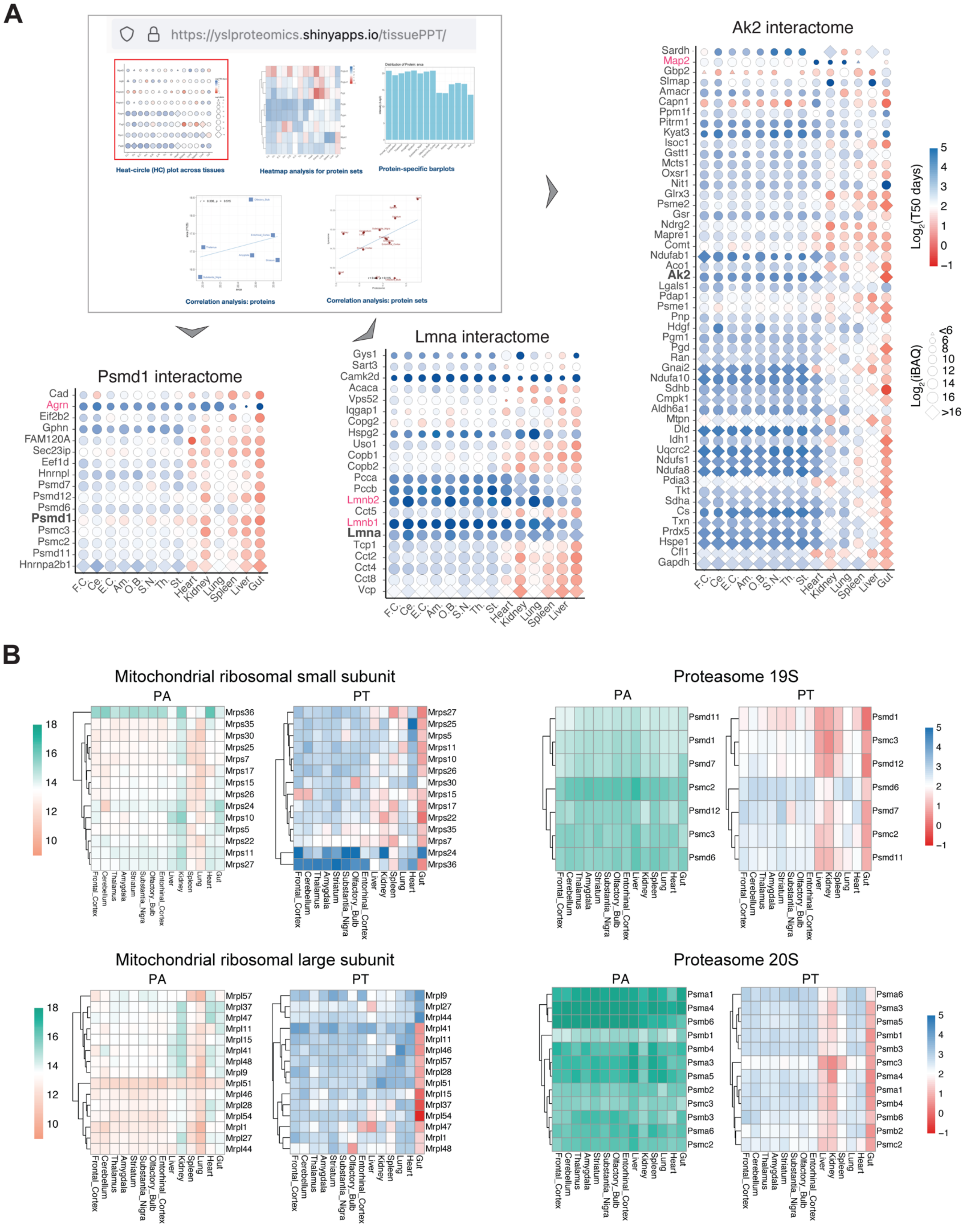
A
https://yslproteomics.shinyapps.io/tissuePPT/
Heat-circle (HC) plot across tissues
Heatmap analysis for protein sets
Protein-specific barplots
Correlation analysis: proteins
Correlation analysis: protein sets
Psmd1 interactome
Lmna interactome
Ak2 interactome
Log2(T50 days)
Log2(iBAQ)
F.C. Ce. E.C. Am. O.B. S.N. Th. St. Heart Kidney Lung Spleen Liver Gut
B
Mitochondrial ribosomal small subunit
Mitochondrial ribosomal large subunit
Proteasome 19S
Proteasome 20S
PA
PT
Frontal_Cortex Cerebellum Thalamus Amygdala Striatum Substantia_Nigra Olfactory_Bulb Entorhinal_Cortex Liver Kidney Spleen Lung Heart Gut

**Figure S3.** ***Tissue-PPT*** **App supports the functional discovery of proteins and protein sets by visualizing both protein abundance (PA) and protein lifetime (PT) and their relationships across tissues. (Related to Figure 2 and 4).**

(A) The Shiny *Tissue-PPT* App (https://yslproteomics.shinyapps.io/tissuePPT/) was established for accessing and analyzing both PA and PT data with examples. The App includes the five major functions: Heat-circle (HC) plot across tissues, Heatmap analysis for protein sets, Protein-specific bar plots, Correlation analysis at the protein level, and Correlation analysis for protein sets. HC plot examples of protein-protein interaction partners of PSMD1, LMNA, and AK2 proteins in across tissues (see Figure 4E).

(B) Examples of heatmaps of both PA and PT generated for proteasome subunits (19S and 20S) and Mitochondrial ribosomal subunits (small/large) across tissues. The scale of PA and PT are kept consist across proteome facilitating direct comparison between individual proteins and protein lists. In this case, PA of proteosome subunits is much higher than that of mitochondrial ribosomal subunits, while both complexes are long-lived. The orange-to-green color bar and red-to-blue color bar denote the small-to-large PA and PT values ($Log_2$-transformed), respectively.

### Figure S4

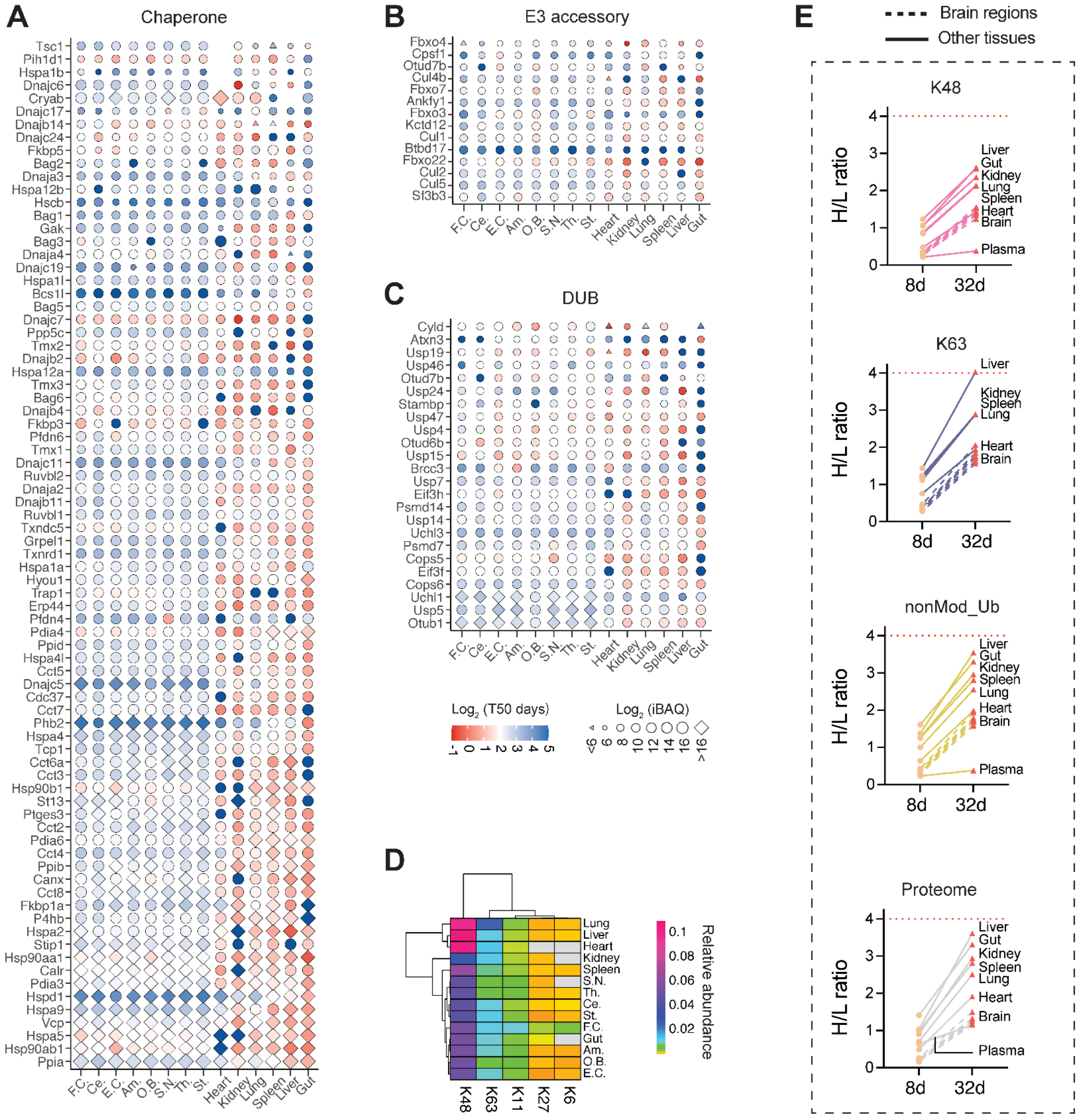
A
Chaperone
Tsc1
Pih1d1
Hspa1b
Dnajc6
Cryab
Dnajc17
Dnajb14
Dnajc24
Fkbp5
Bag2
Dnaja3
Hspa12b
Hscb
Bag1
Gak
Bag3
Dnaja4
Dnajc19
Hspa1l
Bcs1l
Bag5
Dnajc7
Ppp5c
Tmx2
Dnajb2
Hspa12a
Tmx3
Bag6
Dnajb4
Fkbp3
Pfdn6
Tmx1
Dnajc11
Ruvbl2
Dnaja2
Dnajb11
Ruvbl1
Txndc5
Grpel1
Txnrd1
Hspa1a
Hyou1
Trap1
Erp44
Pfdn4
Pdia4
Ppid
Hspa4l
Cct5
Dnajc5
Cdc37
Cct7
Phb2
Hspa4
Tcp1
Cct6a
Cct3
Hsp90b1
St13
Ptges3
Cct2
Pdia6
Cct4
Ppib
Canx
Cct8
Fkbp1a
P4hb
Hspa2
Stip1
Hsp90aa1
Calr
Pdia3
Hspd1
Hspa9
Vcp
Hspa5
Hsp90ab1
Ppia
F.C.
Ce.
E.C.
Am.
O.B.
S.N.
Th.
St.
Heart
Kidney
Lung
Spleen
Liver
Gut
B
E3 accessory
Fbxo4
Cpsf1
Otud7b
Cul4b
Fbxo7
Ankfy1
Fbxo3
Kctd12
Cul1
Btbd17
Fbxo22
Cul2
Cul5
Sf3b3
F.C.
Ce.
E.C.
Am.
O.B.
S.N.
Th.
St.
Heart
Kidney
Lung
Spleen
Liver
Gut
C
DUB
Cyld
Atxn3
Usp19
Usp46
Otud7b
Usp24
Stambp
Usp47
Usp4
Otud6b
Usp15
Brcc3
Usp7
Eif3h
Psmd14
Usp14
Uchl3
Psmd7
Cops5
Eif3f
Cops6
Uchl1
Usp5
Otub1
F.C.
Ce.
E.C.
Am.
O.B.
S.N.
Th.
St.
Heart
Kidney
Lung
Spleen
Liver
Gut
Log2 (T50 days)
-1 0 1 2 3 4 5
Log2 (iBAQ)
<6 6 8 10 12 14 16 >16
D
Lung
Liver
Heart
Kidney
Spleen
S.N.
Th.
Ce.
St.
F.C.
Gut
Am.
O.B.
E.C.
K48
K63
K11
K27
K6
0.1
0.08
0.06
0.04
0.02
Relative abundance
E
Brain regions
Other tissues
K48
H/L ratio
0 1 2 3 4
8d
32d
Liver
Gut
Kidney
Lung
Spleen
Heart
Brain
Plasma
K63
H/L ratio
0 1 2 3 4
8d
32d
Liver
Kidney
Spleen
Lung
Heart
Brain
nonMod_Ub
H/L ratio
0 1 2 3 4
8d
32d
Liver
Gut
Kidney
Spleen
Lung
Heart
Brain
Plasma
Proteome
H/L ratio
0 1 2 3 4
8d
32d
Liver
Gut
Kidney
Spleen
Lung
Heart
Brain
Plasma

**Figure S4. Tissue-specific degradation associated proteins, distinctive distribution patterns of linkage-specific ubiquitination, and their renewal rates. (Related to Figure 3).**

(A-C) Heat-circle (HC) plots of protein folding chaperones, E3 accessory proteins, and deubiquitinating enzymes (DUBs) across tissues.

(D) Hierarchical clustering heatmap of ubiquitin (Ub) linkages K63, K48, K27, K11, and K6 across tissues. The color bar indicates the relative abundance of Ub chains, normalized using unmodified ubiquitin peptides (see Methods).

(E) Heavy-to-light (H/L) ratios during pSILAC labeling for K48- and K63-linked Ubs, the unmodified ubiquitin peptide TLSDYNIQK, and the averaged proteome (from top to bottom). The dashed lines indicate brain regions, while the solid lines denote other main tissues.

### Figure S5

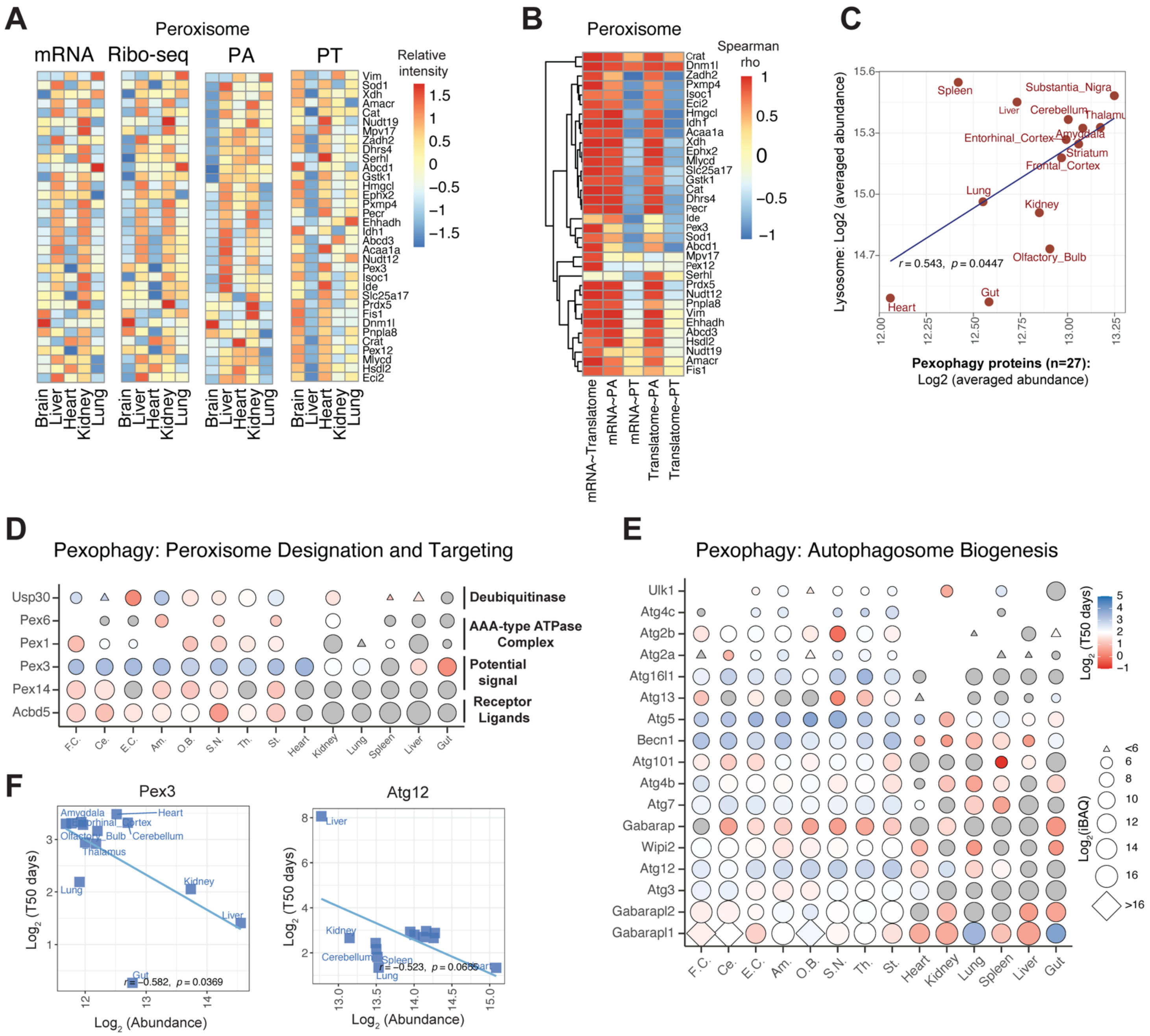

A
Peroxisome
mRNA
Ribo-seq
PA
PT
Relative intensity
Vim
Sod1
Xdh
Amacr
Cat
Nudt19
Mpv17
Zadh2
Dhrs4
Serhl
Abcd1
Gstk1
Hmgcl
Ephx2
Pxmp4
Pecr
Ehhadh
Idh1
Abcd3
Acaa1a
Nudt12
Pex3
Isoc1
Ide
Slc25a17
Prdx5
Fis1
Dnm1l
Pnpla8
Crat
Pex12
Mlycd
Hsdl2
Eci2
Brain
Liver
Heart
Kidney
Lung
B
Peroxisome
Spearman rho
mRNA~Translatome
mRNA~PA
mRNA~PT
Translatome~PA
Translatome~PT
C
Lysosome: Log2 (averaged abundance)
Pexophagy proteins (n=27): Log2 (averaged abundance)
Spleen
Liver
Substantia_Nigra
Cerebellum
Thalamus
Entorhinal_Cortex
Amygdala
Striatum
Frontal_Cortex
Lung
Kidney
Olfactory_Bulb
Gut
Heart
r = 0.543, p = 0.0447
D
Pexophagy: Peroxisome Designation and Targeting
Usp30
Pex6
Pex1
Pex3
Pex14
Acbd5
Deubiquitinase
AAA-type ATPase Complex
Potential signal
Receptor Ligands
F.C.
Ce.
E.C.
Am.
O.B.
S.N.
Th.
St.
Heart
Kidney
Lung
Spleen
Liver
Gut
E
Pexophagy: Autophagosome Biogenesis
Ulk1
Atg4c
Atg2b
Atg2a
Atg16l1
Atg13
Atg5
Becn1
Atg101
Atg4b
Atg7
Gabarap
Wipi2
Atg12
Atg3
Gabarapl2
Gabarapl1
Log2 (T50 days)
Log2(iBAQ)
<6
>16
F
Pex3
Log2 (T50 days)
Log2 (Abundance)
Amygdala
Heart
Entorhinal_Cortex
Olfactory_Bulb
Cerebellum
Thalamus
Lung
Kidney
Liver
Gut
r = −0.582, p = 0.0369
Atg12
Liver
Kidney
Cerebellum
Spleen
Lung
Heart
r = −0.523, p = 0.0655

**Figure S5. Multi-omic analysis of peroxisome and pexophagy proteins. (Related to Figure 5).**

(A) Heatmap of individual peroxisome proteins among five tissues using the quantitative values from mRNA, translatome, PA, and PT layers. The brain results were determined by averaging all brain regions. The heatmap was created by the R pheatmap package with the row sale. Scale bar: red color is higher value per layer; blue color is lower value.

(B) Heatmap visualizing the cross-tissue Spearman correlation of each peroxisome protein between two omics layers across tissues. The blue-to-red scale bar denotes the increasing Spearman correlation. The translatome data was measured by ribosome sequencing (Ribo-seq).

(C) The negative correlation between the abundances of pexophagy-associated proteins (n=27) and the lysosome protein levels (see Figure 3b). Plots were generated by the "Correlation analysis: protein sets" function in *Tissue-PPT*. Pearson correlation (r) and the significance of correlation (P value) are shown.

(D) HC plots of proteins identified for pexophagy-associated proteins involved in peroxisome designation and targeting. Plots were generated by the "Heat-circle (HC) plot across tissues" function in *Tissue-PPT*.

(E) HC plots of proteins identified for pexophagy-associated proteins involved in autophagosome biogenesis.

(F) Correlation between Log2 (Protein abundance) and Log2 ($T_{50}$) for protein examples of Pex3 and Atg12 involved in pexophagy. Plots were generated by the "Correlation analysis: proteins" function in *Tissue-PPT*.

**Figure S6**

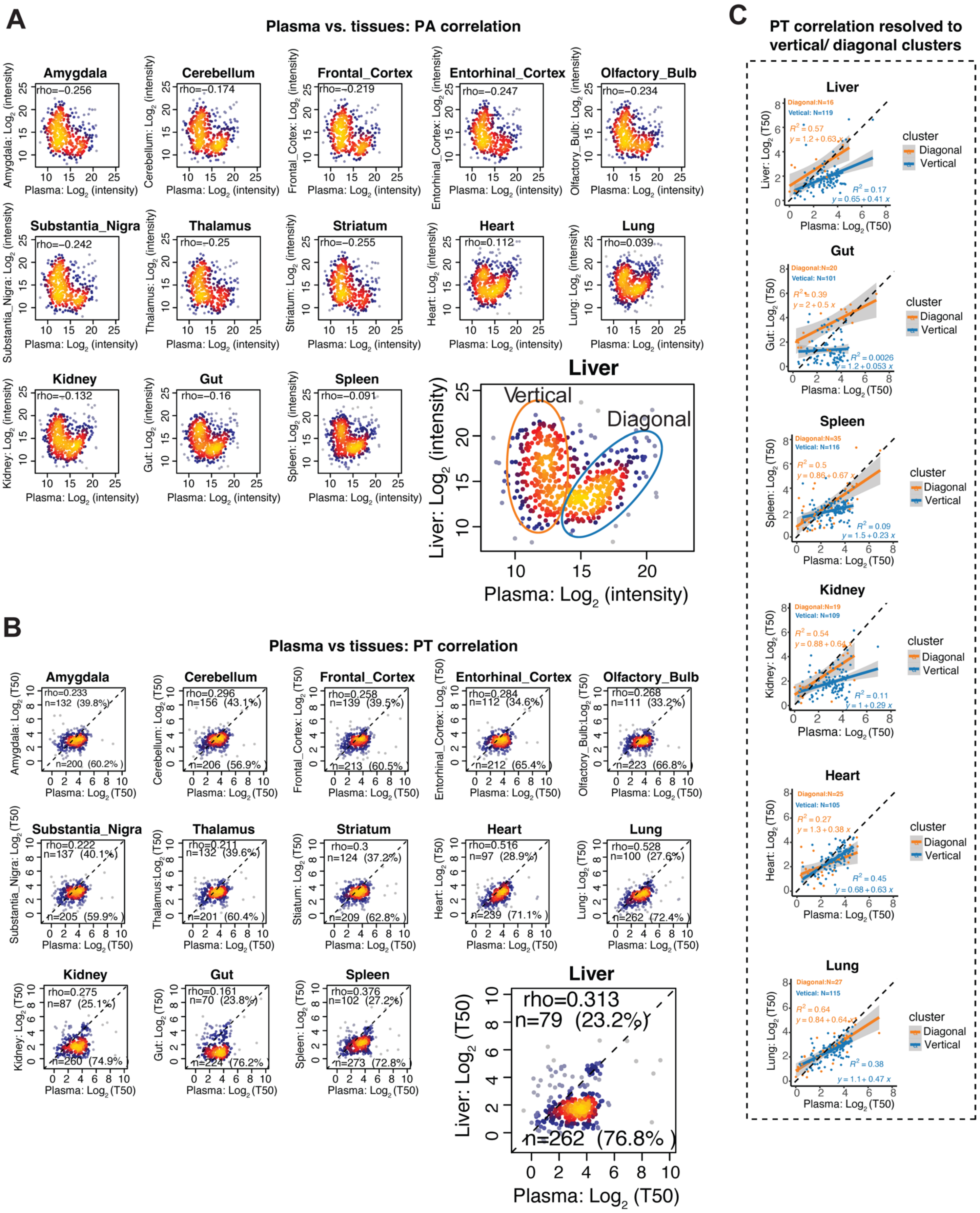
A
Plasma vs. tissues: PA correlation
Amygdala
rho=−0.256
Cerebellum
rho=−0.174
Frontal_Cortex
rho=−0.219
Entorhinal_Cortex
rho=−0.247
Olfactory_Bulb
rho=−0.234
Substantia_Nigra
rho=−0.242
Thalamus
rho=−0.25
Striatum
rho=−0.255
Heart
rho=0.112
Lung
rho=0.039
Kidney
rho=−0.132
Gut
rho=−0.16
Spleen
rho=−0.091
Liver
Vertical
Diagonal
Plasma: Log2 (intensity)
Liver: Log2 (intensity)
B
Plasma vs tissues: PT correlation
Amygdala
rho=0.233
n=132 (39.8%)
n=200 (60.2%)
Cerebellum
rho=0.296
n=156 (43.1%)
n=206 (56.9%)
Frontal_Cortex
rho=0.258
n=139 (39.5%)
n=213 (60.5%)
Entorhinal_Cortex
rho=0.284
n=112 (34.6%)
n=212 (65.4%)
Olfactory_Bulb
rho=0.268
n=111 (33.2%)
n=223 (66.8%)
Substantia_Nigra
rho=0.222
n=137 (40.1%)
n=205 (59.9%)
Thalamus
rho=0.211
n=132 (39.6%)
n=201 (60.4%)
Striatum
rho=0.3
n=124 (37.2%)
n=209 (62.8%)
Heart
rho=0.516
n=97 (28.9%)
n=239 (71.1%)
Lung
rho=0.528
n=100 (27.6%)
n=262 (72.4%)
Kidney
rho=0.275
n=87 (25.1%)
n=260 (74.9%)
Gut
rho=0.161
n=70 (23.8%)
n=224 (76.2%)
Spleen
rho=0.376
n=102 (27.2%)
n=273 (72.8%)
Liver
rho=0.313
n=79 (23.2%)
n=262 (76.8%)
Plasma: Log2 (T50)
Liver: Log2 (T50)
C
PT correlation resolved to vertical/ diagonal clusters
Liver
Diagonal:N=16
Vetical: N=119
R2 = 0.57
y = 1.2 + 0.63 x
R2 = 0.17
y = 0.65 + 0.41 x
Gut
Diagonal:N=20
Vetical: N=101
R2 = 0.39
y = 2 + 0.5 x
R2 = 0.0026
y = 1.2 + 0.053 x
Spleen
Diagonal:N=35
Vetical: N=116
R2 = 0.5
y = 0.86 + 0.67 x
R2 = 0.09
y = 1.5 + 0.23 x
Kidney
Diagonal:N=19
Vetical: N=109
R2 = 0.54
y = 0.88 + 0.64 x
R2 = 0.11
y = 1 + 0.29 x
Heart
Diagonal:N=25
Vetical: N=105
R2 = 0.27
y = 1.3 + 0.38 x
R2 = 0.45
y = 0.68 + 0.63 x
Lung
Diagonal:N=27
Vetical: N=115
R2 = 0.64
y = 0.84 + 0.64 x
R2 = 0.38
y = 1.1 + 0.47 x
cluster
Diagonal
Vertical
Plasma: Log2 (T50)

**Figure S6. A bimodal distribution of both PAs and PTs in the plasma proteome highlights the divergent tissue origins of blood plasma proteins.**

(A) The correlation of $Log_2$ (DIA-intensity) for the same proteins identified and measured in plasma and tissues. The scatter plots derive two clusters of proteins, namely "diagonal" and "vertical" ones, which were observed for most of tissues, with solid tissues like liver being most significant. For each of the plasma proteins, its X axis denotes PA in plasma while y axis represents PA in other tissues for the same protein. *Rho*, Spearman correlation coefficient.

(B) The correlation of $Log_2$ ($T_{50}$) for the same proteins identified and measured in plasma and tissues. For each of the plasma proteins, its X axis denotes the PT in plasma while y axis represents PT in other tissues for the same protein. The dashed line indicates the diagonal line of y=x (i.e., identical PT between tissue and plasma), which divides the proteins into two groups. The number and percentage of proteins in each of these two groups are shown, suggesting the PTs of most plasma proteins are generally higher than in plasma than other tissues (i.e., the majority of the proteins are below the diagonal line). For example, 76.8% of the liver proteins overlapping to the plasma proteome have a longer lifetime in blood plasma.

(C) PT linear regression between plasma and different tissues, separately correlated for plasma proteins classified as "diagonal" and "vertical" clusters. The "diagonal" and "vertical" clusters were defined by the intensity correlation of the plasma proteins, as shown in (A). A "diagonal" protein was defined by its intensity $Log_2$ (DIA-intensity) >14 in the plasma sample, whereas other proteins were classified into the "vertical" cluster. Orange and blue lines depict the fitted PT linear regression curve for "diagonal" and “vertical” clusters in each of the tissue, respectively. The brain regions were not shown due to the small number of plasma proteins for the “diagonal” cluster profiled with PT in the brain. The intriguing pattern that the orange lines are closer to the y=x line for most tissues suggests the plasma proteins classified as "diagonal" group tend to have the same PT between plasma and tissues, possibly due to shared protein origination, as compared to the “vertical” group which has longer PT in plasma.

**Figure S7**

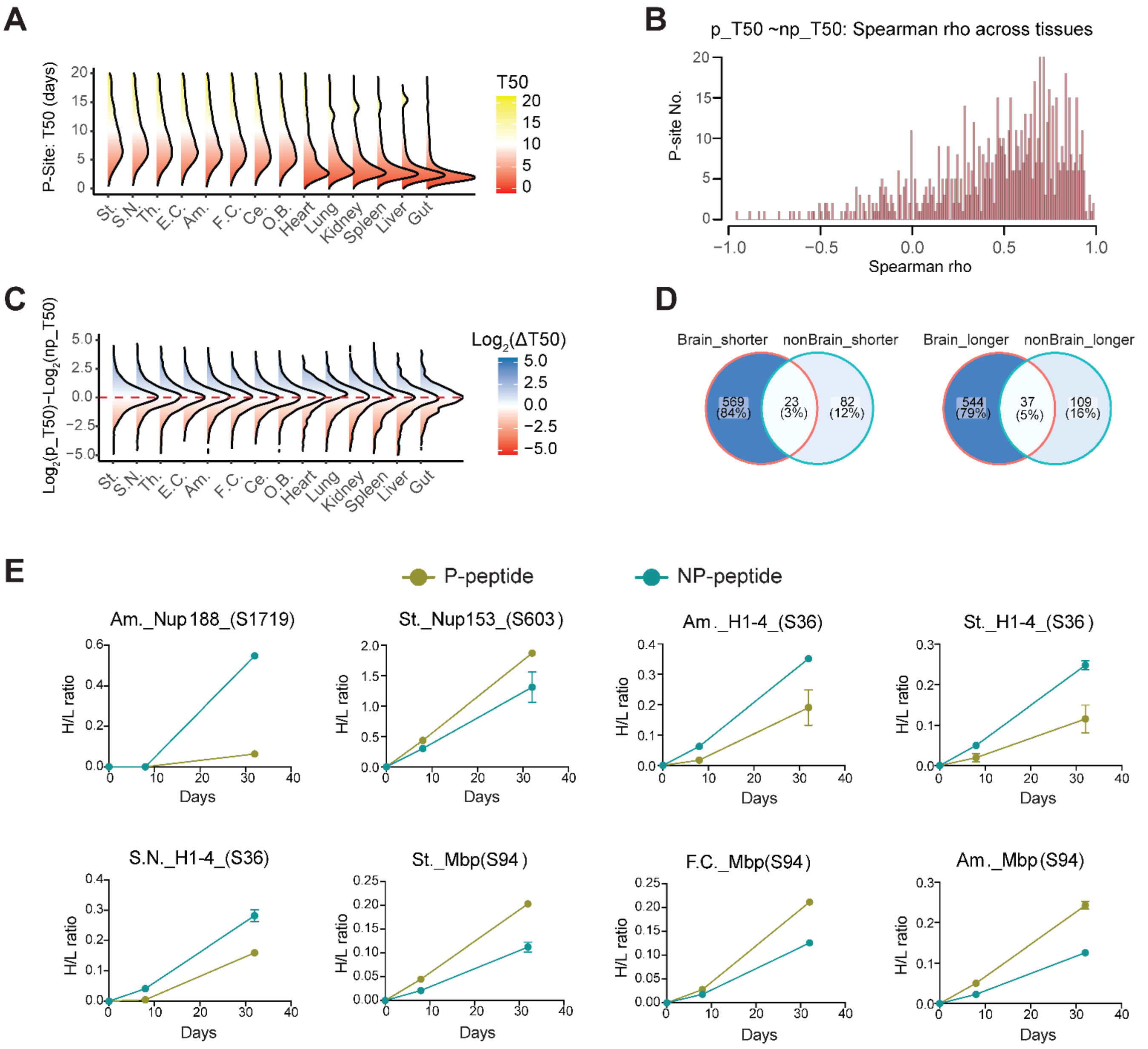


**Figure S7. Phosphoproteome turnover measurement across mouse tissues reveals the different T50 of the phosphorylated peptides and their non-phosphorylated counterparts. (Related to Figure 6 and 7).**

(A) Density plot of the quantified T50 values of phosphorylation sites (P-sites) across mouse tissues based on all phosphopeptides measured.

(B) Distribution of Spearman correlation between the $T_{50}$ of a phosphorylated (p) peptide and the $T_{50}$ of its non-phosphorylated (np) peptide counterpart across tissues and regions for all P-sites. The p and np peptides share the same amino acid sequence and only differ in the phosphorylation modification.

(C) Density plot of Delta$T_{50}$ ($\Delta T_{50}$) that represents the difference between the $T_{50}$ of p peptide to the $T_{50}$ of its corresponding np peptide across tissues.

(D) Venn diagram comparing the numbers of P-sites showing destabilization (T50 of p peptide is shorter than T50 of np peptide) or stabilization effects (T50 of p peptide is shorter than T50 of np peptide) in brain and non-brain areas. The significance is inferred by P-value <0.05 (Student's t-test) and fold change >1.5 (in brain) and >1.2 (in non-brain tissues).

(E) Heavy/light (H/L) ratio curves during the labeling process for a p peptide (green) and its np peptide counterpart (blue) of the same sequence for examples of long-lived proteins.

**Figure S8**

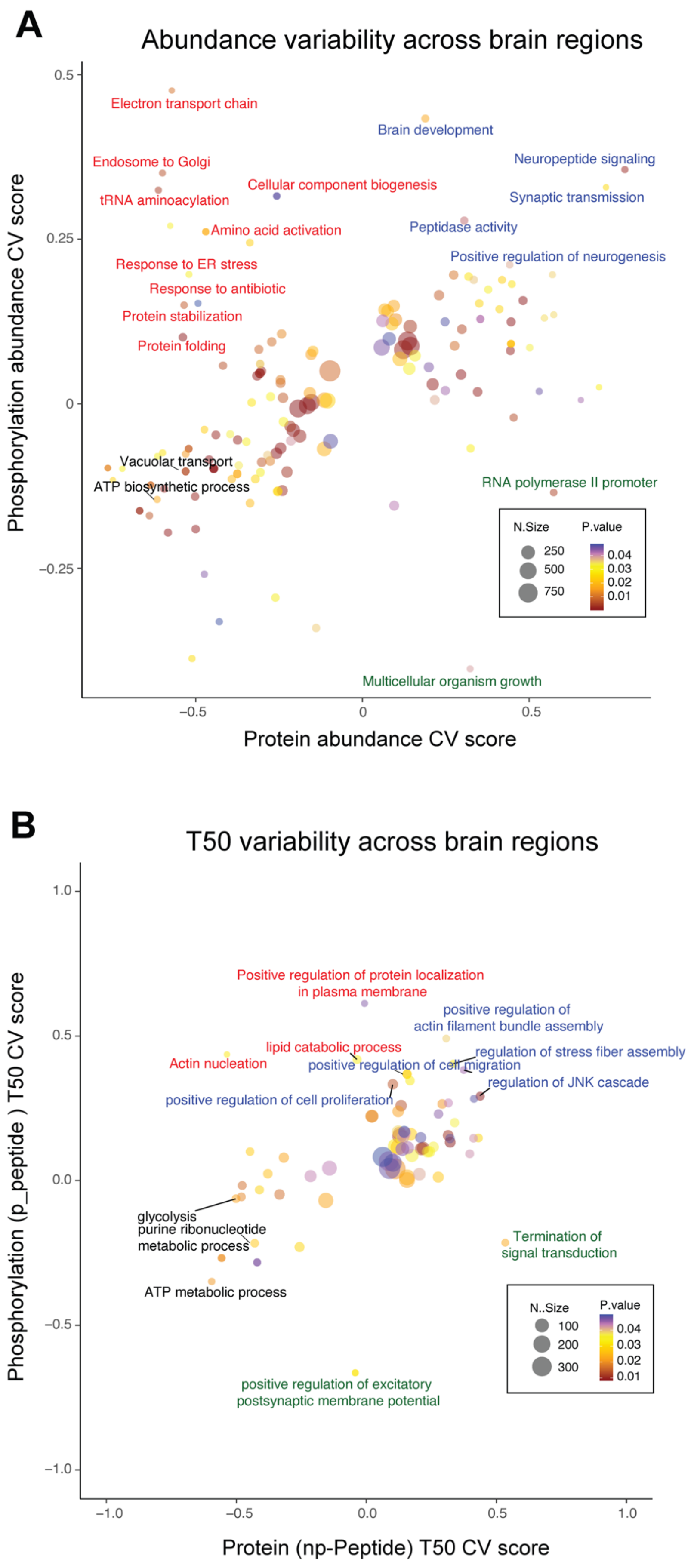

A
Abundance variability across brain regions
Phosphorylation abundance CV score
Protein abundance CV score
0.5
0.25
0
−0.25
−0.5
0
0.5
Electron transport chain
Endosome to Golgi
tRNA aminoacylation
Cellular component biogenesis
Amino acid activation
Response to ER stress
Response to antibiotic
Protein stabilization
Protein folding
Brain development
Neuropeptide signaling
Synaptic transmission
Peptidase activity
Positive regulation of neurogenesis
Vacuolar transport
ATP biosynthetic process
RNA polymerase II promoter
Multicellular organism growth
N.Size
250
500
750
P.value
0.04
0.03
0.02
0.01
B
T50 variability across brain regions
Phosphorylation (p_peptide ) T50 CV score
Protein (np-Peptide) T50 CV score
1.0
0.5
0.0
−0.5
−1.0
−1.0
−0.5
0.0
0.5
1.0
Positive regulation of protein localization
in plasma membrane
positive regulation of
actin filament bundle assembly
lipid catabolic process
Actin nucleation
regulation of stress fiber assembly
positive regulation of cell migration
regulation of JNK cascade
positive regulation of cell proliferation
glycolysis
purine ribonucleotide
metabolic process
ATP metabolic process
Termination of
signal transduction
positive regulation of excitatory
postsynaptic membrane potential
N..Size
100
200
300
P.value
0.04
0.03
0.02
0.01

**Figure S8. Two dimensional (2D) functional enrichment analysis comparing proteome and phosphoproteome abundance and turnover variability across brain regions. (Related to Figure 6).**

(A) Bubble plot of abundance variability between proteome and phosphoproteome across brain regions. CV was calculated from phosphorylated protein abundance (y-axis, an average of all phosphosites) or protein abundance (x-axis) across all brain regions. 2D enrichment was performed using Perseus software and the bubble plot was visualized using ggplot2. Enriched GO terms were filtered with p-value $<0.05$ and number of proteins $>5$. The cycle size denotes the number of proteins in each GO term. The color scale represents the enriched GO p-value.

(B) Bubble plot of lifetime variability between proteome and phosphoproteome across brain regions. CV was calculated from phosphorylated protein lifetime (y-axis, an average of all phosphosites) and protein lifetime (x-axis) across all brain regions. The cycle size denotes the number of proteins in each GO term. The color scale represents the enriched GO p-value.
